# Tools for connectomic reconstruction and analysis of a female *Drosophila* ventral nerve cord

**DOI:** 10.1101/2022.12.15.520299

**Authors:** Anthony Azevedo, Ellen Lesser, Jasper S. Phelps, Brandon Mark, Leila Elabbady, Sumiya Kuroda, Anne Sustar, Anthony Moussa, Avinash Kandelwal, Chris J. Dallmann, Sweta Agrawal, Su-Yee J. Lee, Brandon Pratt, Andrew Cook, Kyobi Skutt-Kakaria, Stephan Gerhard, Ran Lu, Nico Kemnitz, Kisuk Lee, Akhilesh Halageri, Manuel Castro, Dodam Ih, Jay Gager, Marwan Tammam, Sven Dorkenwald, Forrest Collman, Casey Schneider-Mizell, Derrick Brittain, Chris S. Jordan, Michael Dickinson, Alexandra Pacureanu, H. Sebastian Seung, Thomas Macrina, Wei-Chung Allen Lee, John C. Tuthill

## Abstract

Like the vertebrate spinal cord, the insect ventral nerve cord (VNC) mediates limb sensation and motor control. Here, we apply automated tools for electron microscopy volume alignment, neuron segmentation, and synapse prediction toward creating a connectome of an adult female *Drosophila* VNC. To interpret a connectome, it is crucial to know its relationship with the rest of the body. We therefore mapped the muscle targets of leg and wing motor neurons in the connectome by comparing their morphology to genetic driver lines, dye fills, and X-ray nano-tomography of the fly leg and wing. Knowing the outputs of the connectome allowed us to identify neural circuits that coordinate the wings and legs during escape takeoff. We provide the reconstruction of VNC circuits and motor neuron atlas, along with tools for programmatic and interactive access, as community resources to support experimental and theoretical studies of how the fly nervous system controls behavior.

## Introduction

A principal function of the nervous system is to move the body. A longstanding question in neuroscience is how neural circuits control adaptive motor behaviors, such as limbed locomotion. In vertebrate animals, much effort has been dedicated to understanding cortical and subcortical circuits that plan movements and issue descending commands (Arber and Costa, 2022; Inagaki et al., 2022; Leiras et al., 2022). There is also an extensive body of literature on the physiology and function of motor neurons (MNs) that directly control muscle activity (Binder et al., 2020; Kernell, 2006). In comparison, less is known about the function of intermediate circuits of the spinal cord, where proprioceptive feedback signals are integrated with descending motor commands to coordinate patterns of muscle activity (Goulding, 2009; Leiras et al., 2022; Ruder and Arber, 2019). Understanding the architecture and activity patterns of spinal circuits has been challenging due to their experimental inaccessibility, cell-type diversity, and the large number of spinal interneurons and MNs.

The adult fruit fly, *Drosophila melanogaster*, is a tractable model system to investigate neural circuits for adaptive motor control (Tuthill and Wilson, 2016). Leg and wing motor circuits in the fly are contained within the ventral nerve cord (VNC), which functions like the vertebrate spinal cord to sense and move the limbs (Court et al., 2020). Each of the fly’s six legs is controlled by 60-70 uniquely identifiable MNs (Baek and Mann, 2009; Brierley et al., 2012; Phelps et al., 2021). For comparison, a single calf muscle in the cat is innervated by ∼600 MNs (Burke, 2011).

Despite differences in scale, limb motor control systems of diverse animal species possess key similarities (Manuel et al., 2019). For example, recent work has shown that *Drosophila* leg MNs are hierarchically recruited according to the size principle (Azevedo et al., 2020), a model formulated 60 years ago to explain relationships between MN anatomy and muscle force production in the cat leg (Henneman et al., 1965a, 1965b; Henneman and Olson, 1965; Mcphedran et al., 1965; Wuerker et al., 1965). Evidence that the size principle contributes to hierarchical MN recruitment has been found in many species, from crayfish (Hill and Cattaert, 2008) to zebrafish (Ampatzis et al., 2013) to humans (Heckman and Enoka, 2012). Thus, studying the compact motor circuits of *Drosophila* has the potential to identify general circuit mechanisms with relevance for understanding motor control in other limbed animals.

Flies have two distinct modes of locomotion: walking and flying. Unlike the leg, the *Drosophila* wing has anatomically, physiologically, and functionally distinct muscles for power and steering (Dickinson and Tu, 1997). The wing’s power and steering muscles attach to the thorax and wing hinge, respectively. They are controlled by just 29 MNs on each side; the dendritic anatomy and muscle innervation patterns of most of the wing MNs have been previously described from light microscopy (Ehrhardt et al., 2023; O’Sullivan et al., 2018). As with the leg system, the number of wing MNs is remarkably small compared with their vertebrate counterparts — a single pectoralis muscle of a hummingbird is innervated by ∼2000 MNs (Donovan et al., 2013).

Here, we report two advances for understanding how the fly’s central nervous system controls the body. First, we apply machine learning tools for automated neuron segmentation and synapse prediction to an EM volume of the *Drosophila* **F**emale **A**dult **N**erve **C**ord (FANC) (Phelps et al., 2021). We describe the application of new software tools for cell-type annotation, querying connectivity, and identification of genetic driver lines based on reconstructed neuronal morphology. The FANC connectome is being actively proofread and analyzed by a consortium of 30 labs. We now provide the FANC segmentation and interactive tools as resources to the broader neuroscience community. This work complements a parallel project to reconstruct the connectome of an adult male VNC (Cheong et al., 2024; Marin et al., 2023; Takemura et al., 2023), as well as two existing connectomes for the adult female *Drosophila* brain (Dorkenwald et al., 2023a; Xu et al., 2020).

We also address a fundamental limitation of existing EM datasets of the adult *Drosophila* brain and VNC, namely that they contain only the central nervous system. The fly’s body and peripheral nervous system, including muscles, sensory neuron somas and dendrites, and MN axons, are typically dissected away during sample preparation. The inability to reconstruct the peripheral inputs and outputs of the connectome poses a particular challenge for investigating how central circuits mediate motor control of the body. There does not currently exist a comprehensive projection map of MN to muscle connectivity in any limbed animal. Here, we determined which MNs in the FANC connectome control which muscles. We assembled a comprehensive atlas of MNs by combining EM reconstruction, sparse genetic driver lines (Meissner et al., 2023), and X-ray nanotomography of the fly’s front leg and wing (Kuan et al., 2020). Because the morphology of MNs is stereotyped from fly to fly, this projection map can be used to identify MNs in existing and future connectomes (Cheong et al., 2024). Together, the reconstruction of a female VNC and the identification of the peripheral targets of MNs provide a foundation for investigating how sensorimotor circuits move the body and integrate descending signals from the brain to control behavior.

## Results

### Automated reconstruction and tools for analysis of an EM volume of a female fly VNC

We previously imaged an electron microscopy volume of a *Drosophila* Female Adult Nerve Cord (FANC, pronounced “fancy”) and manually reconstructed a small number of VNC neurons and synapses (Phelps et al., 2021). To accelerate efforts to map the VNC connectome, here we applied deep-learning based approaches to automatically reconstruct neurons and synaptic connections throughout FANC. Proper alignment of serial sections is crucial for automated segmentation and data analysis, so we first refined the alignment of FANC’s image data using self-supervised convolutional neural networks (Popovych et al., 2022). We then used convolutional neural networks (CNNs) to segment the dataset into neurons and fragments of neurons (Macrina et al., 2021) and to predict synapse locations (**Figure 1c**), as well as pre- and post-synaptic partners (Buhmann et al., 2021). To identify all cells intrinsic to the VNC, we detected nuclei using another CNN (**Figure 1e-f** and Methods) (Lee et al., 2017; Mu et al., 2021), which identified 17,076 putative cells. Through manual inspection, we classified 14,621 of these nuclei as neuronal (85.6%). The remaining objects were glia nuclei (11.9%) or non-nucleus objects (2.5%, **Extended Data Figure 1a-c**). The CNN, trained on neuronal nuclei, likely did not detect most glia nuclei.

**Figure 1.**
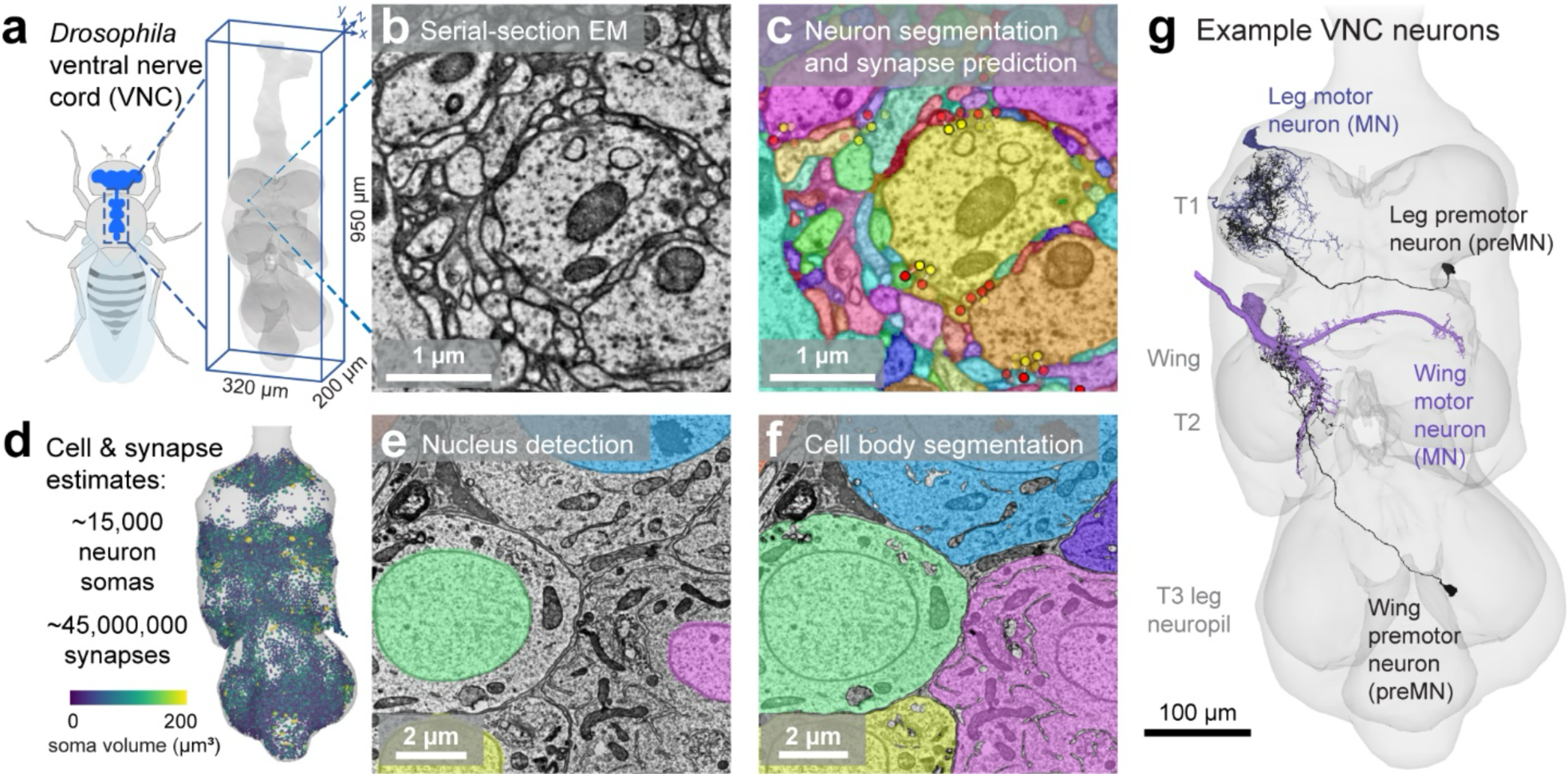
Automated tools for connectomic reconstruction of neural circuits in the *Drosophila* VNC. **a,** We aligned, segmented, and analyzed a serial-section electron microscopy dataset of a *Drosophila* <u>F</u>emale <u>A</u>dult ventral <u>N</u>erve <u>C</u>ord (FANC). **b,** Example section of raw EM image data from the FANC dataset. **c,** We automatically segmented neurons using convolutional neural nets and mean affinity agglomeration with semantic and size constraints. Each segmented cell is shaded with a different color. We applied automated methods for synapse identification across the entire FANC dataset. Example presynaptic sites are labeled with yellow dots and postsynaptic sites are labeled red. **d,** We counted the total number of neurons in FANC by automatically detecting cell nuclei **e** and segmenting cell bodies **f**. **g,** We visualized and proofread segmented cells in Neuroglancer, a WebGL-based viewer for volumetric data.

The automated cell segmentation was ingested into a ChunkedGraph (Dorkenwald et al., 2022), which allowed human experts to proofread errors in the reconstructions, such as attaching cell bodies and splitting neuronal processes from glia, through the web interface Neuroglancer (Maitin-Shepard et al., 2021). We imported synapse and cell body predictions into the Connectome Annotation Versioning Engine (CAVE), so that the associated cell segmentation for each annotation would be dynamically updated during proofreading (Dorkenwald et al., 2023b). This system allowed us to query up-to-date connectivity and associated metadata, such as cell-type annotations. The suite of tools available for analysis of the FANC dataset are described in **Extended Data Figure 2a-e**, including a web platform (https://braincircuits.io) for performing color-depth MIP searches (Otsuna et al., 2018) to find genetic driver lines that label any neuron within the FANC dataset. Following the example of FlyWire (Dorkenwald et al., 2022), we organized a community currently comprising over 200 users from over 30 labs who use these tools to proofread the FANC cell segmentation and analyze their circuits of interest. Representative examples of proofread motor and premotor neurons are shown in **Figure 1g**. (Links within the manuscript lead to 3D visualizations of reconstructed neurons).

To assess the quality of the FANC automated segmentation and synapse prediction, we focused on leg MNs (**Figure 1a**). As in other animals (Manuel et al., 2019), MNs are among the largest cells in the *Drosophila* nervous system (**Extended Data Figure 1a-c**). The largest leg MN in the fly, the fast tibia flexor, has nearly 1 cm of total dendritic length (a typical *Drosophila* is ∼3 mm in length). Manual tracing and synapse annotation of the fast tibia flexor MN took an expert tracer approximately 200 hours. For comparison, proofreading the automated reconstruction of this same MN to correct errors in the automated segmentation took only 2 hours. Overlaying the manual and automated reconstructions for individual MNs (**Figure 2b-c**) illustrates that the automated methods, followed by proofreading, effectively captured the major dendritic branches and >80% of the total cable (**Figure 2d**).

**Figure 2.**
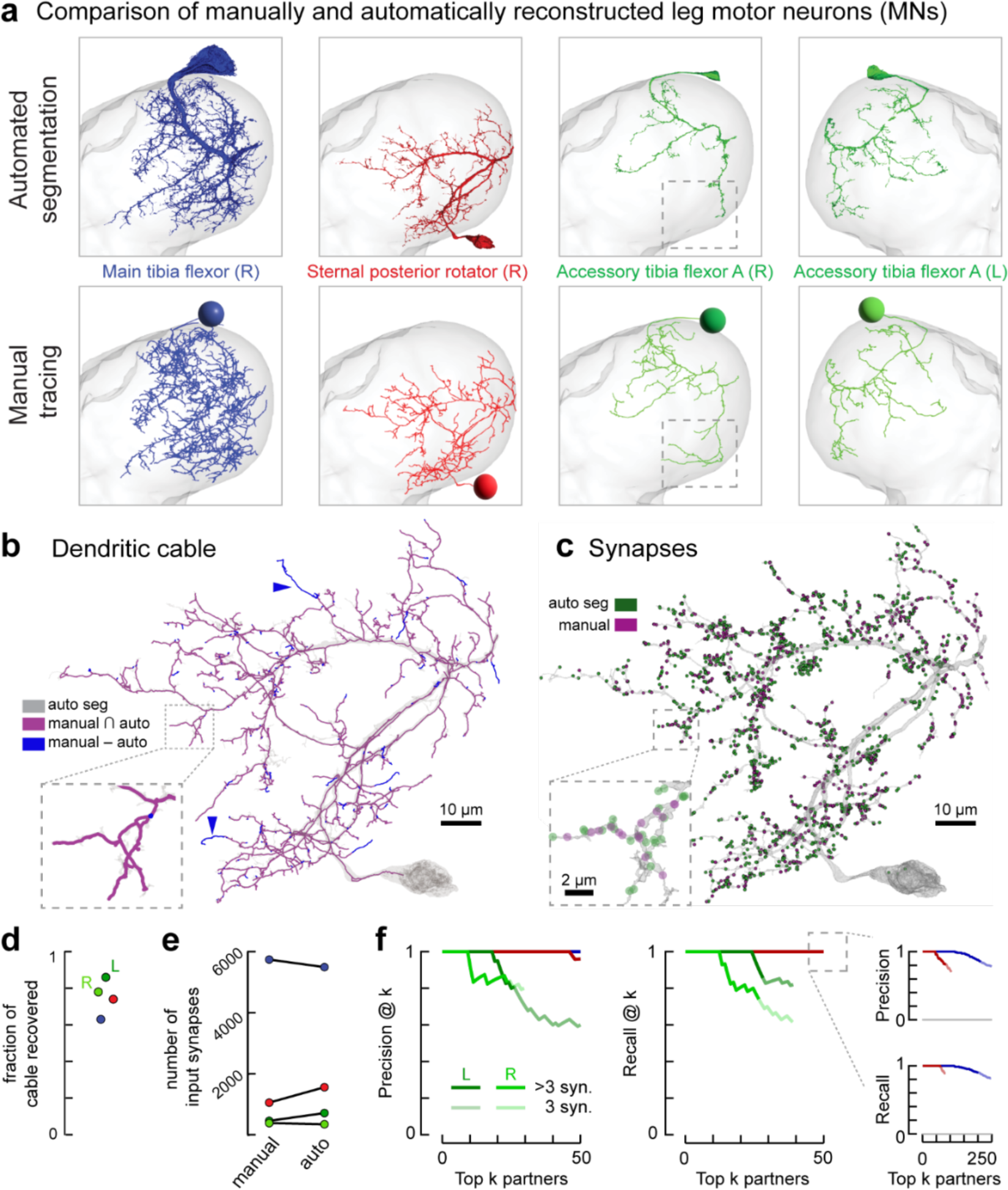
Validation of automated methods for segmentation and synapse prediction. **a,** To validate the accuracy of the segmentation, we compared automated and manual reconstruction for four leg motor neurons (MNs) innervating the T1 leg; three were from the right T1 neuromere (R) and one was from the left (L). Neurons were manually traced using CATMAID. Segmented neurons were initially proofread in Neuroglancer without knowledge of the manual “ground-truth”. **b,** Comparison of automated and reconstruction for the sternal posterior rotator MN. **c,** Comparison of automated synapse prediction and manual synapse annotation for the sternal posterior rotator MN. Note, manual synapse annotation identified additional neurites that had to be merged to the proofread reconstruction. Examples indicated by blue arrowheads, compare to **b**. **d,** The automated reconstruction effectively segmented most of the dendritic cable for all four MNs. The exceptions were typically fine dendritic branches, as illustrated in **b**. **e,** Input synapse counts from automated synapse prediction compared to manual annotation for all four MNs. **f,** Precision and recall of top presynaptic partners for each automatically reconstructed MN, for connections of three or more synapses. Dark colors indicate connections of more than three synapses, the light colors indicate connections with three synapses. Colors indicate MN identity as in **a**. For example, 80% of the top predicted presynaptic partners with more than 3 synapses onto the left accessory tibia flexor (dark green) are also partners according to the manual annotations (precision), and 80% of the ground-truth partners are found (recall). Inset: The precision and recall of the top 250 partners for the larger MNs.

Most differences between manual tracing and automatic segmentation occurred at fine branch endpoints (**Figure 2b**). The number of manually and automatically annotated synapses differed by <9% for the largest tibia flexor neuron and the left accessory tibia flexor (**Figure 2e**). For the other two MNs, automatic synapse detection found ∼50% more synapses for the other two MNs due to incomplete manual tracing in synapse-dense regions of the neuropil (**Extended Data Figure 3e**). The mean distance from a manually annotated synapse to a predicted synapse was less than 1.5 um (**Extended Data Figure 3b**). These results demonstrate the precision of automatic synapse prediction.

To assess how well the automatic segmentation and synapse prediction recover synaptic connectivity, we computed precision and recall curves for each of the four MNs (**Figure 2f**). We created lists of presynaptic partners from either the automatically predicted synapses or the manually annotated synapses, sorted the lists according to the number of synapses in the connection, and applied a threshold of three synapses (Hulse et al., 2021). Precision is the percentage of the top *k* predicted partners that are in the list of “true” partners based on manual synapse annotation (**Figure 2f, left)**. Recall is the fraction of the top *k* ground-truth partners that are in the list of predicted partners (**Figure 2f, right)**. For each MN, we found that >80% of predicted partners with more than three synapses are true partners (precision), and more than 80% of true top *k* partners with more than three synapses are predicted partners (recall).

In summary, we report the alignment, segmentation, and automatic synapse prediction for the FANC EM-volume. Our comparison with manual tracing and synapse annotation provides confidence that the automated segmentation, followed by appropriate proofreading (see Methods), provides a level of accuracy comparable to manual tracing. On average, proofreading MNs and their presynaptic partners required 20 edits per neuron. Proofreading automatically segmented neurons requires a fraction of the time required for manual tracing, thus dramatically accelerating reconstruction of the connectome.

### Identification of leg motor neurons (MNs) and target muscles

Knowing which MNs innervate which muscles is crucial to interpreting the function of motor circuits in the connectome. We therefore set out to identify the peripheral muscle targets of all MNs innervating the fly’s front (T1) legs. To reconstruct the MN projection map, we integrated information across three imaging datasets that collectively span the VNC and leg (**Figure 3a**). First, we identified and proofread all the MNs innervating the T1 leg in the FANC EM dataset (69 in left T1, 70 in right T1). Second, we traced the motor nerves and axons into their target muscles using an X-ray holographic nano-tomographic (XNH) dataset of the fly’s front leg (Kuan et al., 2020). Third, we screened a large collection of VNC neurons (Meissner et al., 2023) sparsely labeled with the multi-color Flp-out (MCFO) technique (Nern et al., 2015) to identify Gal4 driver lines (Jenett et al., 2012) labeling leg MNs. We imaged GFP expression of each genetic driver line in the front leg to identify the muscle target of each MN axon. We then compared the dendritic morphology of the genetically-labeled MNs to those reconstructed from FANC (**Figure 3**). We focused our efforts on the left MNs because the left prothoracic leg nerve (ProLN) of the FANC dataset is more intact than the right ProLN (**Extended Data Figure 4a-g**).

**Figure 3.**
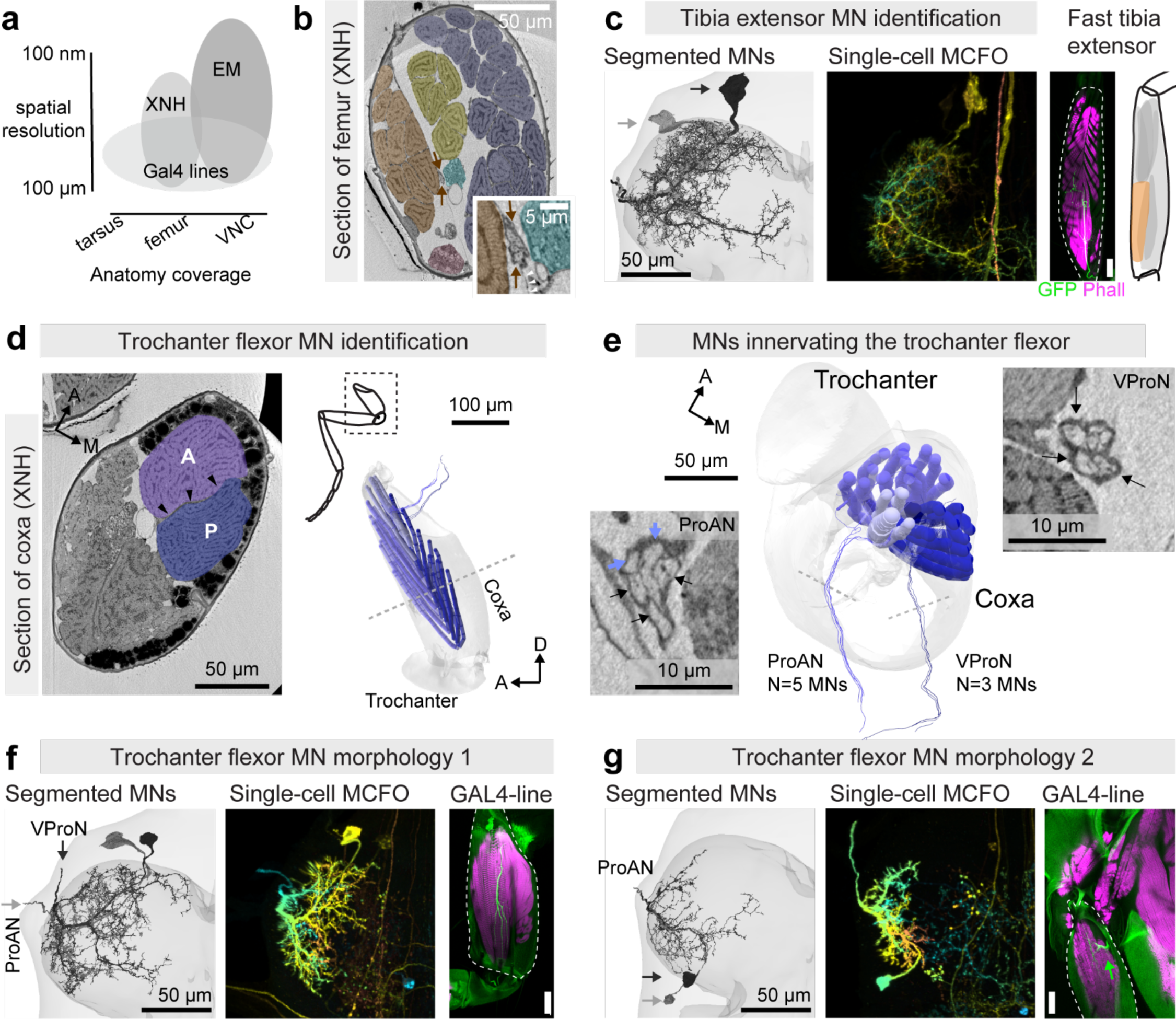
Matching motor neurons (MNs) in the connectome to leg muscle targets. **a,** Schematic of spatial resolution vs. coverage of anatomical tools. **b,** XNH cross-section of the femur, false-colored to indicate anatomy. Orange: tibia extensor muscle fibers; blue: tibia flexor muscle fibers; yellow: long tendon muscle fibers; pink: femoral chordotonal organ. Inset: magnified view of axons (arrows) passing through a fascia membrane (arrowheads). **c,** Identification of SETi (light) and FETi (dark) MNs. Left: tibia extensor MNs reconstructed in FANC. Center: depth-colored maximum-intensity projection (MIP) of a single MN labeled by multi-color FLP-out (MCFO; VT017399-Gal4). Right: GFP expression in the femur, driven by VT017399-Gal4 (green, projection), with phalloidin staining of the muscle (single section), schematized at right. Scale bar is 50 μm. **d,** Left: XNH cross-section through coxa. Arrowheads indicate the trochanter (tr.) flexor tendon. The tr. flexor tendon separates the anterior muscle fibers (light blue) from posterior fibers (dark), with proximal fibers (pale blue) inserting at the proximal tip of the tendon (not shown). Right: Annotated tr. flexor muscle fibers in XNH. **e,** Projected 3D view along the long axis of the coxa. Three MNs leave the ventral prothoracic nerve (VProN) to innervate the posterior fibers (right inset). Five MNs leave the prothoracic accessory nerve (ProAN) to innervate the anterior fibers (left inset); two traced MNs in ProAN innervating proximal fibers are indicated with blue arrows. **f,** Two of six FANC MNs with characteristic morphology, one of 3 that exit via ProAN, and one of 3 that exit via VProN. MCFO clones and GAL4-driven GFP in coxa (VT063626-Gal4). **g,** Two FANC MNs with small posterior somas. MCFO clones and GAL4-driven GFP in coxa (VT025963-Gal4) provide evidence that at least one MN with a posterior soma innervates the proximal fibers of the trochanter flexor muscle.

We illustrate the process of MN identification across the three imaging modalities with two examples (**Figure 3**). First consider the fast and slow tibia extensor MNs (**Figure 3b, c**) that innervate the tibia extensor muscle. These neurons have been studied extensively in other insects, where they are referred to as the FETi and SETi (Bässler and Büschges, 1998; Burrows, 1996). In the XNH volume, we identified two candidate MN axons that leave the leg nerve in the femur and pass through a membrane separating the tibia extensor muscles from the tibia flexor muscles (arrows in **Figure 3b**). We identified four GAL4 lines from the Janelia database (Meissner et al., 2023) labeling two MNs that project through the same nerve to innervate the extensor muscles. The dendritic morphology of the genetically labeled MNs closely matched the dendritic morphology of two MNs that we reconstructed in the FANC EM dataset (**Figure 3c**). We therefore concluded that these are the FETi and SETi MNs. As in the locust (Burrows and Horridge, 1974), they can be distinguished from each other because the FETi branches more extensively than the SETi in the VNC and innervates more and larger muscle fibers in the femur. The SETi receives 7090 input synapses, and the FETi receives 14,904, the most for any front leg (T1) MN. The only other neuron in FANC with more synapses is the hind leg FETi, with 15,474 input synapses.

As a second example, consider the large trochanter flexor muscle in the coxa (**Figure 3d**). Compared to the tibia, whose extension is controlled by the SETi and FETi MNs, little is known about the neural control or biomechanics of the trochanter. The trochanter flexor tendon bisects the muscle, with anterior muscle fibers attaching to one side of the tendon and posterior fibers attaching to the other (**Figure 3d**). We traced eight MNs innervating this muscle in the XNH volume: five innervate the anterior fibers via the prothoracic accessory nerve (ProAN) and three innervate the posterior fibers via the ventral prothoracic nerve (VProN) (**Figure 3e**). In FANC, we found six MNs with morphology that resembled previous images of trochanter flexor MNs (Enriquez et al., 2015): three that leave via the ProAN and three that leave via the VProN (**Figure 3f**). If we assume the same number of trochanter flexor neurons exist in both FANC and XNH datasets (from different flies), what about the two other MNs that innervate the muscle from the ProAN? In FANC, twelve MNs exit via the ProAN, six have somas on the posterior cortex of the neuropil, four of which likely innervate the sternal posterior rotator muscle. Confocal imaging of genetic driver lines showed that at least one MN with a posterior soma innervates the proximal fibers of the trochanter flexor (**Figure 3g**) (Baek and Mann, 2009). From these strands of evidence, we deduced the identity of all eight MNs that innervate the trochanter flexor muscle.

We applied a similar process of deduction to map all 69 left T1 MNs to 18 target muscles (**Figure 4**). A comprehensive description of leg MN identification from EM, XNH, and light-level imaging of genetic driver lines is provided in the **Supplementary Methods**. We also used the XNH dataset to determine the number of muscle fibers in each leg muscle (**Figure 4a**), along with fiber origins and cuticle attachment points. Combining this anatomical information with joint kinematics recorded from walking flies (**Figure 4b**) (Karashchuk et al., 2021), leads to predictions of how MNs may contribute to distinct aspects of the step cycle, such as leg swing vs. stance (**Figure 4c**). We note, however, that the precise timing of MN spikes can dictate how muscles contribute to behavior (Dickinson et al., 2000). Because the dendritic morphology of leg MNs is stereotyped from fly to fly, this atlas (**Figure 4c-d**) can be used to identify MNs in future datasets. Indeed, it has already been used to identify MNs in a separate EM volume of the male VNC (MANC), including across different VNC segments (Cheong et al., 2024).

**Figure 4.**
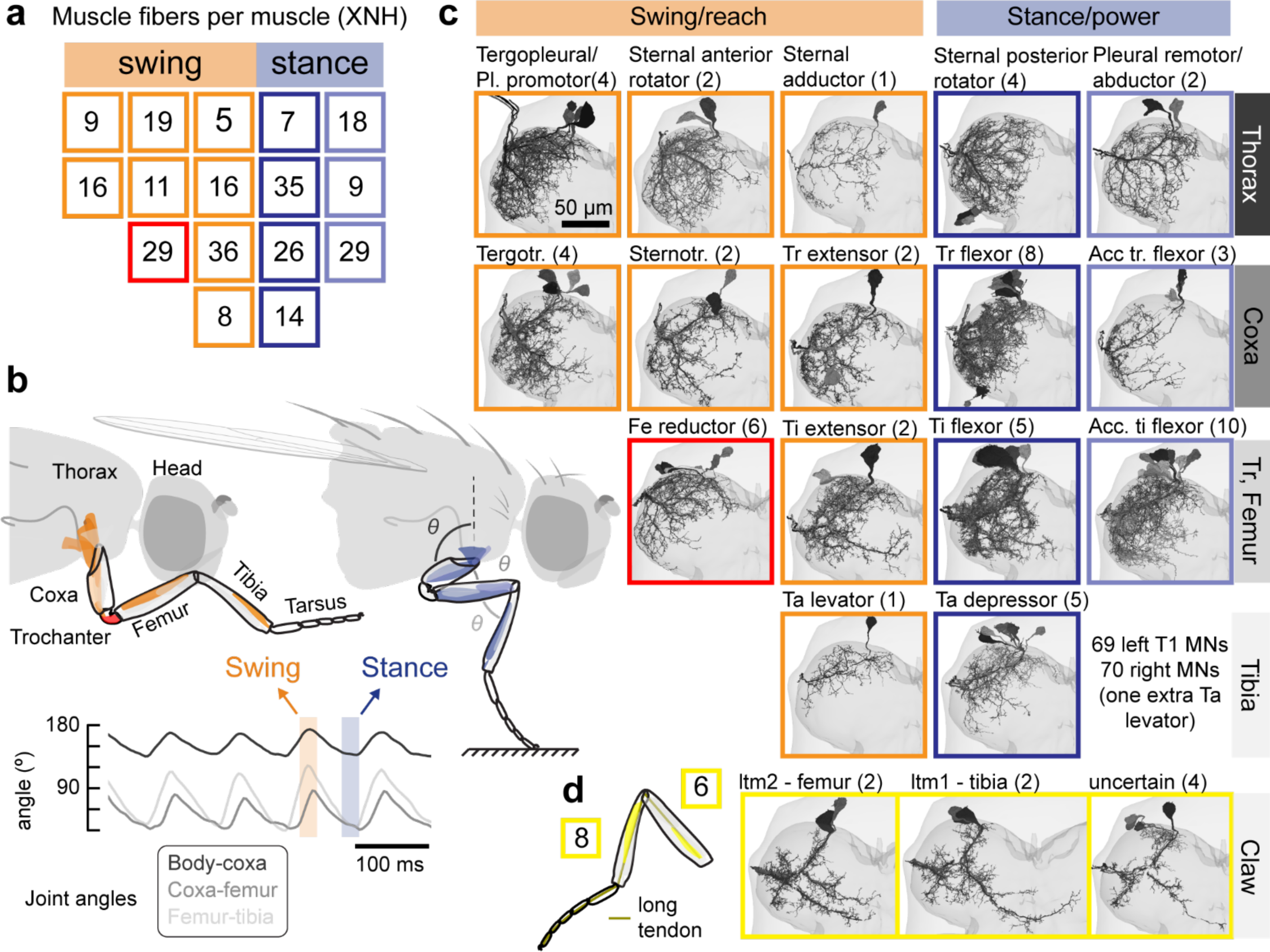
Identification of the specific muscle innervated by each left T1 motor neuron (MN) in FANC. **a,** Counts of fibers in each leg muscle from XNH volume. Color code for muscles is shown in **c**. **b,** Schematic of musculature, separated into muscle groups that drive the leg to swing forward or reach (orange), vs. muscles that push the fly’s body forward (blue). Inset: joint angles measured from a fly walking on a spherical treadmill (data from Karashchuk et al., 2021). **c,** EM-reconstructed MNs, grouped by segment (rows, labeled at right) and by muscle target (each square). Gray scale indicates different MNs, orange vs. blue indicates swing vs. stance. The femur reductor MNs that target the trochanter, whose function is unknown, are indicated by a red square. **d,** Left: schematic of the long-tendon muscle (ltm), a multi-joint muscle with fibers in both the femur (ltm2) and tibia (ltm1) that insert onto the same long tendon (retractor unguis, dark line in the schematic) and control the tarsal claw. Left: four ltm MNs have extensive medial branches; two target ltm2, two target ltm1. The specific targets of four smaller ltm MNs cannot be resolved from the XNH volume.

In the process of reconstructing and identifying leg MNs, we made other novel observations that were only possible due to the reconstruction of neuromuscular anatomy across imaging scales (**Figure 3a**). For example, the right front leg of FANC contained 70 MNs (compared to 69 in the left T1). The extra cell appears to be a second tarsus levator MN. Supernumerary MNs have been previously described in the locust leg motor system (Siegler, 1982), but have not been described in flies. Further, six MNs innervate the femur reductor muscle in the trochanter. It was previously assumed that the *Drosophila* trochanter and femur are functionally fused (Hartenstein, 2006); however, the fact that the trochanter segment still possesses its own dedicated MNs and musculature suggests that either the two segments retain some independent biomechanical function or that the current morphology represents a palimpsest of the ancestral state.

In cases where we were able to trace MN axons to their specific target muscle fibers, we found that large (fast) MNs innervate more and larger muscle fibers than small (slow) MNs (e.g. **Supplementary Methods Figures 11 and 12**). This relationship, which has also been observed in other vertebrate and invertebrate species (Kernell, 2006; Snodgrass, R.E., 1935), is consistent with a gradient in force production across muscle fibers.

We found that some muscle fibers are innervated by multiple MNs, confirming that adult fly muscles can exhibit polyneuronal innervation, as is the case in locusts (Hoyle, 1983) and larval *Drosophila* (Landgraf et al., 1997). Whereas mammalian muscle fibers are often poly-innervated in neonatal animals, they are pruned during development, leading to singly-innervated fibers in adults (Brown et al., 1976). We found clear evidence from the XNH data for polyneuronal innervation in the long tendon muscle (LTM), the proximal fibers of the trochanter flexor (**Figure 4e**, not shown), and the femur reductor muscle. LTM fibers originate in both the femur (ltm2) and the tibia (ltm1) and insert on the long tendon (retractor unguis), which extends from the femur to the claw at the distal tip of the leg (**Figure 4g**, gold line) (Radnikow and Bässler, 1991). Polyneuronal innervation may endow the motor system with more flexibility and may help explain how insect limbs achieve precision despite their sparse MN innervation. Another mechanism for enhancing motor flexibility in other arthropods is the presence of GABAergic MNs that innervate many target muscles (Wolf, 2014); however, we did not find such “common inhibitor” MNs in the fly leg.

We did not observe any chemical output synapses from leg MNs in the VNC, consistent with past work in the locust (Burrows et al., 1989). This lack of output synapses is notable because the vast majority of fly neurons possess apposed input and output synapses, even in dendritic compartments. Larval fly MNs innervating abdominal muscles also lack output synapses (Mark et al., 2021; Schneider-Mizell et al., 2016). However, larval MNs do appear to form gap junctions in specific segments, allowing motor neurons to participate in rhythm generation (Matsunaga et al., 2017). In other arthropods, such as crustaceans, the central output synapses of MNs are essential for their role in rhythmic pattern generation (Marder et al., 2017).

### Identification of wing motor neurons

We next focused our analysis on the wing motor system. Wing muscles (**Figure 5a**) can be broadly divided into three groups based on their functional roles in flight (Dickinson and Tu, 1997; Pringle, 1957). First, 13 large muscle fibers power flight indirectly by causing the thorax to resonate at high frequencies, which changes the conformation of the mechanically complex wing hinge to drive the wings to flap back and forth (Deora et al., 2017). The 13 fibers are organized into four muscles on each side, a single dorsal-longitudinal muscle (consisting of 6 fibers, DLM1-6), and three dorsoventral muscles (consisting of three fibers (DVM1), and two fibers (DVM2 and DVM3)). A second set of 12 steering muscles attach directly to the hardened cuticle elements (sclerites) of the wing hinge to rapidly adjust wing motion. Third, the lateral thorax is equipped with four tension muscles, which, although not directly connecting to the sclerites at the base of the wing, are thought to adjust wing motion via their effects on the mechanical stiffness of the thorax (Nachtigall and Wilson, 1967; Pringle, 1957).

**Figure 5.**
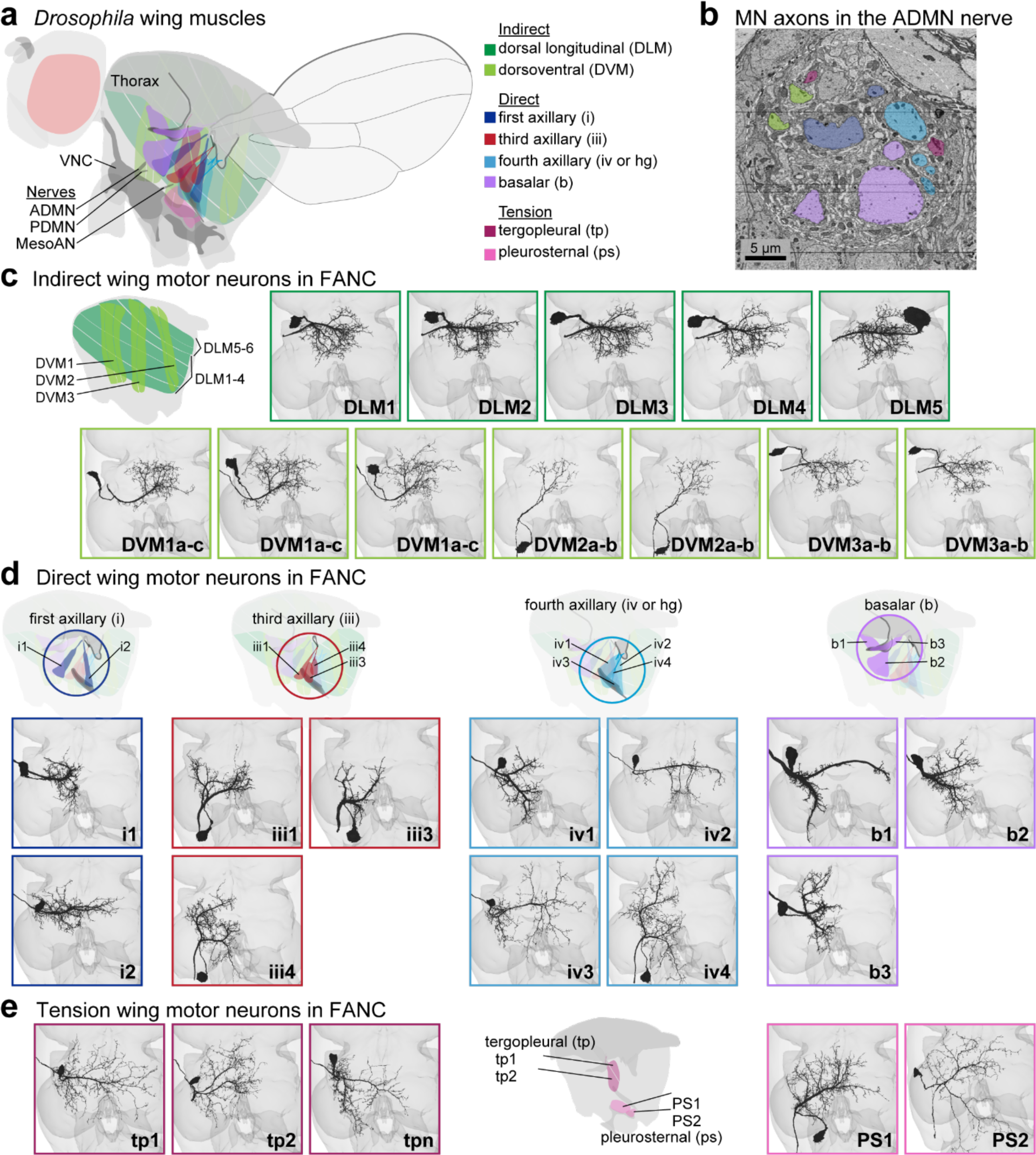
Identification of wing motor neurons (MNs) in FANC. **a,** Schematic showing muscles that power and finely control wing motion. Indirect power muscles (green) span the thorax along the anterior-posterior and dorsal-ventral axes; their antagonistic contractions resonate the thorax at high frequencies and cause the wings to move back and forth during flight. Direct muscles attach directly to sclerites of the wing hinge and finely adjust the wing motion. Tension muscles attach to the inner wall of the thorax and internal apodemes, and their contractions are thought to alter the tension of the thorax, thus modifying the oscillations generated by the indirect power muscles. Also pictured is the VNC, with the nerves that carry MN axons to wing muscles (PDMN, ADMN, MesoAN). **b,** EM image from FANC showing a cross section of the ADMN. MN axons are colored according to the key in **a**. MNs that innervate muscles with similar attachment points often fasciculate together in the nerve. Horizontal black lines are due to missing slices from the serial-sectioned reconstructed volume. **c,** Segmented and proofread MNs from FANC that innervate indirect MNs. We could not differentiate the MNs that innervate individual DVM fibers within each muscle (DVM 1, DVM 2, or DVM 3), so they share a common label (i.e. DVM 1a-c refers to all three MNs that innervate DVM 1). See methods for details on identification for **c-e**. **d,** Segmented and proofread MNs from FANC that innervate direct muscles. **e,** Segmented and proofread MNs from FANC that innervate tension muscles.

We identified the MNs that innervate wing muscles by locating neurons in FANC with cell bodies in the VNC and axons that leave through one of the three nerves associated with the wing neuropil (**Figure 5b**): the anterior dorsal mesothoracic nerve (ADMN), posterior dorsal mesothoracic nerve (PDMN), and the mesothoracic accessory nerve (mesoAN) (Court et al., 2020; Phelps et al., 2021). We found 37 neurons on each side, 29 of which we determined to be wing MNs (the other eight are discussed below and in **Extended Data Figure 5a-c**). We identified left-right homologous pairs, but here we focus our analysis on the left side.

As for the legs, we identified the muscle targets of all the wing MNs in FANC. Most identifications were based on dendritic morphology from the literature, including MNs that innervate direct and tension muscles (Cheong et al., 2024; Ehrhardt et al., 2023; O’Sullivan et al., 2018; Trimarchi and Schneiderman, 1994), and MNs that innervate indirect muscles (Ikeda and Kaplan, 1974; Schlurmann and Hausen, 2007) (**Figure 5c**).

The MNs of the DLMs were the only MNs we found to possess presynaptic sites in the VNC (**Extended Data Figure 5d**). These synapses were rare (∼10 predicted synapses in total). The presence of weak synaptic connectivity between DLM MNs complements recent work demonstrating that gap junctions between DLM MNs offset their spike timing during flight (Hürkey et al., 2023).

We found three leg MNs and four efferent neurons that innervate peripheral nerves, but which we did not categorize as MNs. One is the previously described peripherally synapsing interneuron, which plays an important role in escape behavior (King and Wyman, 1980). Three are previously undescribed efferent neurons, one of which has a branch that ascends to the brain (**Extended Data Figure 5a-c**).

### A neural circuit to retract the front legs during escape take-off

With knowledge of the muscle targets of the front leg and wing MNs, the connectome points to clear hypotheses about the physiological and behavioral function of premotor circuits. We illustrate the potential of combining the VNC connectome and MN projection map with identification and analysis of premotor neurons that coordinate the fly’s legs and wings during escape takeoff (**Figure 6a**).

**Figure 6.**
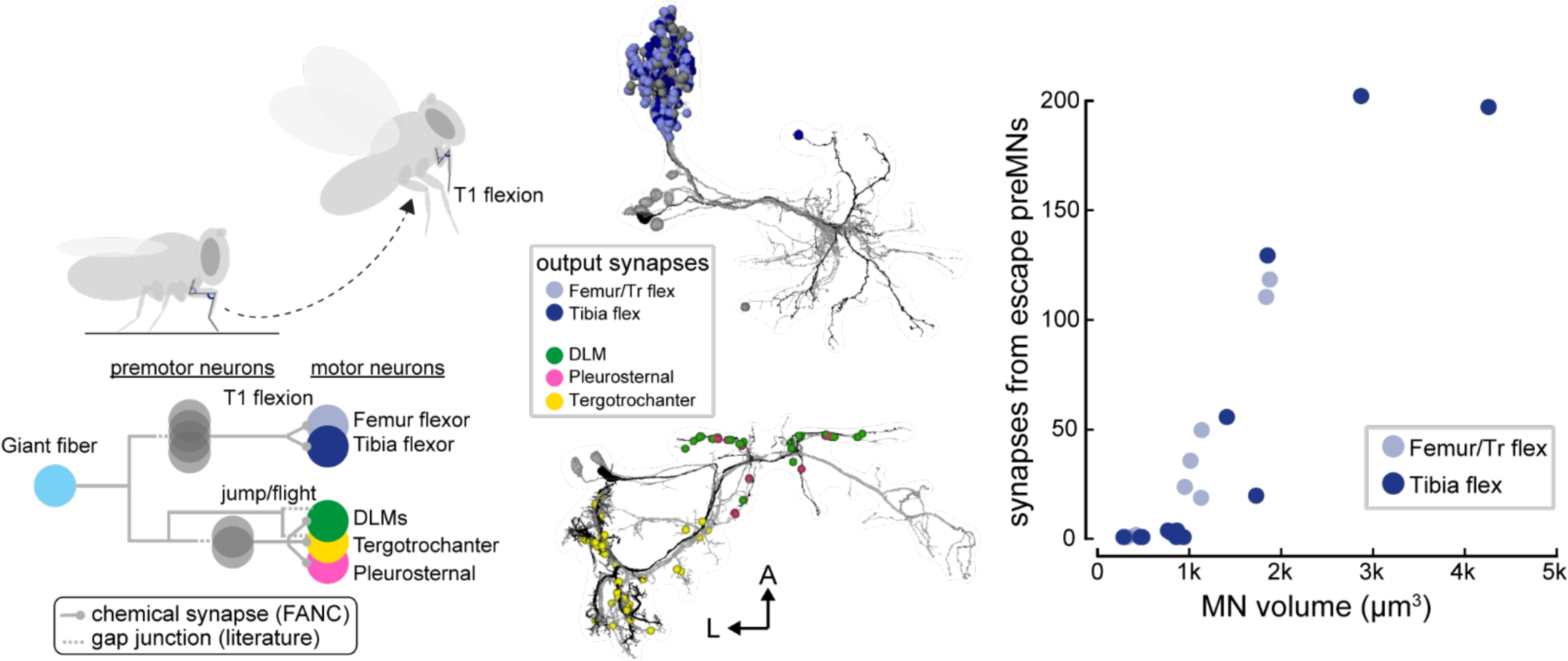
Circuits that coordinate the wings and legs during escape takeoff. **a,** Schematic of a proposed circuit for escape based on prior literature and FANC synapse predictions. The Giant Fiber (GF) excites wing and middle (T2) leg MNs as well as premotor neurons (preMNs) through gap junctions. In FANC, preMNs that are electrically coupled to the GF target T1 tibia and trochanter-femur flexor MNs as well as MNs that innervate T2 leg tergotrochanter muscles, DLMs, and thorax tension muscles (pleurosternal). **b,** We identified interneurons that have been previously shown to be electrically coupled to the giant fiber. (Top) GFC4 preMNs from hemilineage 11A. (Bottom) GFC2 preMNs from hemilineage 18B. The interneurons make synapses (spheres) onto MNs that drive jumping in the T2 legs (yellow), flexion of the T1 legs (blue), and initiate the flight motor (green and pink). A single interneuron in each group is represented in black to show morphology. Additional colors in T1 indicate synapses onto MNs other than tibia and trochanter flexors. **c,** Premotor neurons that are electrically coupled to the giant fiber exclusively target the largest MNs innervating tibia and trochanter-femur flexor muscles, suggesting that this circuit motif bypasses the recruitment hierarchy to execute the fast high-force movements necessary for escape.

Previous research identified a class of VNC local neurons (giant fiber coupled or GFC) that are electrically coupled to the giant fiber (Kennedy and Broadie, 2018), a large descending neuron that drives evasive escape takeoff (Tanouye and Wyman, 1980; von Reyn et al., 2014). In the FANC volume, we identified and proofread two subtypes of GFC interneurons (GFC2 and GFC4). We then used FANC synapse predictions to understand how they are synaptically connected to leg and wing MNs (**Figure 6b**). The morphology of the GFC2 and GFC4 interneurons indicates that they develop from stem cell hemilineages (18B and 11A, respectively) that release the excitatory neurotransmitter acetylcholine (Lacin et al., 2019).

We found that the GFC2 interneurons synapse onto indirect and tension wing MNs as well as tergotrochanter (TT) MNs that innervate the middle leg (T2), which is consistent with previous literature (Tanouye and Wyman, 1980). In addition to its role in T2 trochanter-femur extension during jumping, the enormous TT muscle is also thought to quickly initiate the first cycle of mechanical oscillation in the power muscles that maintain flight (Nachtigall and Wilson, 1967). Thus, GFC2 interneurons are positioned to excite MNs that drive two parallel motor programs: jumping and initiation of the flight motor. The other interneuron subtype, GFC4, synapses onto front leg (T1) MNs that flex the trochanter-femur or tibia (**Figure 3d-g**). This connectivity suggests that descending input from the giant fiber drives synchronized extension of the middle legs (jumping) and flexion of the front legs (lift-off). Although the role of the T1 leg retraction during take-off has not been previously described, it is visible in high-speed videos of *Drosophila* escape (e.g. von Reyn et al. 2014, Supplementary Video 1) (Card and Dickinson, 2008; Trimarchi and Schneiderman, 1993; von Reyn et al., 2014).

Leg MNs in *Drosophila* and other animals are typically recruited in a hierarchical sequence, with small, low-force producing MNs firing prior to large, high force producing MNs. In paired electrophysiological recordings from tibia flexor MNs, we previously observed occasional violations of the recruitment order, in which MNs at the top of the hierarchy fire before MNs at the bottom (Azevedo et al., 2020). We hypothesized that these violations could occur during ballistic escape behaviors, such as escape takeoff. Consistent with this prediction, the only MNs that receive synaptic input from GFC interneurons are the largest MNs that innervate femur-trochanter and tibia flexor muscles (**Figure 6c**). Thus, our analysis of the connectome suggests that descending input from the giant fiber is positioned to subvert the standard leg MN recruitment order to drive rapid, ballistic escape behavior via the GFC4 interneurons.

## Discussion

By revealing the structure of the neurons and synapses that underlie animal behavior, connectomes provide a foundation for generating and falsifying theories of neural circuit function. While comprehensive connectomes have been mapped for small crawling animals like *C. elegans* (Cook et al., 2019) and the *Drosophila* larva (Winding et al., 2023), we previously lacked a synapse-level wiring diagram of motor circuits for any limbed animal. Here, we provide tools for collaboratively proofreading a draft connectome of a female adult *Drosophila* VNC, an atlas linking the outputs of the connectome to the body, and analysis of a circuit that coordinates the legs and wings during escape behavior. Our effort to reconstruct the connectome of a female VNC (FANC) complements a parallel project to reconstruct a male VNC (MANC) (Cheong et al., 2024; Marin et al., 2023; Takemura et al., 2023). The existence of multiple VNC connectomes will enable comparison of neural wiring across individuals and the identification of sexually dimorphic neural circuits.

We identified the muscle innervation of leg and wing MNs by combining data from EM, X-ray nanotomography, light-level imaging of genetic driver lines, and past literature (Baek and Mann, 2009; Brierley et al., 2012; Cheong et al., 2024; Ehrhardt et al., 2023; Enriquez et al., 2015; Ikeda and Koenig, 1988; Kuan et al., 2020; Meissner et al., 2023; O’Sullivan et al., 2018; Phelps et al., 2021; Schlurmann and Hausen, 2007; Trimarchi and Schneiderman, 1994; Venkatasubramanian et al., 2019). We demonstrate the necessity of combining the connectome and MN projection map by quantifying premotor connections that are positioned to coordinate the legs and wings during escape takeoff. We combined measurements of synaptic connectivity with measurements of cell volume and found that descending escape signals may subvert the typical MN recruitment hierarchy by preferentially exciting fast (large) MNs over slow (small) MNs.

The community effort to comprehensively reconstruct neurons and measure synaptic connections within FANC is ongoing. At the time of publication, the community has proofread less than half of the total FANC dataset, yet the automated reconstruction, proofreading environment, and analysis tools already enable interactive inquiry into the neurobiology of the VNC. Our work parallels similar efforts in the adult fly brain (Dorkenwald et al., 2023a), the male VNC (Cheong et al., 2024; Marin et al., 2023; Takemura et al., 2023) and the larval nervous system (Eschbach and Zlatic, 2020). Further, the distributed nature of the community-driven FANC effort is a strength, as it allows the connectome to be continuously refined as each circuit is proofread by experts in that area. Even before the connectome is complete, researchers can analyze subcircuits as a means of generating and falsifying hypotheses. Using tools to match VNC neurons reconstructed in the EM volume with GAL4 lines imaged by light microscopy (**Extended Data Figure 2**), it is then possible to test these hypotheses with *in vivo* recordings and manipulations of neural activity. Additional software tools exist to facilitate comparison of neuronal morphology and connectivity across individual animals and to bridge EM datasets (Schlegel et al., 2023). Even with a single connectome, however, it is possible to gain insight into morphology and neural circuitry that differ between related species. For example, our reconstruction revealed independent control of the femur and trochanter leg segments, which were previously thought to be mechanically fused in *Drosophila* (Hartenstein, 2006).

Many challenges remain for interpreting a synaptic wiring diagram (Bargmann and Marder, 2013). First, synapses can be either excitatory or inhibitory. Fortunately, VNC neurons develop from stem cell hemilineages that share identifiable morphological features as well as neurotransmitter identity (Lacin et al., 2019). Therefore, hemilineage identification of VNC cell types can identify candidate neurotransmitters, as we did for the escape circuitry (**Figure 6**). As done previously for the whole fly brain EM dataset, we can also use hemilineage identities as ground-truth for training convolutional neural networks to predict neurotransmitter identity directly from the EM image data (Dorkenwald et al., 2023a; Eckstein et al., 2020).

A second challenge is the inability to see electrical synapses or sites of neuromodulation in connectome datasets. The spatial resolution of existing connectomic datasets is insufficient to resolve gap junctions, which are known to play an important role in sensorimotor processing within the VNC (Agrawal et al., 2020; Chen et al., 2021; Tanouye and Wyman, 1980). Sample-preserving imaging approaches like serial-section TEM have the capacity for higher-resolution (e.g., 2 nm/pixel) imaging at identified locations, potentially allowing visualization of gap junctions. It is also not currently possible to identify sites of neuromodulation, which are essential for flexible motor control (Howard et al., 2019). Because EM reconstruction provides a map of chemical synaptic transmission between neurons, physiological measurements will also be necessary to test for the effects of gap junctions and neuromodulators.

Many organisms use their limbs to maintain posture in the face of gravity, navigate uneven terrain, and manipulate objects in the environment. Connectomic-driven discovery represents a paradigm shift for investigating these fundamental motor control problems. Previous work on the neural control of movement has attempted to link neural dynamics to motor output, in part to infer the connectivity and function of premotor networks (Kawai et al., 2015; Lindén et al., 2022; Sauerbrei et al., 2020; Shenoy et al., 2013; Vyas et al., 2020). A connectome allows one to work backwards from the MNs to identify premotor networks that coordinate specific patterns of muscle activation and inhibition. The adult fly VNC also provides a unique opportunity to investigate two distinct modes of limbed locomotion, walking and flight.

The FANC connectome will have many uses beyond the study of limb motor control. For example, it will enable the identification and analysis of neural circuits for descending modulation (Aymanns et al., 2022; Namiki et al., 2018), ascending communication with the brain (Chen et al., 2022; Cheong et al., 2024), and sensory organs distributed across the fly limbs, thorax, and abdomen (Tuthill and Wilson, 2016). By creating a bridge between the VNC connectome and the body, the MN projection map will facilitate development and analysis of neuromechanical models for flexible motor control (Lobato-Rios et al., 2022; Wang-Chen et al., 2023). The compact sensorimotor circuits that mediate robust control of the fly leg and wing may provide inspiration for engineering of micro-scale robotic systems (Dallmann et al., 2023). Matching cell types and comparing connectivity motifs to the larval connectome (Mark et al., 2021; Winding et al., 2023; Zarin et al., 2019) may provide insight into the development and evolution of sensorimotor circuits (Agrawal and Tuthill, 2022; Heckscher et al., 2015). Finally, as volumetric EM datasets from other animal species are collected, comparative connectomics provides a powerful tool to study the evolution of the nervous system (Barsotti et al., 2021).

## Acknowledgements

We thank Rachel Wilson for financial support of SG during development of braincircuits.io (via U19NS104655 and R01NS129647 to RIW). We thank Richard Mann, Han Cheong, Erica Ehrhardt, Gwyneth Card, and Greg Jefferis for assistance with leg and wing MN identification. This work was supported by a Searle Scholar Award, a Klingenstein-Simons Fellowship, a Pew Biomedical Scholar Award, a McKnight Scholar Award, a Sloan Research Fellowship, the New York Stem Cell Foundation, and a UW Innovation Award to JCT; a Genise Goldenson Award to WAL; NIH U19NS104655 to JCT and MD; NIH R01MH117808 to JCT, WAL, and HSS. JCT is a New York Stem Cell Foundation – Robertson Investigator.

## Methods

### EM Dataset Alignment, Segmentation, Annotation

We refined the alignment of the FANC image data using self-supervised convolutional neural networks (Popovych et al., 2022). We next trained a CNN to identify knife marks that occurred during serial sectioning. We then used CNNs to segment the dataset into neurons and fragments of neurons and to predict synapse locations (Macrina et al., 2021), excluding regions of the data with knife marks. We also used a CNN to predict pre and postsynaptic partners (Buhmann et al., 2021). The automated segmentation was ingested into the ChunkedGraph data structure (Dorkenwald et al., 2022).

We imported synapse and cell body predictions into the Connectome Annotation Versioning Engine (CAVE), so that the associated cell segmentation objects for each annotation would be dynamically updated during proofreading (Dorkenwald et al., 2023b). This system allowed us to query the up-to-date connectivity graph and associated metadata, such as cell-type annotations. A suite of tools we created for analysis of the FANC dataset are described in **Extended Data Figure 2**.

To assess the quality of the FANC automated segmentation and synapse prediction, we manually traced and annotated synapses for a subset of leg MNs using CATMAID (Schneider-Mizell et al., 2016).

### Proofreading

We corrected errors in the segmentation through manual proofreading with Google’s Neuroglancer interface (Maitin-Shepard et al., 2021). We adopted a two-pass heuristic to proofread MNs: a first pass to correct major errors, and a second pass to more closely inspect the neuron morphology. Examples of major errors included a missing soma, requiring merging two objects; a missing branch, e.g. from the primary neurite, requiring merging the branch onto the larger MN object; or incorrect merges with glia, which wrap around the MN axons in the nerve and can be segmented with the MN, requiring splitting the object (Dorkenwald et al., 2022) (**Extended Data Figure 2a**). These major errors were typically due to two types of artifacts in the EM data that impacted the segmentation and reconstruction. First, the serial sectioning procedure left several knife marks that could cross an entire neuropil. These knife marks were identified and excluded from the segmentation, but branches that cross the knife marks were segmented as separate objects. The left T1 neuromere in particular had several large knife marks which could lead to somas or large branches being detached from the neurons. The second type of artifact was the darker appearance of the cytosol of many peripheral axons. This led to merges between glia and the axons of sensory and MNs.

After a first pass, e.g. to reattach somas, we made a second pass over the MN morphology to correct smaller errors like incorrectly merged branches that in fact belong to other objects (requiring a split), or merging smaller branches that branched distally from the primary neurite. Common causes of these errors include darkly stained lipid aggregates in the EM image which often occurred where two neurites were closely apposed, and the segmentation algorithm could incorrectly identify the process that emerged on the other side.

The MNs were also initially manually traced in CATMAID (Phelps et al., 2021). We took advantage of this to import the manually traced skeletons for each MN and check that no further branches were missing. Note, the four MNs we used to ground-truth synapse prediction and reconstruction were extensively manually traced to include the “twigs” at the ends of the dendrites, though some were missed in synapse dense regions (**Extended Data Figure 3e**). That level of detail was too time consuming to apply to all the MNs, and unnecessary given the improved accuracy of the segmentation. As a result, proofread MNs tended to have more twigs and more extensive dendrites than the manually traced skeletons.

Currently, ∼30% of non-sensory neurons have been marked as proofread by the community. The distributed responsibility of community-based proofreading stems from a need to pool resources across the community, in lieu of building a dedicated connectomics team charged with proofreading. As a result, circuits of more direct interest to the community have likely received more attention. On the other hand, the proofread reconstruction remains dynamic and circuits that are particularly difficult to segment can still be proofread, like the sensory neurons in FANC.

### Automatic detection of neuron nuclei

We used a 3D CNN to detect neuronal cell bodies (Mu et al., 2021). Our 3D CNN estimated the probability that a 64 ✕ 64 ✕ 45 nm^3^ voxel belonged to a nucleus (Lee et al., 2017). Training labels were generated by selecting automatically segmented nuclei from the cell segmentation. The probability map was thresholded at 0.7, connected component labels were generated, and centroid and size for each connected component were computed. Objects with size dimension below (1376 nm, 1376 nm, 1800 nm) for (x, y, z) were excluded, leaving 17,076 putative nuclei (**Figure 1d**).

In the automatic cell segmentation, the nucleus was frequently segmented as a separate object from the remainder of the cell. To assist with analysis, we merged the nucleus with its cell within the cell segmentation. For each object in the dedicated nucleus segmentation, we shifted the dedicated nucleus segment by one 68.8 ✕ 68.8 ✕ 45 nm^3^ voxel in every direction and identified the most frequent object within the cell segmentation to determine the associated cell. (**Figure 1f**).

### Validation of the nucleus segmentation results

Even with the size threshold, we still found some glia and non-nucleus objects in the list of 17,076 putative neuronal nuclei. We manually inspected each nucleus and its associated cell. Following three rounds of quality checks, we categorized 14,621 neurons, 2,030 glia, and 410 false positives (**Extended Data Figure 1a**). Details regarding quality checks can be found at the git-hub repository for this paper (github.com/EllenLesser/Azevedo_Lesser_Phelps_Mark_2023/).

### Identification of leg MN targets

The identification of leg MNs is described in detail in **Supplementary Methods**, the atlas of FANC left T1 MNs. To identify MNs, we used three imaging datasets that collectively span the VNC and leg (main **Figure 3a**). We first compared the easy-to-characterize features, like whether the soma was anterior or posterior of the neuropil, and which of the four nerves the axon exits. Then, we used a computational analysis from Phelps et al. (2021) (Phelps et al., 2021), which relied on the NBLAST algorithm(Costa et al., 2016) to identify bundles, or groups of neurons with primary neurites that run together from the soma to the axon. Phelps et al. limited this analysis to the primary neurites of the manually-traced MNs. As a result, a large bundle in the leg nerve in fact included neurons with distinguishing dendritc morphology, like the extensive medial dendrites of the long-tendon MNs. We used these obvious dendritic features to distinguish specific groups next. Careful visual inspection was the final step.

We developed an independent, parallel quantitative analysis that corroborates the MN-to-muscle matches (**Supplementary Methods Figure 18**). In this analysis, we divided the T1 neuromere into 8 µm cubic voxels and calculated the synaptic density of each MN in each voxel (1891 voxels x 69 MNs). We then reduced the dimensionality of the voxel matrix to 2D using the UMAP algorithm (McInnes et al., 2020). The algorithm created clear clusters of MNs that innervate synergistic muscles, confirming our assignments based on cross-modality comparison of morphological features.

**Supplementary Table 1** contains links to view leg motor neurons in neuroglancer.

### Identification of wing MNs

Unlike the leg, previous literature has already linked fly MN morphologies to muscle targets using muscle back-fills and specific GAL4 drivers (Ikeda and Koenig, 1988; O’Sullivan et al., 2018; Schlurmann and Hausen, 2007; Trimarchi and Schneiderman, 1994), so we were able to identify many of the neurons based on morphology alone. We identified wing and thorax MNs in FANC by finding neurons with cell bodies in the VNC and axons leaving through one of three wing-associated nerves: the anterior dorsal mesothoracic nerve (ADMN), posterior dorsal mesothoracic nerve (PDMN), and the mesothoracic accessory nerve (MesoAN; (Court et al., 2020). We found 37 neurons on each side. The right side MesoAN is partially severed (**Extended Data Figure 4h**), so some of the MNs could not be reliably traced to a cell body. We only discuss the left side, but all neurons could be paired across the midline by eye. Of the 37 neurons, three innervate the large tergotrochanter “jump” muscle unique to the middle leg and five were eliminated as non-MN efferent neurons. One is the peripherally synapsing interneuron (King and Wyman, 1980), and four are previously undescribed and predicted to be neuromodulatory. We categorized the wing and thorax MNs as indirect, direct, or tension, based on their proposed function (Dickinson and Tu, 1997).

The indirect muscles contain two antagonistic groups: dorsal longitudinal muscles (DLMs) and dorsoventral muscles (DVMs). The six DLM fibers are innervated by five MNs (Ikeda and Koenig, 1988). We differentiated the first four DLMs based on the position of their axons along the anterior-posterior axis; DLM 5, which innervates muscle fibers five and six, is identifiable by its large contralateral cell body. It also has the posterior-most axon, so we labeled the DLM MN with the anterior-most axon as DLM 1. We found three MNs in the ADMN that innervate DVM 1, two MNs in the MesoAN that innervate DVM 2, and two MNs in the PDMN that innervate DVM 3, according to the morphologies and nomenclature established in *Calliphora* (Schlurmann and Hausen, 2007).

Unlike in the leg, direct wing muscles are innervated by a single MN (Heide, 1983). We relied on two papers to identify most of the direct MNs (Trimarchi and Schneiderman, 1994), and identified the rest based on morphology and fasciculation in FANC and consultation with other experts in the field (now published in Ehrhardt et. al., 2023 and Cheong et. al., 2024). The first axillary (i) muscle i1 was identified from Trimarchi and Schneiderman; i2 is identified with lower confidence due to its projection across the midline in FANC, which was not previously observed using light microscopy. It does, however, fasciculate with i1. The third axillary MNs iii1 and iii3 were identified from literature. We determined iii4 based on its similar morphology and fasciculation in the MesoAN. The fourth MN that looks similar to the iii MNs was determined to be iv4 (also known as “hg4”; Cheong et. al., 2024). The fourth axillary MNs iv1 and iv2 were determined based on literature; iv3 was determined based on its similar morphology to iv2. Basalar MNs b1 and b2 were identified from literature. We identified the third basalar MN b3 based on its fasciculation with MNs b1 and b2.

There are two sets of tension muscles, defined by the sclerites they attach to: the tergopleural (tp) muscles and the pleurosternal (ps) muscles. We identified tp1, tp2, and tpn from literature, although the morphologies we observed in FANC were not as similar for tergopleural MNs as other MNs. For the pleurosternal muscles, we found three candidate MNs instead of two. One is predicted to be either neuromodulatory or may also innervate one or both of the pleurosternal muscles, like the tpn MN; it is shown in **Extended Data Figure 5c**.

The main tergotrochanter MN was identified based on previous literature (Bacon and Strausfeld, 1986). Two small MNs whose axons travel alongside the large tergotrochanter axon are predicted to either innervate the large TT muscle or innervate the intracoxal depressor and levator, respectively (muscles 67 and 68, according to Miller (Miller, 1950)).

We did not use the same synapse density-based analysis for wing neurons that we used to validate our identification of leg neurons (**Supplementary Methods Figure 18**). This was primarily because we had higher confidence in comparing FANC MNs with the extensive literature identifying wing MNs than for leg neurons.

**Supplementary Table 2** contains links to view wing motor neurons in neuroglancer.

### Data and Code Availability

The EM dataset and reconstructions are freely available (Phelps et al., 2021).

Code for alignment, segmentation, automatic detection of synapses and cell nuclei, and annotation software (CAVE) can be found via the citations in the Methods section, “EM Dataset Alignment, Segmentation, Annotation.”

The segmentation of the Female Adult Nerve Cord dataset is available by joining the FANC community. Instructions on joining the FANC community can be found at https://github.com/htem/FANC_auto_recon/wiki#collaborative-community. Code for interacting with FANC can be found at https://github.com/htem/FANC_auto_recon.

Code for analysis and figures is available at (github.com/EllenLesser/Azevedo_Lesser_Phelps_Mark_2023/).

---

**Extended Data Figure 1.**
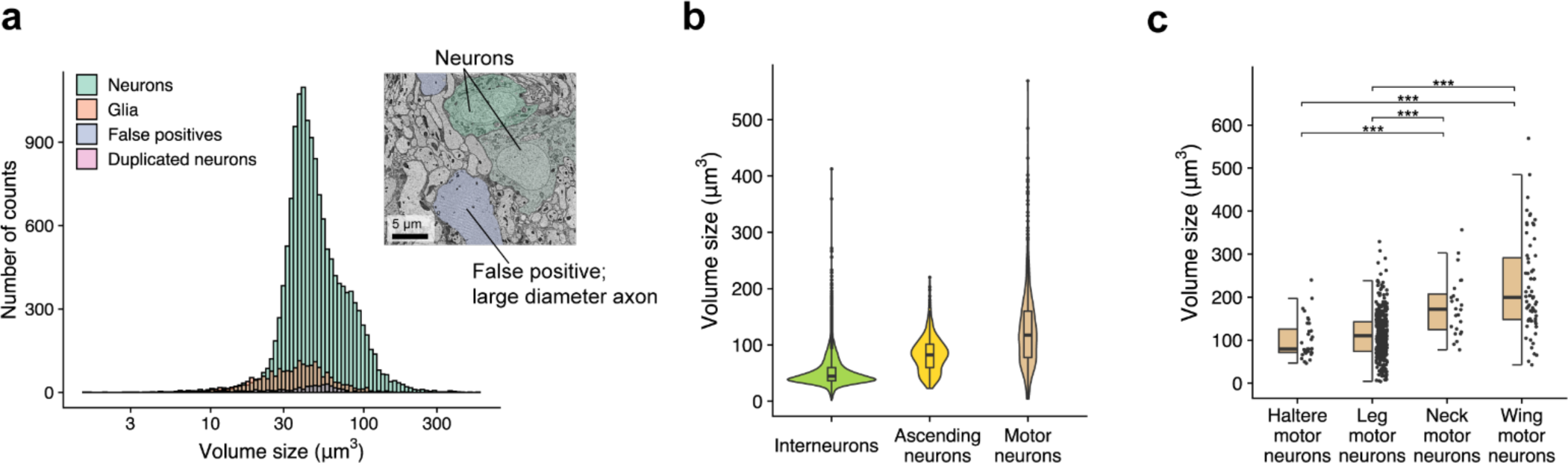
Soma segmentation in FANC. **a,** Size distribution of all 17,076 putative nuclei. We manually inspected each putative nucleus and found 14621 neurons (85.6%), 2030 glia (11.9%), 410 false positives (2.4%), and 15 fragments of neuron nuclei detected twice (Duplicated neurons, 0.1%). Volume size is calculated based on the number of voxels within each detected objects. Inset: example of a large diameter axons that was falsely predicted as a putative nucleus. **b,** Violin plot showing the size distribution of three major neuronal cell types that have cell bodies in VNC: interneurons (n=12,468), ascending neurons (n=1,668), and motor neurons (n=485). (χ2=1760.7, p<0.001, Kruskal-Wallis test.) VNC neurons with arbors projecting to the neck connective were labeled as ascending neurons. Motor neurons include haltere motor neurons (n=32), leg motor neurons (all T1, T2, and T3, n=371), neck motor neurons (n=24), and wing motor neurons (ADMN, PDMN, and MesoAN, n=58). Haltere, leg, and neck motor neurons were identified based on their skeleton nodes previously reported in CATMAID (Phelps et al., 2021). **c,** Comparison of volume size between four motor neurons: haltere motor neurons, leg motor neurons, neck motor neurons, wing motor neurons. (χ2=84.816, p<0.001, Kruskal-Wallis test, ***p<0.001, post-hoc Benjamini–Hochberg procedure-corrected Dunn’s test for multiple comparisons.)

**Extended Data Figure 2.**
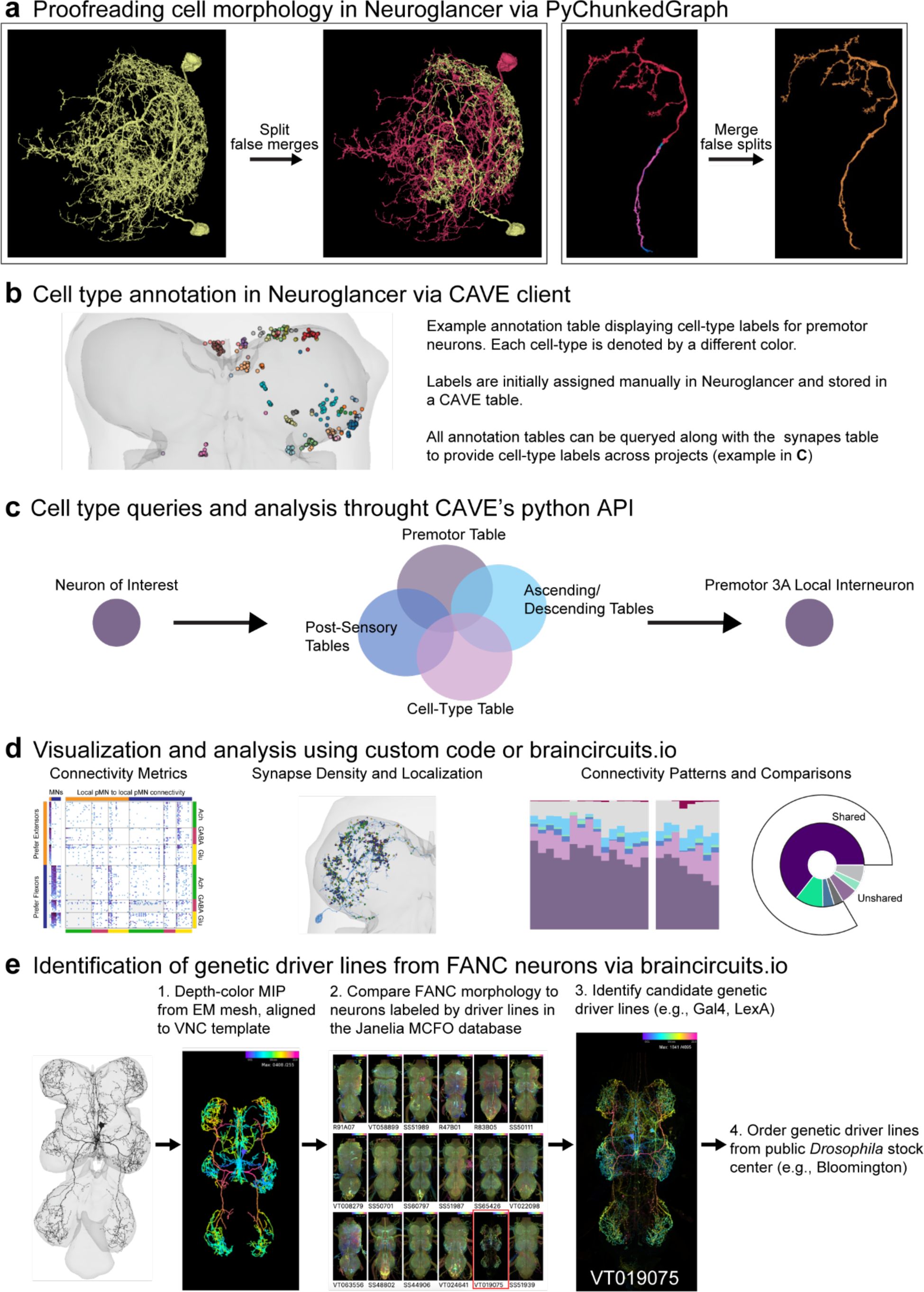
Summary of FANC software tools for cell proofreading: **a,** Proofreading of cell morphology in Neuroglancer via PyChunkedGraph, **b,** cell type annotation, **c,** identification, **d,** cellular and circuit analysis, and **e,** identification of genetic driver lines.

**Extended Data Figure 3.**
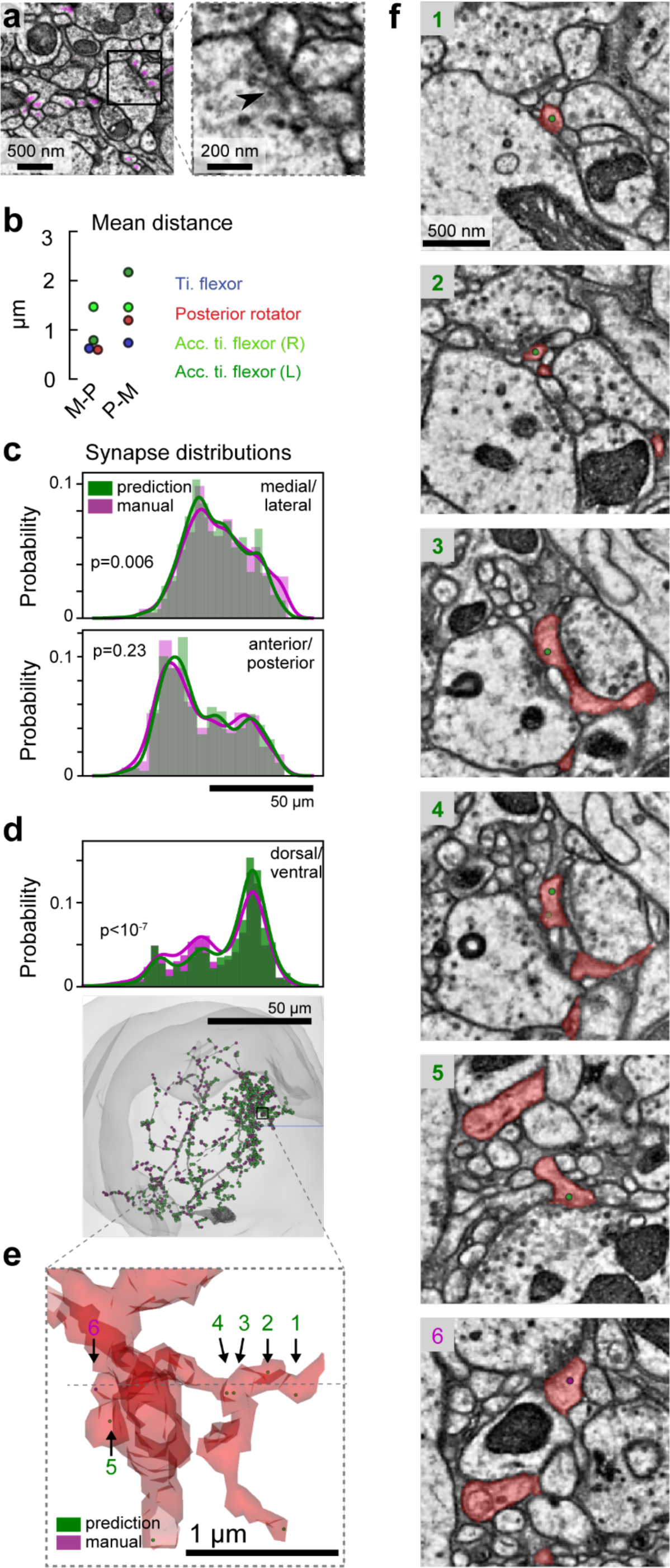
Comparison of automatic synapse prediction with manually annotated ground truth. **a,** Example synapses from the EM volume. Pink blobs indicate predicted postsynaptic sites. The arrowhead in the inset indicates the “T-bar” at the presynaptic site. **b,** Average distance manual to predicted synapses (M-P), and from predicted to manual synapses (P-M) for the four MNs shown in Figure 2. The larger predicted-to-manual distances are consistent with larger numbers of predicted synapses (Figure 2e). **c,** Distributions of manual vs. predicted synapse annotations along the medial/lateral axis (top) and the anterior/posterior axis (bottom) for the posterior rotator MN, shown in Figure 2b. Note, ∼50% more synapses are predicted compared to manual annotations for this neuron (Figure 2E). **d,** Synapse distributions along the dorsal/ventral axis, aligned to the neuron and the synapses below. The distributions are significantly different (Mann-Whitney AUC=0.56, p<10^-7^), with a small increase in the peak of the distribution of predicted synapses. **e,** An example dendrite in the synapse rich region of the dorsal/ventral distribution in **d**. Five synapses are predicted, but only one annotated (#6). **f,** Predicted synapses appear in a region that was not manually traced. We conclude that comparing manual annotation with automatic synapse prediction in these synapse dense reconstructions includes four sources of noise (Schneider-Mizell et al. 2016): 1) completeness of manual tracing of fine branches, 2) completeness of manual synapse annotation, 3) completeness/accuracy of segmentation, and 4) accuracy of synapse prediction. Here, we have ensured that all the manually traced dendrites are proofread and reattached to the segmented reconstructed, thus limiting the third source of noise. Thus, we reiterate our statement in the text to say that the appropriate level of proofreading is likely dictated by the degree to which conclusions based on connectomes would be affected by those sources of noise. In Figure 2f, we show that for the four neurons we examine here, most of the synapses are from a small number of partners, and those partners are largely correctly identified from both the manual and predicted synapses.

**Extended Data Figure 4.**
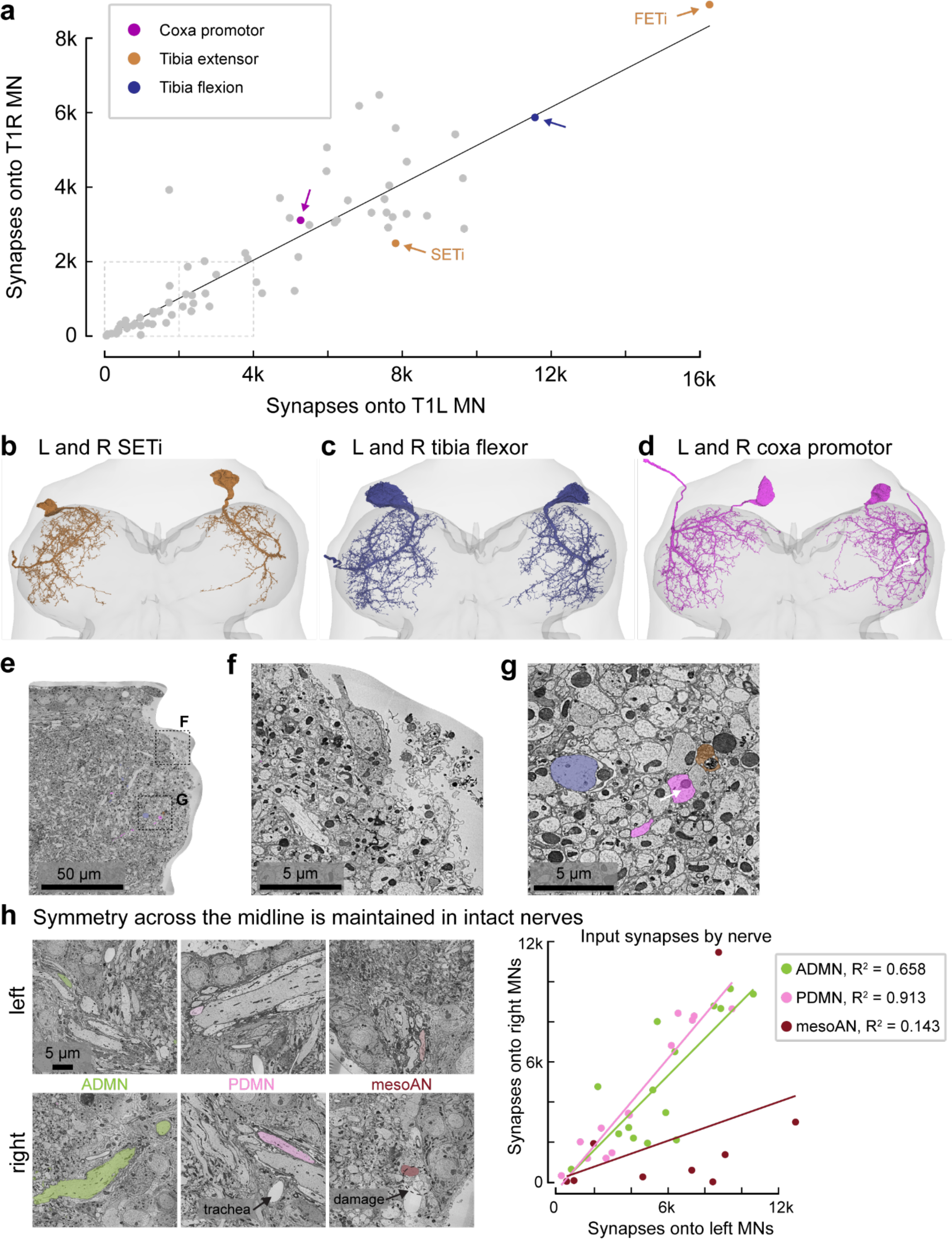
MNs in T1R have half as many input sites as MNs in T1L, likely due to rough dissection of the right T1 leg nerves. **a,** The number of synapses onto the right T1 MN (y-axis) vs. onto the paired left T1 MN (x-axis). Colors indicate MN pairs in **b-d**. The slope of the relationship is 0.51, with a Pearson’s correlation coefficient of 0.89, p<10^-22^. **b,** Left and right SETi MNs. The right T1 neurons tend to appear smoother, with fewer fine twigs. **c,** Left and right main tibia flexor MNs (Fast flexor, (Azevedo et al., 2020). **d,** Left and right pleural coxa promotor MNs. Even though the axons exit the PrDN, rather than the ProLN, many of the dendrites of the right T1 neuron run through the damaged regions. Blebby boutons can be seen (white arrow). **e,** EM image of the damaged area of right T1. **f,** Magnified view of the damaged area. **g,** Magnified view of branches of MNs (colors) near the damaged area, including a bleb (arrow) in the pleural promotor MN (magenta). The bleb diameter is on the order of the primary neurite of the largest cell in the T1 neuromeres (blue). **h,** For comparison, the right side mesoAN is clearly damaged while the right side ADMN and PDMN are intact. Left and right pairs of wing MNs in the ADMN and PDMN have similar numbers of postsynaptic sites, but the right side mesoAN MNs have fewer synapses than the left side mesoAN MNs.

**Extended Data Figure 5.**
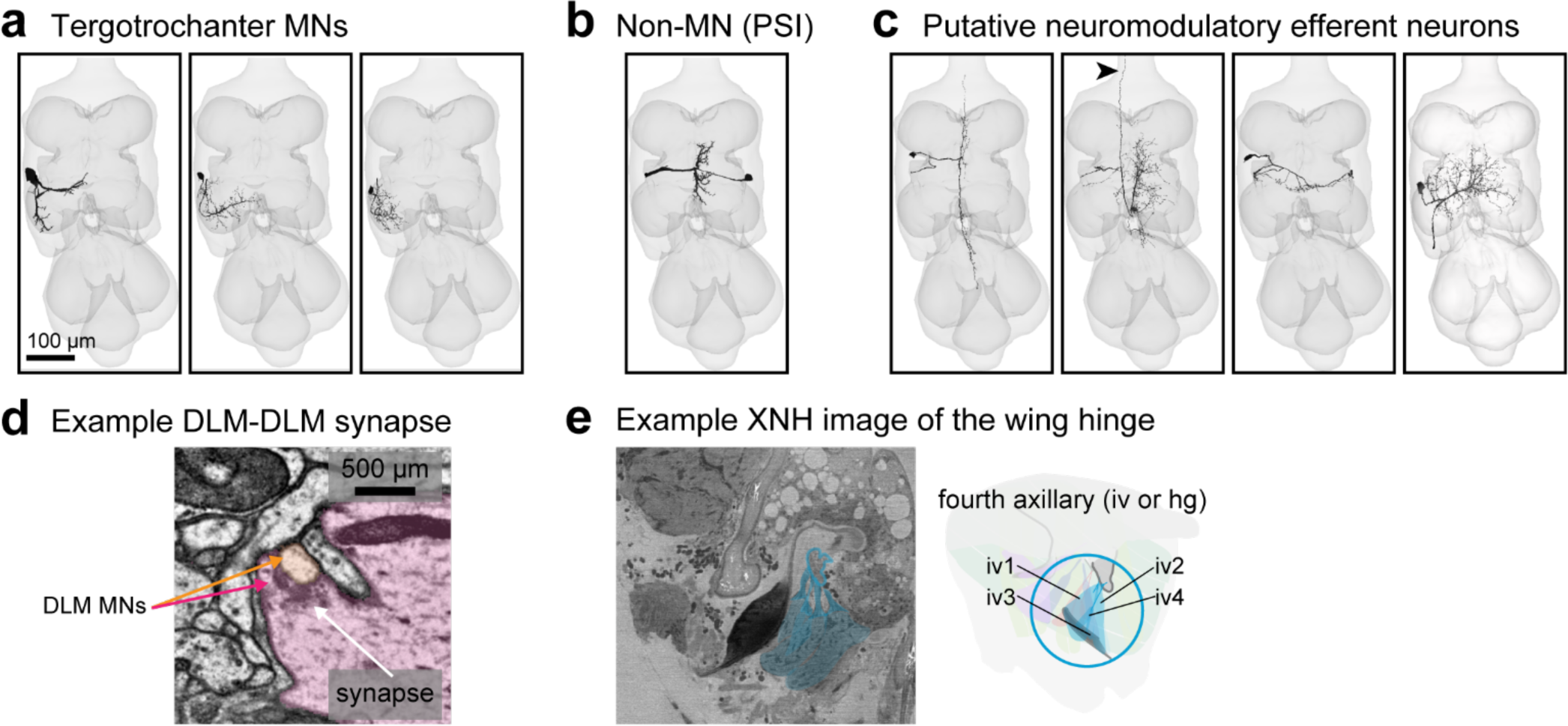
FANC efferent neurons with axons in the ADMN or PDMN that are not wing MNs. **a,** MNs that innervate the T2 tergotrochanter leg muscle send axons through the PDMN. The two small neurons have not been identified previously. We identify them as TT MNs here based on their fasciculation with the main TT MN. We predict that they also innervate the main TT muscle, or the intracoxal depressor and levator, respectively (muscles 67 and 68 according to (Miller, A., 1950). **b,** The peripherally synapsing interneuron (PSI) sends an axon into the PDMN and synapses onto DLM axons but does not innervate muscles (King and Wyman, 1980). **c,** Four other unidentified neurons have axons in the ADMN or PDMN. Their dendrites are thinner than any other motor neurons, and not like anything previously shown using light microscopy. One has an ascending process (indicated with an arrow), and its projection into PDMN does not travel to the end of the dissection so it is not an MN. One (right-most) shares a majority of its input with pleurosternal MNs, and is likely neuromodulatory or may play a similar role to the tpn MN. **d,** DLM MNs are the only MNs we observed with output synapses, and they synapse onto each other. **e,** Recently collected XNH images of the wing and wing hinge were used to help inform the anatomical cartoon schematics.

**Extended Data Figure 6.**
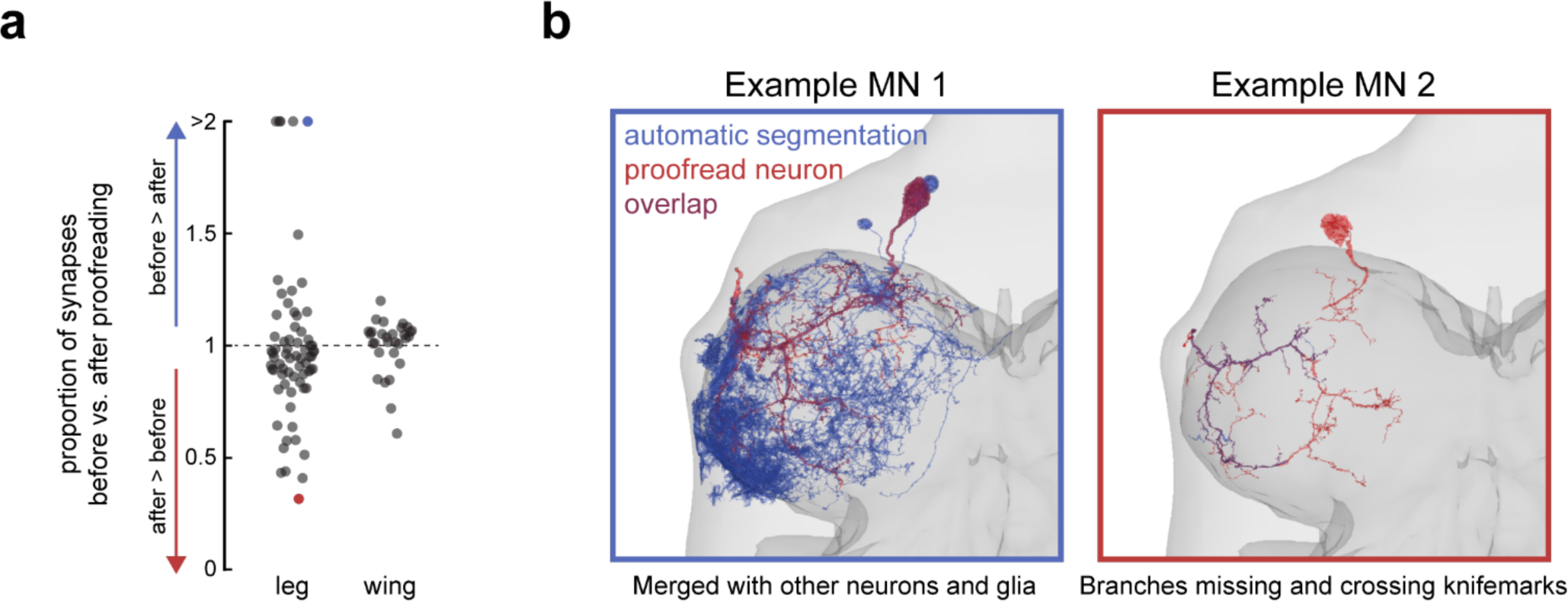
Proofreading of motor neurons. **a,** Overview of how proofreading affected the number of input synapses to each MN. Many of the MN meshes changed substantially, some increased in size as objects were merged (# of synapses before vs. after <1), and some decreased in size as objects were split (# of synapses before vs. after >1). Leg MNs underwent larger changes than wing MNs, relative to starting volume. **b,** Example leg MNs before and after proofreading. Left) An example MN that was initially merged with glia and other neurons (blue, extra somas are visible). Right) An example MN that required branches to be merged across knife marks.

**Supplementary Table 1.** FANC links to leg motor neurons.

| **Nerve** | **Segment** | **Function** | **Muscle** | **MNs** |  |
| --- | --- | --- | --- | --- | --- |
| DProN | thorax | swing | tergopleural promotor, pleural promotor | 4 | link |
| VProN | thorax | swing | sternal anterior rotator | 2 | link |
| ProAN | thorax | swing | sternal adductor | 1 | link |
| ProAN | thorax | stance | pleural remotor and abductor | 2 | link |
| ProAN | thorax | stance | sternal posterior rotator | 4 | link |
| Ventral | thorax | extend | tergotrochanter | 4 | link |
| Ventral | thorax | extend | sternotrochanter extensor | 2 | link |
| ProLN | coxa | extend | trochanter extensor | 2 | link |
| ProAN, VProN | coxa | flex | trochanter flexor | 8 | link |
| ProLN | coxa | flex | accessory trochanter flexor | 3 | link |
| ProLN | trochanter | reductor | femur reductor | 6 | link |
| ProLN | femur | extend | tibia extensor | 2 | link |
| ProLN | femur | flex | main tibia flexor | 5 | link |
| ProLN | femur | flex | accessory tibia flexor | 10 | link |
| ProLN | tibia | levate | tarsus levator | 1 | link |
| ProLN | tibia | depress | tarsus depressor | 4 | link |
| ProLN | tibia | depress | retro tarsus depressor | 1 | link |
| ProLN | femur | claw | ltm femur | 2 | link |
| ProLN | tibia | claw | ltm tibia | 2 | link |
| ProLN |  | claw | ltm uncertain | 2 | link |
| ProLN | femur and tibia | claw | ltm dipalpha | 2 | link |

**Supplementary Table 2.** FANC Links to wing motor neurons.

| **Nerve** | **Muscle** | **Insertion** | **MNs** |  |
| --- | --- | --- | --- | --- |
| PDMN | dorsal longitudinal | thorax | 5 | link |
| ADMN | dorsoventral 1 | thorax | 3 | link |
| mesoAN | dorsoventral 2 | thorax | 2 | link |
| PDMN | dorsoventral 3 | thorax | 2 | link |
| ADMN | i1 | first axillary | 1 | link |
| ADMN | i2 | first axillary | 1 | link |
| mesoAN | iii1 | third axillary | 1 | link |
| mesoAN | iii2 | third axillary | 1 | link |
| mesoAN | iii3 | third axillary | 1 | link |
| ADMN | iv1 | fourth axillary | 1 | link |
| ADMN | iv2 | fourth axillary | 1 | link |
| ADMN | iv3 | fourth axillary | 1 | link |
| mesoAN | iv4 | fourth axillary | 1 | link |
| ADMN | b1 | basalar apophysis | 1 | link |
| ADMN | b2 | basalar apophysis | 1 | link |
| ADMN | b3 | basalar apophysis | 1 | link |
| ADMN | tp1 | meso-scutum | 1 | link |
| ADMN | tp2 | meso-scutum | 1 | link |
| ADMN | tp & tp2 | meso-scutum | 1 | link |
| mesoAN | PS1 | pleural apophysis | 1 | link |
| mesoAN | PS2 | pleural apophysis | 1 | link |
| PDMN | middle leg tergotrochanter |  | 3 | link |
| PDMN | peripherally synapsing interneuron |  | 1 | link |

