## Appendix for "Tools for connectomic reconstruction and analysis of a female *Drosophila* ventral nerve cord"

**Supplementary Methods. Identification of leg motor neuron targets**

Here we describe the effort to match leg motor neurons to their targets. We first describe the resources and general methods. We then give an overview of Appendix A., Figures A1-17, the atlas of FANC left T1 MNs, including an explanation for how the figures are laid out. Finally, specific evidence for each neuron or group of neurons is given in the legends for Figures A2-17.

*Anatomy datasets*

Interpreting the connectome requires knowing which MNs control which muscles. We therefore sought to identify the peripheral muscle targets of all MNs innervating the fly's front (T1) leg. To do so, we used three imaging datasets that collectively span the VNC and leg (main Figure 3A).

1) The FANC dataset establishes the number of MNs in the T1 neuropils: 69 T1L, 70 in right T1. It also shows which of four nerves each MN axon exits the neuropil through: the Prothoracic Accessory Nerve (ProAN), the Dorsal Prothoracic Nerve (DProN), the Prothoracic Leg Nerve (ProLN) and the Ventral Prothoracic Nerve (VProN).
2) An X-ray holographic nano-tomographic (XNH) dataset of the fly's front leg (Kuan et al., 2020)(Kuan et al., 2020) shows each of the prothoracic nerves and their branches into the musculature, and shows both where sensory axons join and where motor axons leave the nerve. For many MNs with large axons, we could even trace their axons to their target muscle fibers. We also used the XNH dataset to determine leg muscle fiber origins, insertions and numbers, as well as how the tendons move the leg joints (Figure A1). At the proximal end of the volume, the insertions of the thoracic muscles onto tendons and apodemes that contact the coxa are visible. In most cases, the origins of the thoracic muscle fibers on the thoracic cuticle are not visible. Distally, the dataset ends ~¾ the length of the tibia. Many of the tibia muscle fibers are visible except for a few notable fibers. The tibia-tarsus joint is not visible.
3) We screened a large collection of VNC neurons sparsely labeled with the multi-color Flp-out (MCFO) technique to identify GAL4 driver lines labeling leg MNs (Meissner et al., 2020). We imaged GFP expression of each genetic driver line in the T1 leg to identify the muscle target of each MN axon. We then compared the dendritic morphology of the genetically-labeled MNs to those reconstructed from FANC (Figure 3C).

Past work showed that most leg MNs have clear matches on the left and right sides (Phelps et al., 2021), which we confirmed through the identification exercise. Thus, for Figures A2-17, we attempt to match only the FANC neurons in left T1, but we use MCFO clones in either left T1 or right T1, and assume that they have a contralateral match.

Each anatomical tool has its drawbacks. For one, the EM volume does not show the muscle targets for each neuron. For another, while it is possible to trace large neurons in the XHN volume, it is difficult to resolve thin axons, particularly within the muscle fibers, making it difficult to precisely count the number of neurons targeting each muscle, or the number of muscle fibers contacted by every neuron. A third drawback, we found 196 images of single motor neurons in the Janelia MCFO collection, but the GAL4 driver lines–from which the MCFO clones are generated–are typically not sparse. In cases where a single MN was labeled by the GAL4 line, we could make a direct one-to-one match between dendrite morphology and muscle target. More often, several motor neurons are labeled by a GAL4 line. However, along with evidence from the literature, the tools can together compensate for these and other drawbacks.

*Drosophila leg motor neurons in the literature.*

To confirm our findings in the anatomy datasets, we cross-referenced the following studies that describe aspects of motor neuron morphology and muscle innervation.

Baek and Mann (Baek and Mann, 2009) used the MARCM technique to label individual neuron clones with GFP, to image their dendritic morphology in the VNC, and to image their axons in the leg. Based on their results, they determined which lineages produced neurons targeting specific leg segments, and the birth order of neurons within different motor neuron lineages. The authors generously shared their data to help confirm the MN identification in this study. Brierley et al. (Brierley et al., 2012) used a similar technique and made similar observations. Together, the studies complemented each other to label and image clones of many of the neurons present in FANC.

Subsequent studies have probed the molecular mechanisms of motor neuron identity and muscle targeting. These papers include additional context, as well as images of motor neuron morphology and their muscle targets, which we have found useful in confirming our MN identification (Enriquez et al., 2015; Guan et al., 2022; Venkatasubramanian et al., 2019). Finally, in our own previous work characterizing the electrophysiology, force generation and neural activity of several specific MNs, we filled tibia flexor neurons with neurobiotin or biocytin, allowing us to definitively match dendritic morphology to axon morphology and functional characteristics (Azevedo et al., 2020).

*Drosophila leg musculature in the literature.*

The names of *Drosophila* leg muscles differ across the literature. Miller (Miller, 1950) applied the nomenclature for locust leg muscles (Snodgrass, R.E., 1935) to *Drosophila*, which has largely been adopted. Here, we define and use synonyms for the musculature to help clarify how a muscle actuates its joint (Table A1). In some cases, we have made novel observations from the XNH volume on how a specific muscle actuates a joint, and we offer new names for the muscle as a result.

Soler et al. (Soler et al., 2004) used genetic techniques to label muscles and tendons, and established nomenclature for the leg musculature, based on older work by Miller (Miller, 1950). As we have argued previously (Azevedo et al., 2020), we believe Soler et al. misidentified the accessory tibia flexors as a tibia reductor muscle, possibly a misreading of earlier work. They did not identify the thoracic muscles, so we have relied on the work by Miller for their names and suspected function.

As a specific example, the tibia levator muscle (Snodgrass, 1935), a.k.a. “tilm” (Soler et al., 2004), refers to the muscle that extends the tibia to “lift” it off the ground. We find the term “levator” unsatisfying for several reasons. First, the terms “levator” and “depressor” are not commonly applied to limbed vertebrates in modern literature, whereas “extensor” and “flexor” are common terms. Second, the action that levator (or depressor) muscles have on a joint is not always the same: the trochanter levator *flexes* the coxa-trochanter/femur joint, whereas the tibia levator *extends* the femur-tibia joint. Third, the well-studied FETi and SETi MNs extend the tibia, so it is simpler to refer to both the extensor muscle and to the extensor MNs. Thus, we call this muscle the tibia extensor muscle in the main text and figures.

We retain the terms “levator” vs. “depressor” for the muscles that “lift” or “push down” the tarsus. When the fly is standing, the tarsus bends back towards the tibia and the tibia-tarsus joint flexes. The depressor muscle causes the tarsus to extend, to push the fly off the substrate, while the levator muscle appears to flex the tibia-tarsus joint further. We note that the terms levator vs depressor imply the animal is standing upright with respect to gravity, and flies often hang from surfaces and walls.

**Table A1.** Muscle nomenclature across the literature.

| Updated muscle name | Action | Atlas Figure, Appendix A | Snodgrass, 1935 | Miller, 1950 | Soler et al., 2004 |
|---|---|---|---|---|---|
| Tergopleural promotor | Promote (move anteriorly) the coxa | Figure A2 | | 28 | |
| Pleural promotor | Promote (move anteriorly) the coxa | Figure A2 | | 30 | |
| Pleural remotor and abductor | Remote (move posteriorly) and abduct (move laterally) the coxa | Figure A4 | | 29 | |

| | | | | | |
|---|---|---|---|---|---|
| Sternal anterior rotator | Anterior movement of coxa | Figure A3 | | 31 | |
| Sternal posterior rotator | Posterior movement of coxa | Figure A4 | | 32 | |
| Sternal adductor | Adduct (move medially) the coxa | Figure A3 | | 33 | |
| Tergotrochanter extensor | Extend the coxa-trochanter joint | Figure A5 | P | Extracoxal trochanteral depressor | |
| Sternotrochanter extensor | Extend the coxa-trochanter joint | Figure A6 | | Extracoxal trochanteral depressor | |
| Trochanter extensor | Extend the coxa-trochanter joint | Figure A6 | Trochanter depressor | Trochanter depressor | Trochanter depressor (trlm) |
| Trochanter flexor | Flex the coxa-trochanter joint | Figure A7-8 | Trochanter levator | Trochanter levator | Trochanter levator (trlm) |
| Accessory trochanter flexor | Flex<br>coxa-trochanter joint | Figure A9 | | | Trochanter reductor<br>(trrm) |
| Femur reductor | Unknown | Figure A10 | Femur reductor | Femur reductor | Femur reductor<br>(ferm) |
| | | | | | Femur depressor (fedm) |
| Tibia extensor | Extend the femur-tibia joint | Figure A11 | Tibia levator | Tibia levator | Tibia levator<br>(tilm) |

| | | | | | |
|---|---|---|---|---|---|
| Tibia flexor | Flex the femur-tibia joint | Figure A12 | Tibia levator | Tibia levator | Tibia depressor<br><br>(tidm) |
| Accessory tibia flexor | Flex the femur-tibia joint | Figure A13-14 | Accessory tibia levator | Accessory tibia levator | Tibia reductor (tirm) |
| Tarsus depressor muscle | Extend the tibia-tarsus joint. The joint is flexed when the fly is standing. Extension moves the fly's body away from the substrate. | Figure A17 | Tarsus depressor | Tarsus depressor | Tarsus depressor muscle<br><br>(tadm) |
| Tarsus retro depressor muscles | Muscle fibers originate on tibia cuticle that is distal to their insertion sites on the tarsus depressor tendon. | Figure A17 | Tarsus depressor | Tarsus depressor | Tarsus reductor muscles 1 and 2.<br><br>(tarm 1 and 2) |
| Tarsus levator muscle | Flex the tibia-tarsus joint.<br><br>The joint is flexed when supporting the fly's weight. Flexion brings the fly closer to the substrate. | Figure A17 | Tarsus levator | Tarsus levator | Tarsus levator muscle<br><br>(talm) |
| Long tendon muscle 2 | Located in femur. Pull on the long tendon. | Figure A15-16 | | | Long tendon muscle 2 |
| Long tendon muscle 1 | Located in the tibia. Pull on the long tendon | Figure A15-16 | | | Long tendon muscle 1 |

*Matching motor neuron dendrite morphology across datasets*

To match motor neurons, we relied on expert visual recognition of specific morphological features for each MN, rather than on numerical algorithms like NBLAST (Costa et al., 2016). NBLAST was successfully used previously to match between left and right T1 MNs, and to classify axon bundles (Phelps et al., 2021). We used the bundle identification together with the following distinguishing characteristics to match motor neurons.

**Prothoracic nerves.** The names and abbreviations of the peripheral nerves come from (Court et al., 2020). As reported in Phelps et al (2021), the following number of motor neurons exit through the four prothoracic nerves in the FANC volume:

1) Prothoracic accessory nerve (ProAN) - 12 MNs. The ProAN follows the leg nerve but splits off just after leaving the neuropil.

2) Dorsal prothoracic nerve (DProN) - 4 MNs. The DProN exits the neuropil laterally, more anteriorly and dorsally than the other nerves.
3) Prothoracic leg nerve (ProLN) - 42 MNs.
4) Ventral prothoracic nerve (VProN) - 11 MNs. The VProN exits the neuromere laterally, more anteriorly than the leg nerve, but ventral-posterior to the DProN.

We assumed that a similar number of motor neurons travel along each nerve in the XNH dataset. We could distinguish motor axons from sensory axons in each nerve in the XNH dataset when they could be traced to their target muscles or source sensory organ, respectively. Sensory neurons often have extremely thin axons, making them difficult to trace, but a limited number have larger axons and are traceable.

**Soma location.** Most MN somas are on the anterior cortex of T1 (Figure 4C). Six neurons have cell bodies on the posterior cortex (Figures A4 and A7) and one additional neuron has a cell body on the dorsal cortex (Figure A6). For neurons with anterior somas, we did not assume that the specific location of the soma was a reliable indicator of identity; We and others have found that across different flies, somas of identified motor neurons can be in different locations within the anterior cluster (Azevedo et al., 2020; Baek & Mann 2009).

**Neurite tracts and neurite bundles.** As shown previously, neurons exiting the same nerve could be further classified into specific bundles based on close proximity of the primary neurites, i.e. the branch running between the soma and the axon (Phelps et al., 2021). In some cases, 3D visualization of FANC MNs in neuroglancer revealed sub-bundles. In many cases, these bundles were associated with muscle targets, so once one MN was identified, we could estimate the number of neurons innervating the muscle.

**Dendrite morphological features.** Most MNs have distinctive, identifying projections within the VNC, which could be used to match neurons in FANC to MCFO clones (Enriquez et al., 2015). We rely heavily on these distinguishing features to group motor neurons together to estimate how many neurons share features, and how they differ morphologically, such as whether there is a gradient in soma, primary neurite size, or number and extent of dendrites. Some of these features are subtle and can differ between neurons that target the same muscle. Many of these features appear indistinguishable in 2D projections, but 3D visualization and depth-colored MCFO projections often reveal distinctions.

**Axon pathways and targets.** Each MN axon innervates a stereotyped set of muscle fibers (Venkatasubramanian et al., 2019). Consequently, each axon also leaves the peripheral nerve to enter a muscle at a roughly stereotyped point. We can observe axons leaving the nerve in the XNH volume, allowing us to count the number of neurons we expect to innervate each leg segment. When several axons innervate a given segment or muscle, the axons can have different thicknesses. Because thicker axons tend to come from MNs with larger somas and larger diameter primary neurites (Azevedo et al., 2020), the gradient of axon thickness should correlate with the number of EM-reconstructed neurons and any gradient in their dendritic properties. Ideally, tracing the full axon branching patterns in the XNH dataset would allow us to create an atlas of axon anatomy that we could compare our light-level leg imaging with. Unfortunately, many of the MN axons are too thin to be traced given the ~200 nm resolution of the XNH dataset, so while the neurons we can fully trace give us valuable information, we were only able to reconstruct a subset of the complete population.

By identifying these features across the datasets, we could estimate the numbers of neurons that share features. We could then use the GAL4 line expression to match axon targets to dendritic morphology, as well as rule out possible matches. Finally, we used inference and process of elimination to buttress direct evidence.

*Confocal imaging*

Fly prothoracic (front) legs were immersed in a 4% formaldehyde (PFA) PBS solution for 20 minutes, followed by three rinses in PBS with 0.2% Triton X-100 (PBT). The legs were then incubated in a PBS solution containing 1:50 phalloidin (Alexa-phalloidin-647, Fisher A222287) and the following reagents that improve tissue penetrance: 1% Triton X-100, 0.5% DMSO, 0.05 mg/ml Escin (Sigma-Aldrich, E1378), and 3% normal goat serum. Legs were incubated for one week at 4 °C with occasional rocking. After staining, legs were rinsed 3x with PBS-Tx, 1 rinse with PBS, and were mounted onto slides in Vectashield. To image MNs innervating the coxal muscles in the thorax, the fly was fixed as above, then hemisected along the parasagital plane with a fine razor blade in Tissue-Tek O.C.T. compound (Sakura 4583) frozen for 10 seconds on dry ice. Hemisected thoraces were rinsed 3x in PBT and stained as above.

Mounted legs or thoraxes were imaged on a Confocal Olympus FV1000. At least one image stack of each segment of the leg was acquired. If GFP was expressed in a motor neuron in a particular segment, two image stacks of the segment were acquired. Images are available upon request. Image stacks were processed in FIJI (Rueden et al., 2017).

*Fly strains for genetically labeling motor neurons*

We screened a large collection of VNC neurons sparsely labeled with the multi-color Flp-out (MCFO) technique to identify GAL4 driver or split-GAL4 hemidriver lines labeling leg MNs (Meissner et al., 2020). The confocal images in Figures A2-A17 come from a resulting collection of 75 lines driving expression of GFP expression in the leg (Table A2). An additional 31 lines were imaged that showed no MN expression in the leg. For "Gen1" Janelia GAL4 lines, the genotype was P{<GMR>-GAL4}attP2/P{20XUAS-06XGFP}attP2. For VDRC Vienna Tile DBD hemidrivers, the genotype was TI{2A-p65(AD)::Zip+}VGlut[2A-p65AD]/+; P{y[+t7.7]w[+mC]=<VT>-GAL4.DBD}attP2/P{20XUAS-06XGFP}attP2.

*Blinding*

Experimenters were not blinded to the genotype when acquiring images.

*Randomization*

Imaging experiments were not intentionally randomized, but lines were imaged as they were ordered and crossed, without care for any particular order.

**Table A2.** Motor neurons expression driven by GAL4 lines and split-GAL4 DBD hemidrivers. The 69 leg MNs in T1 in FANC are indicated in the rows. The columns list specific GAL4 lines (Gen1 Janelia GMR lines, e.g. R10B11) and split-GAL4 DBD hemidrivers (VT lines). Numbers and gray scale indicate a confidence heuristic that a specific MN is labeled by the GAL4 reagent.

| Cell_type | R10B11 | VT023820 | VT029517 | VT000353 | VT025963 | R14B04 | R22g01 | VT002042 | VT017399 | VT049481 | VT063540 | VT043166 | VT023820 | VT025963 | VT063626 | VT063540 | VT015822 | R20C08 | VT056372 | R19G01 | VT003238 | VT019916 | VT000816 | R10G10 | R15D08 | VT008452 | VT026387 | VT034804 | VT054805 | VT064565 |
|---|---|---|---|---|---|---|---|---|---|---|---|---|---|---|---|---|---|---|---|---|---|---|---|---|---|---|---|---|---|---|
| tergopleural_promotor_pleural_promotor_miller_28_30 | 0.16 | | | | | | | | | | | | | | | | | | | | | | | | | | | | | |
| tergopleural_promotor_pleural_promotor_miller_28_30 | 0.16 | | | | | | | | | | | | | | | | | | | | | | | | | | | | | |
| tergopleural_promotor_pleural_promotor_miller_28_30 | 0.16 | | | | | | | | | | | | | | | | | | | | | | | | | | | | | |
| tergopleural_promotor_pleural_promotor_miller_28_30 | 0.16 | | | | | | | | | | | | | | | | | | | | | | | | | | | | | |
| sternal_anterior_rotator_in_thorax_miller_31 | 0.16 | | | | | | | | | | | | | | | | | | | | | | | | | | | | | |
| sternal_anterior_rotator_in_thorax_miller_31 | 0.16 | | | | | | | | | | | | | | | | | | | | | | | | | | | | | |
| sternal_adductor_miller_33 | | | | | | | | | | | | | | | | | | | | | | | | | | | | | | |
| sternal_posterior_rotator_miller_32 | | 0.25 | 0.25 | 0.25 | 0.25 | 0.25 | | | | | | | | | | | | | | | | | | | | | | | | |
| sternal_posterior_rotator_miller_32 | | 0.25 | 0.25 | 0.25 | 0.25 | 0.25 | | | | | | | | | | | | | | | | | | | | | | | | |
| sternal_posterior_rotator_miller_32 | | 0.25 | 0.25 | 0.25 | 0.25 | 0.25 | | | | | | | | | | | | | | | | | | | | | | | | |
| sternal_posterior_rotator_miller_32 | | 0.25 | 0.25 | 0.25 | 0.25 | 0.25 | | | | | | | | | | | | | | | | | | | | | | | | |
| sternal_posterior_rotator_miller_29 | | | | | | | | | | | | | | | | | | | | | | | | | | | | | | |
| sternal_posterior_rotator_miller_29 | | | | | | | | | | | | | | | | | | | | | | | | | | | | | | |
| tergotrochanter_extensor | | | | | | | 0.25 | 0.25 | 0.25 | 0.25 | 0.25 | | | | | | | | | | | | | | | | | | | |
| tergotrochanter_extensor | | | | | | | 0.25 | 0.25 | 0.25 | 0.25 | 0.25 | | | | | | | | | | | | | | | | | | | |
| tergotrochanter_extensor | | | | | | | 0.25 | 0.25 | 0.25 | 0.25 | 0.25 | | | | | | | | | | | | | | | | | | | |
| tergotrochanter_extensor | | | | | | | 0.25 | 0.25 | 0.25 | 0.25 | 0.25 | | | | | | | | | | | | | | | | | | | |
| sternotrochanter_extensor | | | | | | | | | | | | | | | | | | | | | | | | | | | | | | |
| sternotrochanter_extensor | | | | | | | | | | | | | | | | | | | | | | | | | | | | | | |
| trochanter_extensor | | | | | | | | | | | | 0.5 | | | | | | | | | | | | | | | | | | |
| trochanter_extensor | | | | | | | | | | | | 0.5 | | | | | | | | | | | | | | | | | | |
| trochanter_flexor (ProAN, posterior soma) | | | | | | | | | | | | | 0.5 | 0.5 | | | | | | | | | | | | | | | | |
| trochanter_flexor (ProAN, posterior soma) | | | | | | | | | | | | | 0.5 | 0.5 | | | | | | | | | | | | | | | | |
| trochanter_flexor (ProAN) | | | | | | | | | 0.33 | 0.33 | | | | | 0.33 | 0.33 | 0.33 | 0.33 | | | | | | | | | | | | |
| trochanter_flexor (ProAN) | | | | | | | | | 0.33 | 0.33 | | | | | 0.33 | 0.33 | 0.33 | 0.33 | | | | | | | | | | | | |
| trochanter_flexor (ProAN) | | | | | | | | | 0.33 | 0.33 | | | | | 0.33 | 0.33 | 0.33 | 0.33 | | | | | | | | | | | | |
| trochanter_flexor (VProN) | | | | | | | | | 0.33 | 0.33 | | | | | 0.33 | 0.33 | 0.33 | 0.33 | | | | | | | | | | | | |
| trochanter_flexor (VProN) | | | | | | | | | 0.33 | 0.33 | | | | | 0.33 | 0.33 | 0.33 | 0.33 | | | | | | | | | | | | |
| trochanter_flexor (VProN) | | | | | | | | | 0.33 | 0.33 | | | | | 0.33 | 0.33 | 0.33 | 0.33 | | | | | | | | | | | | |
| accessory_trochanter_flexor | | | 0.33 | | | | | | | | | | 0.33 | | | | | | 0.33 | 0.33 | 0.33 | 0.33 | 0.33 | | | | | | | |
| accessory_trochanter_flexor | | | 0.33 | | | | | | | | | | 0.33 | | | | | | 0.33 | 0.33 | 0.33 | 0.33 | 0.33 | | | | | | | |
| accessory_trochanter_flexor | | | 0.33 | | | | | | | | | | 0.33 | | | | | | 0.33 | 0.33 | 0.33 | 0.33 | 0.33 | | | | | | | |
| femur_reductor | | | | | | | | | | | | | | | | | | | | | | | | 0.33 | | | 0.33 | | | |
| femur_reductor | | | | | | | | | | | | | | | | | | | | | | | | 0.33 | | | 0.33 | | | |
| femur_reductor | | | | | | | | | | | | | | | | | | | | | | | | 0.33 | | 0.5 | 0.33 | | | |
| femur_reductor | | | | | | | | | | | | | | | | | | | | | | | | | | 0.5 | | 0.5 | | |
| femur_reductor | | | | | | | | | | | | 0.5 | | | | | | | | | | | | | 0.5 | | | 0.5 | 0.5 | 0.5 |
| femur_reductor | | | | | | | | | | | | 0.5 | | | | | | | | | | | | | 0.5 | | | | 0.5 | 0.5 |
| tibia_extensor (SETi) | | | | | | | | | | 1 | | | | | | | 1 | | | | | | | | | | | | | 1 |
| tibia_extensor (FETi) | | | | | | | | | 1 | | | | | | | | | | | | | | | | | | | | | |
| tibia_flexor | | | | | | | | | | | | | | | | | | | | | | | | | | | | | | |
| tibia_flexor | | | | | | | | 0.5 | | 0.5 | | | | | | | | | | | | | | | | | | | | |
| tibia_flexor | | | | | | | | 0.5 | | 0.5 | | | | | | | | 1 | | | | | | | | | | | | |
| tibia_flexor | | | | | | | | | | | | | | | | | | | | | | | | | | | | | | |
| tibia_flexor | | | | | | | | | | | | | | | | | | | | | | | | | | | | | | |
| accessory_tibia_flexor (A, anterior fibers) | | | | | | | | | | | | | | | | | | | | | | | | | | | | 0.25 | | |
| accessory_tibia_flexor (A, anterior fibers) | | | | | | | | | | | | | | | | | | | | | | | | | | | | 0.25 | | |
| accessory_tibia_flexor (A, anterior fibers) | | | | | | | | | | | | | | | | | | | | | | | | | | | | 0.25 | | |
| accessory_tibia_flexor (A, anterior fibers) | | | | | | | | | | | | | | | | | | | | | | | | | | | | 0.25 | | |
| accessory_tibia_flexor (B, anterior fibers) | | | | | | | | | | | | | | | | | | | | | | | | | | | | | | |
| accessory_tibia_flexor (B, anterior fibers) | | | | | | | | | | | | | | | | | | | | | | | | | | | | | | |
| accessory_tibia_flexor (B, anterior fibers) | | | | | | | | | | | | | | | | | | | | | | | | | | | | | | |
| accessory_tibia_flexor (B, anterior fibers) | | | | | | | | | | | | | | | | | | | | | | | | | | | | | | |
| accessory_tibia_flexor (B, anterior fibers) | | | | | | | | | | | | | | | | | | | | | | | | | | | | | | |
| accessory_tibia_flexor (C, posterior fibers) | | | | | | | | | | | | | | | | | | | | | | | | | | | | | | |
| ltm (dip-alpha) | | 0.5 | | | | | | | | | | | | | | | | | | | | | | | | | | | | |
| ltm (dip-alpha) | | 0.5 | | | | | | | | | | | | | | | | | | | | | | | | | | | | |
| ltm (non-dip-alpha) | | | | | | | | | | | | | | | | | | | | | | | 0.5 | | | | | | | |
| ltm (non-dip-alpha) | | | | | | | | | | | | | | | | | | | | | | | 0.5 | | | | | | | |
| ltm (non-dip-alpha, tibia) | | | | | | | | | | | | | | | | | | | | | | | | | | 1 | | | 0.5 | |
| ltm (non-dip-alpha, tibia) | | | | | | | | | | | | | | | | | | | | | | | | | | | | | 0.5 | |
| ltm (non-dip-alpha, femur) | | | | | | | | | | | | | | | | | | 0.5 | | | | | | | | | | | | |
| ltm (non-dip-alpha, femur) | | | | | | | | | | | | | | | | | | 0.5 | | | | | | | | | | | | |
| tarsus_depressor (ventralU) | | 1 | | | | | | | | | | | | | | | | | | | | | | | | | | | | |
| tarsus_depressor (retro-depressor) | | | | | | | | | | | | | | | | | | | 1 | | 0.25 | | | | | | | | | |
| tarsus_depressor (A) | | | | | | | | | | | | | | | | | | | | | 0.25 | | | | | | | | | |
| tarsus_depressor (B) | | | | | | | | | | | | | | | | | | | | | 0.25 | | | | | | | | | |
| tarsus_depressor (B) | | | | | | | | | | | | | | | | | | | | | 0.25 | | | | | | | | | |
| tarsus_levator (C) | | | | | | | | | | | | | | | | | | | | | | | | | | | | | | |

| Cell_type | VT063540 | VT003238 | R81A06 | R33A12 | R22A08 | R81A07 | R33E06 | VT065132 | VT058563 | VT040577 | VT023555 | VT022017 | VT019759 | VT015783 | R78E05 | R73F02 | R73A05 | R54H12 | R35C09 | 31H05 | R24E12 | R24E09 | R22B10 | R21G01 | 81A04 | VT012323 | R22B05 | R22H10 | VT039488 | VT027938 |
|---|---|---|---|---|---|---|---|---|---|---|---|---|---|---|---|---|---|---|---|---|---|---|---|---|---|---|---|---|---|---|
| tergopleural_promotor_pleural_promotor_miller_28_30 | | | | | | | | | | | | | | | | | | | | | | | | | | | | | | |
| tergopleural_promotor_pleural_promotor_miller_28_30 | | | | | | | | | | | | | | | | | | | | | | | | | | | | | | |
| tergopleural_promotor_pleural_promotor_miller_28_30 | | | | | | | | | | | | | | | | | | | | | | | | | | | | | | |
| tergopleural_promotor_pleural_promotor_miller_28_30 | | | | | | | | | | | | | | | | | | | | | | | | | | | | | | |
| sternal_anterior_rotator_in_thorax_miller_31 | | | | | | | | | | | | | | | | | | | | | | | | | | | | | | |
| sternal_anterior_rotator_in_thorax_miller_31 | | | | | | | | | | | | | | | | | | | | | | | | | | | | | | |
| sternal_adductor_miller_33 | | | | | | | | | | | | | | | | | | | | | | | | | | | | | | |
| sternal_posterior_rotator_miller_32 | | | | | | | | | | | | | | | | | | | | | | | | | | | | | | |
| sternal_posterior_rotator_miller_32 | | | | | | | | | | | | | | | | | | | | | | | | | | | | | | |
| sternal_posterior_rotator_miller_32 | | | | | | | | | | | | | | | | | | | | | | | | | | | | | | |
| sternal_posterior_rotator_miller_32 | | | | | | | | | | | | | | | | | | | | | | | | | | | | | | |
| sternal_posterior_rotator_miller_29 | | | | | | | | | | | | | | | | | | | | | | | | | | | | | | |
| sternal_posterior_rotator_miller_29 | | | | | | | | | | | | | | | | | | | | | | | | | | | | | | |
| tergotrochanter_extensor | | | | | | | | | | | | | | | | | | | | | | | | | | | | | | |
| tergotrochanter_extensor | | | | | | | | | | | | | | | | | | | | | | | | | | | | | | |
| tergotrochanter_extensor | | | | | | | | | | | | | | | | | | | | | | | | | | | | | | |
| tergotrochanter_extensor | | | | | | | | | | | | | | | | | | | | | | | | | | | | | | |
| sternotrochanter_extensor | | | | | | | | | | | | | | | | | | | | | | | | | | | | | | |
| sternotrochanter_extensor | | | | | | | | | | | | | | | | | | | | | | | | | | | | | | |
| trochanter_extensor | | | | | | | | | | | | | | | | | | | | | | | | | | | | | | |
| trochanter_extensor | | | | | | | | | | | | | | | | | | | | | | | | | | | | | | |
| trochanter_flexor (ProAN, posterior soma) | | | | | | | | | | | | | | | | | | | | | | | | | | | | | | |
| trochanter_flexor (ProAN, posterior soma) | | | | | | | | | | | | | | | | | | | | | | | | | | | | | | |
| trochanter_flexor (ProAN) | | | | | | | | | | | | | | | | | | | | | | | | | | | | | | |
| trochanter_flexor (ProAN) | | | | | | | | | | | | | | | | | | | | | | | | | | | | | | |
| trochanter_flexor (ProAN) | | | | | | | | | | | | | | | | | | | | | | | | | | | | | | |
| trochanter_flexor (VProN) | | | | | | | | | | | | | | | | | | | | | | | | | | | | | | |
| trochanter_flexor (VProN) | | | | | | | | | | | | | | | | | | | | | | | | | | | | | | |
| trochanter_flexor (VProN) | | | | | | | | | | | | | | | | | | | | | | | | | | | | | | |
| accessory_trochanter_flexor | | | | | | | | | | | | | | | | | | | | | | | | | | | | | | |
| accessory_trochanter_flexor | | | | | | | | | | | | | | | | | | | | | | | | | | | | | | |
| accessory_trochanter_flexor | | | | | | | | | | | | | | | | | | | | | | | | | | | | | | |
| femur_reductor | | | | | | | | | | | | | | | | | | | | | | | | | | | | | | |
| femur_reductor | | | | | | | | | | | | | | | | | | | | | | | | | | | | | | |
| femur_reductor | | | | | | | | | | | | | | | | | | | | | | | | | | | | | | |
| femur_reductor | | | | | | | | | | | | | | | | | | | | | | | | | | | | | | |
| femur_reductor | | | | | | | | | | | | | | | | | | | | | | | | | | | | | | |
| femur_reductor | | | | | | | | | | | | | | | | | | | | | | | | | | | | | | |
| tibia_extensor (SETi) | | | | | | | | | | | | | | | | | | | | | | | | | | | | | | |
| tibia_extensor (FETi) | | | | | | | | | | | | | | | | | | | | | | | | | | | | | | |
| tibia_flexor | | | 1 | | | | | | | | | | | | | | | | | | | | | | | | | | | |
| tibia_flexor | | | | | | | 0.5 | | | | | | | | | | | | | | | | | | | | | | | |
| tibia_flexor | 1 | | | | | | 0.5 | | | | | | | | | | | | | | | | | | | | | | | |
| tibia_flexor | | | | 1 | 1 | | | | | | | | | | | | | | | | | | | | | | | | | |
| tibia_flexor | 1 | | | | | 1 | | | | | | | | | | | | | | | | | | | | | | | | |
| accessory_tibia_flexor (A, anterior fibers) | | | 0.11 | | | | | 0.25 | 0.25 | 0.25 | 0.25 | 0.25 | 0.25 | 0.11 | 0.5 | 1 | | 0.25 | | 0.5 | 0.25 | 0.25 | 0.25 | | 1 | | | | | |
| accessory_tibia_flexor (A, anterior fibers) | | | 0.11 | | | | | 0.25 | 0.25 | 0.25 | 0.25 | 0.25 | 0.25 | 0.11 | 0.5 | | 0.33 | 0.25 | 1 | 0.5 | 0.25 | 0.25 | 0.25 | | 1 | | | | | |
| accessory_tibia_flexor (A, anterior fibers) | | | 0.11 | | | | | 0.25 | 0.25 | 0.25 | 0.25 | 0.25 | 0.25 | 0.11 | 0.5 | | 0.33 | 0.25 | | 0.5 | 0.25 | 0.25 | 0.25 | | 1 | | | | | |
| accessory_tibia_flexor (A, anterior fibers) | | | 0.11 | | | | | 0.25 | 0.25 | 0.25 | 0.25 | 0.25 | 0.25 | 0.11 | 0.5 | | 0.33 | 0.25 | | 0.5 | 0.25 | 0.25 | 0.25 | | 1 | | | | | |
| accessory_tibia_flexor (B, anterior fibers) | | | 0.11 | | | | | | | | | | | 0.11 | | | | 0.2 | | | | | | 0.2 | | | | | | |
| accessory_tibia_flexor (B, anterior fibers) | | | 0.11 | | | | | | | | | | | 0.11 | | | | 0.2 | | | | | | 0.2 | | | | | | |
| accessory_tibia_flexor (B, anterior fibers) | | | 0.11 | | | | | | | | | | | 0.11 | | | | 0.2 | | | | | | 0.2 | | | | | | |
| accessory_tibia_flexor (B, anterior fibers) | | | 0.11 | | | | | | | | | | | 0.11 | | | | 0.2 | | | | | | 0.2 | | | | | | |
| accessory_tibia_flexor (B, anterior fibers) | | | 0.11 | | | | | | | | | | | 0.11 | | | | 0.2 | | | | | | 0.2 | | | | | | |
| accessory_tibia_flexor (C, posterior fibers) | | 1 | 1 | | | | | | | | | | | | | | | | | | | | 0.5 | | | | | | | |
| ltm (dip-alpha) | | | | | | | | | | | | | | | | | | | | | 1 | | | | | 0.5 | | | | |
| ltm (dip-alpha) | | | | | | | | | | | | | | | | | | | | | | 1 | | | | 0.5 | | 1 | | |
| ltm (non-dip-alpha) | | | | | | | | | | | | | | | | | | | | | | | | | | 0.5 | 0.5 | | | |
| ltm (non-dip-alpha) | | | | | | | | | | | | | | | | | | | | | | | | | | 0.5 | 0.5 | | | |
| ltm (non-dip-alpha, tibia) | | | | | | | | | | | | | | 0.5 | | | | | | | | | | | | | | 0.5 | 0.5 | 0.5 |
| ltm (non-dip-alpha, tibia) | | | | | | | | | | | | | | 0.5 | | | | | | | | | | | | | | 0.5 | 0.5 | 0.5 |
| ltm (non-dip-alpha, femur) | | | | | | | | | | | | | | | | | | | | | | | | | 0.5 | | | | | |
| ltm (non-dip-alpha, femur) | | | | | | | | | | | | | | | | | | | | | | | | | 0.5 | | | | | |
| tarsus_depressor (ventralU) | | | | | | | | | | | | | | | | | | | | | | | | | | 1 | | | | |
| tarsus_depressor (retro-depressor) | | | | | | | | | | | | | | | | | | | | | 1 | | | | | | | | | |
| tarsus_depressor (A) | | | | | | | | | | | | | | | | | | | | | | | | | | | | | | |
| tarsus_depressor (B) | | | | | | | | | | | | | | | | | | | | | | | | | | | | | | |
| tarsus_depressor (B) | | | | | | | | | | | | | | | | | | | | | | | | | | | | | | |
| tarsus_levator (C) | | | | | | | | | | | | | | | | | | | | | | | | | | | | | | |

| Cell_type | VT016795 | VT012328 | VT005539 | VT002044 | R76E09 | R38c08 | R32B12 | R18h11 | R38D01 | VT043144 | VT040577 | VT037805 | VT004726 | VT021792 | VT000837 |
|---|---|---|---|---|---|---|---|---|---|---|---|---|---|---|---|
| tergopleural_promotor_pleural_promotor_miller_28_30 | | | | | | | | | | | | | | | |
| tergopleural_promotor_pleural_promotor_miller_28_30 | | | | | | | | | | | | | | | |
| tergopleural_promotor_pleural_promotor_miller_28_30 | | | | | | | | | | | | | | | |
| tergopleural_promotor_pleural_promotor_miller_28_30 | | | | | | | | | | | | | | | |
| sternal_anterior_rotator_in_thorax_miller_31 | | | | | | | | | | | | | | | |
| sternal_anterior_rotator_in_thorax_miller_31 | | | | | | | | | | | | | | | |
| sternal_adductor_miller_33 | | | | | | | | | | | | | | | |
| sternal_posterior_rotator_miller_32 | | | | | | | | | | | | | | | |
| sternal_posterior_rotator_miller_32 | | | | | | | | | | | | | | | |
| sternal_posterior_rotator_miller_32 | | | | | | | | | | | | | | | |
| sternal_posterior_rotator_miller_32 | | | | | | | | | | | | | | | |
| sternal_posterior_rotator_miller_29 | | | | | | | | | | | | | | | |
| sternal_posterior_rotator_miller_29 | | | | | | | | | | | | | | | |
| tergotrochanter_extensor | | | | | | | | | | | | | | | |
| tergotrochanter_extensor | | | | | | | | | | | | | | | |
| tergotrochanter_extensor | | | | | | | | | | | | | | | |
| tergotrochanter_extensor | | | | | | | | | | | | | | | |
| sternotrochanter_extensor | | | | | | | | | | | | | | | |
| sternotrochanter_extensor | | | | | | | | | | | | | | | |
| trochanter_extensor | | | | | | | | | | | | | | | |
| trochanter_extensor | | | | | | | | | | | | | | | |
| trochanter_flexor (ProAN, posterior soma) | | | | | | | | | | | | | | | |
| trochanter_flexor (ProAN, posterior soma) | | | | | | | | | | | | | | | |
| trochanter_flexor (ProAN) | | | | | | | | | | | | | | | |
| trochanter_flexor (ProAN) | | | | | | | | | | | | | | | |
| trochanter_flexor (ProAN) | | | | | | | | | | | | | | | |
| trochanter_flexor (VProN) | | | | | | | | | | | | | | | |
| trochanter_flexor (VProN) | | | | | | | | | | | | | | | |
| trochanter_flexor (VProN) | | | | | | | | | | | | | | | |
| accessory_trochanter_flexor | | | | | | | | | | | | | | | |
| accessory_trochanter_flexor | | | | | | | | | | | | | | | |
| accessory_trochanter_flexor | | | | | | | | | | | | | | | |
| femur_reductor | | | | | | | | | | | | | | | |
| femur_reductor | | | | | | | | | | | | | | | |
| femur_reductor | | | | | | | | | | | | | | | |
| femur_reductor | | | | | | | | | | | | | | | |
| femur_reductor | | | | | | | | | | | | | | | |
| femur_reductor | | | | | | | | | | | | | | | |
| tibia_extensor (SETi) | | | | | | | | | | | | | | | |
| tibia_extensor (FETi) | | | | | | | | | | | | | | | |
| tibia_flexor | | | | | | | | | | | | | | | |
| tibia_flexor | | | | | | | | | | | | | | | |
| tibia_flexor | | | | | | | | | | | | | | | |
| tibia_flexor | | | | | | | | | | | | | | | |
| tibia_flexor | | | | | | | | | | | | | | | |
| accessory_tibia_flexor (A, anterior fibers) | | | | | | | | | | | | | | | |
| accessory_tibia_flexor (A, anterior fibers) | | | | | | | | | | | | | | | |
| accessory_tibia_flexor (A, anterior fibers) | | | | | | | | | | | | | | | |
| accessory_tibia_flexor (A, anterior fibers) | | | | | | | | | | | | | | | |
| accessory_tibia_flexor (B, anterior fibers) | | | | | | | | | | | | | | | |
| accessory_tibia_flexor (B, anterior fibers) | | | | | | | | | | | | | | | |
| accessory_tibia_flexor (B, anterior fibers) | | | | | | | | | | | | | | | |
| accessory_tibia_flexor (B, anterior fibers) | | | | | | | | | | | | | | | |
| accessory_tibia_flexor (B, anterior fibers) | | | | | | | | | | | | | | | |
| accessory_tibia_flexor (C, posterior fibers) | | | | | | | | | | | | | | | |
| ltm (dip-alpha) | | | | | | | | | | | | | | | |
| ltm (dip-alpha) | | | | | | | | | | | | | | | |
| ltm (non-dip-alpha) | | | | | | | | | | | | | | | |
| ltm (non-dip-alpha) | | | | | | | | | | | | | | | |
| ltm (non-dip-alpha, tibia) | 0.5 | 0.5 | | 1 | 0.5 | 1 | | | | | | | | | |
| ltm (non-dip-alpha, tibia) | 0.5 | 0.5 | 1 | | 0.5 | | 1 | | | | | | | | |
| ltm (non-dip-alpha, femur) | | | | | | | | | | | | | | | |
| ltm (non-dip-alpha, femur) | | | | | | | | | | | | | | | |
| tarsus_depressor (ventralU) | | | | | | | | | | | | | 1 | | 1 |
| tarsus_depressor (retro-depressor) | | | | | | | | | 1 | 1 | 1 | 1 | | 1 | 0.5 |
| tarsus_depressor (A) | | | | | | | | | | | | | | 0.25 | |
| tarsus_depressor (B) | | | | | | | | | | | | | | 0.25 | |
| tarsus_depressor (B) | | | | | | | | | | | | | | 0.25 | |
| tarsus_levator (C) | | | | | | | | 1 | | | | | | | |

**Table A3.** GAL4 lines and split-GAL4 DBD hemidrivers in which no GFP expression in motor neurons was observed.

| | | |
|---|---|---|
| R21F07 | R36G02 | VT006406 |
| Dh31-Gal4 | R55D06 | VT009849 |
| R10A12 | R60C09 | VT010277 |
| R23C02 | R71C11 | VT047878 |
| R26A08 | R71D08 | VT038822 |
| R26B11 | R74B11 | VT007182 |
| R26E08 | R82E12 | VT015432 |
| R26H12 | R86F11 | VT022004 |
| R27H01 | R92D04 | VT045148 |
| R33C10 | R92D08 | VT058388 |
| R34G07 | VT033623 | |

*Evidence for muscle target identification and format of Supplementary Atlas Figures*

In Figures A2-17, we go MN-by-MN to explain how we assigned each MN in the left T1 neuromere in FANC to its muscle. In the legends for each Appendix A figure, we interpret the evidence for the match. We also describe the muscle architecture and the points of tendon attachment and mechanisms of moving the joint, in cases where such information is novel for *Drosophila*.

We lay out the Supplementary Figures according to the following general format. For a representative figure, see Figure A10.

- The left most column shows the FANC segmentation of the MNs. We show the MNs first, to underscore the objective of the exercise, i.e., to find the muscle target for a set of FANC MNs and cross them off the list. The grayscale indicates different motor neurons, roughly from smallest to largest in volume, numbered from 1 to 69. The same grayscale is used in Figure 3 when the neurons are displayed together.
- The next column typically shows MCFO clones from the Janelia Neuronbridge MCFO collection that show similar morphology to the FANC neurons. The images are depth-colored maximum intensity projections (MIPs). We aimed to include only images of bright, single clones, but we include less clear images (dim, multiple neurons, etc.) when the morphology is recognizable. Our objective with these images was to show examples of the different morphologies we see in FANC. Ideally, we would find GAL4 lines to label each specific MN, but some of the MCFO clones are likely the same neuron labeled by different GAL4 lines.
- The third column typically shows the leg or body expression of GFP, driven by the GAL4 line that produced the MCFO clone to its left. The GAL4 lines often labeled multiple MNs, such that we observed axons that innervated other muscles, which can be seen in several images. In many cases, we could recognize the other MNs, and often they would appear in a different fluorescence channel or separate MCFO sample. Occasionally, no MCFO image would show the additional MNs. In those cases, we would have to circle back to that GAL4 line once we had more information, in order to determine which muscle the MCFO clone targeted.
- The fourth column of the atlas figures shows a schematic of the target muscle and the leg segments. The schematic is taken from the annotated muscle fibers in the XNH volume. The annotated muscle fibers are shown next to the schematic. The muscles can have additional substructure, which we illustrate with the schematic. For instance, the FETi MN innervates the more proximal fibers of the tibia extensor muscle, while the SETi innervates the more distal fibers (Figure A11).
- In the rightmost column, we include images from the XNH volume that corroborate the number of neurons that innervate a particular muscle. For instance, we show cross sections of nerves that we can follow to the muscles, and we account for the other neurons in those nerves. Or, we show images of countable neurons leaving the ProLN leg nerve. In cases where we could trace the motor neuron through the muscle fibers, we include the traced skeleton and its muscle fibers.
- Finally, in some figures, we include novel information about muscles and tendons and how they attach to cuticle and actuate a joint.

To fully appreciate the three-dimensional details of leg biomechanics, the XNH volume of the leg, including annotations, is a public resource available at https://www.lee.hms.harvard.edu/resources.

**Table A4.** Muscle targets of FANC motor neurons, based on identifying features across anatomy datasets.

| Muscle | Atlas Figure, Appendix A | FANC MNs; | Axons, nerve in XNH | Clearest evidence supporting identification |
|---|---|---|---|---|
| Tergopleural promotor, Pleural promotor | Figure A2 | 4 | 1-4, DProN | The MNs in the DProN and only those MNs enter the promotor muscles. Total # of axons in XNH unclear, the nerve travels in a region of the volume that is reconstructed from lower resolution tomographs. |
| Pleural remotor and abductor | Figure A4 | 2 | 2, ProAN | |
| Sternal anterior rotator | Figure A3 | 2 | 2, VProN | |
| Sternal posterior rotator | Figure A4 | 4 | 4, ProAN | MCFO confirms neurons with posterior somas innervate the posterior rotator |
| Sternal adductor | Figure A3 | 1 | 1<br>ProAN | A single axon leaves the ProAN and enters the muscle |
| Tergotrochanter extensor | Figure A5 | 4 | 4, VProN | Thorax imaging of GAL4 lines, MCFO clones with distinctive L-shaped morphology and VProN axons |
| Sternotrochanter extensor | Figure A6 | 2 | 2,<br>VProN | Axons in VProN, dendritic morphology is very similar to trochanter extensor MNs. 2 axons from VProN innervate the muscle in XNH |
| Trochanter extensor | Figure A6 | 2 | 2,<br>ProLN | FANC MNs are tightly bundled, despite somas in different places. Baek and Mann data show that a neuron with a dorsal soma innervates the tr. extensor muscle (LinJ) |
| Trochanter flexor | Figure A7-8 | 8 | 8,<br>ProAN (5)<br>VProN (3) | MCFO clones and GAL4 line imaging confirm that MNs with posterior somas do innervate the proximal fibers of the tr. flexor (Figure A7), which accounts for the remaining 2 MNs with small, posterior somas (of 6 total, with 4 going to the sternal posterior rotator).<br><br>In XNH, 5 axons innervate the tr. flexor from the ProAN, 3 from the VProN.<br><br>FANC morphology of 3 MNs bundled in ProAN and 3 MNs bundled in VProN, which all resemble the morphology characterized in Enriquez et al. 2015. |
| Accessory trochanter flexor | Figure A9 | 3 | ? | Primary neurites bundled with other Tr. flexors, but travel in the ProLN. MCFO clones with similar morphology->GAL4 lines show ProLN axons innervate the acc. tr. flexor muscle. |
| Femur reductor | Figure A10 | 6, w 2 very small | 6, w/ 2 very small<br>ProLN | Enriquez et al. (2015) investigated the genetic determinants of the characteristic morphology of femur reductors neurons. |
| Tibia extensor | Figure A11 | 2 | 2<br>ProLN | 2 neurons exit the ProLN together, pass through fascial membrane. |
| Tibia flexor | Figure A12 | 5 | 4<br>ProLN, possibly a 5th with the fast flexor. | Well-known morphology from Azevedo et al. 2020 and others. Neurons with the visually most elaborate dendrites, entering all regions of the neuropil. |
| Accessory tibia flexor | Figure A13-14 | 10 | 5 anterior<br>5 posterior<br>ProLN | Baek and Mann reported 9 acc. tibia flexor neurons, based on axon morphology. We found 10 neurons with characteristic morphology in FANC that resembles the morphology of the slow tibia flexor in Azevedo et al. 2020 |

| | | | | |
|---|---|---|---|---|
| Tarsus depressor muscle | Figure A17 | 6 | Uncertain, cut off | Early born LinA neuron has a characteristic ventral u-shaped projection and targets the Tarsus depressor muscle. Other MNs share some morphological features except for ventral u-shape. |
| Tarsus retro depressor muscles | Figure A17 | | Uncertain, cut off | GAL4 lines label a neuron with a medial projection that targets the fibers that originate on the distal tibia cuticle. |
| Tarsus levator muscle | Figure A17 | | Tarsus levator ProLN | GAL4 line and MCFO image of a neuron with a medial projection similar to FETi and SETi. MN in left T1 does not have this medial projection, but its pair in right T1 does. |
| Long tendon muscle 2 | Figure A15-16 | 4 | 4<br>ProLN | GAL4 line and MCFO with posterior dendrites, as opposed to more anterior dendrite, which is characteristic of ltm1.<br>GAL4 line and MCFO image of smaller morphology and axon targeting ltm2. |
| Long tendon muscle 1 | Figure A15-16 | 4 | 4<br>ProLN | GAL4 line and MCFO images of cells with an anterior medial dendrite, which is characteristic of ltm1.<br>GAL4 line and MCFO image of smaller morphology and axon targeting ltm1. |

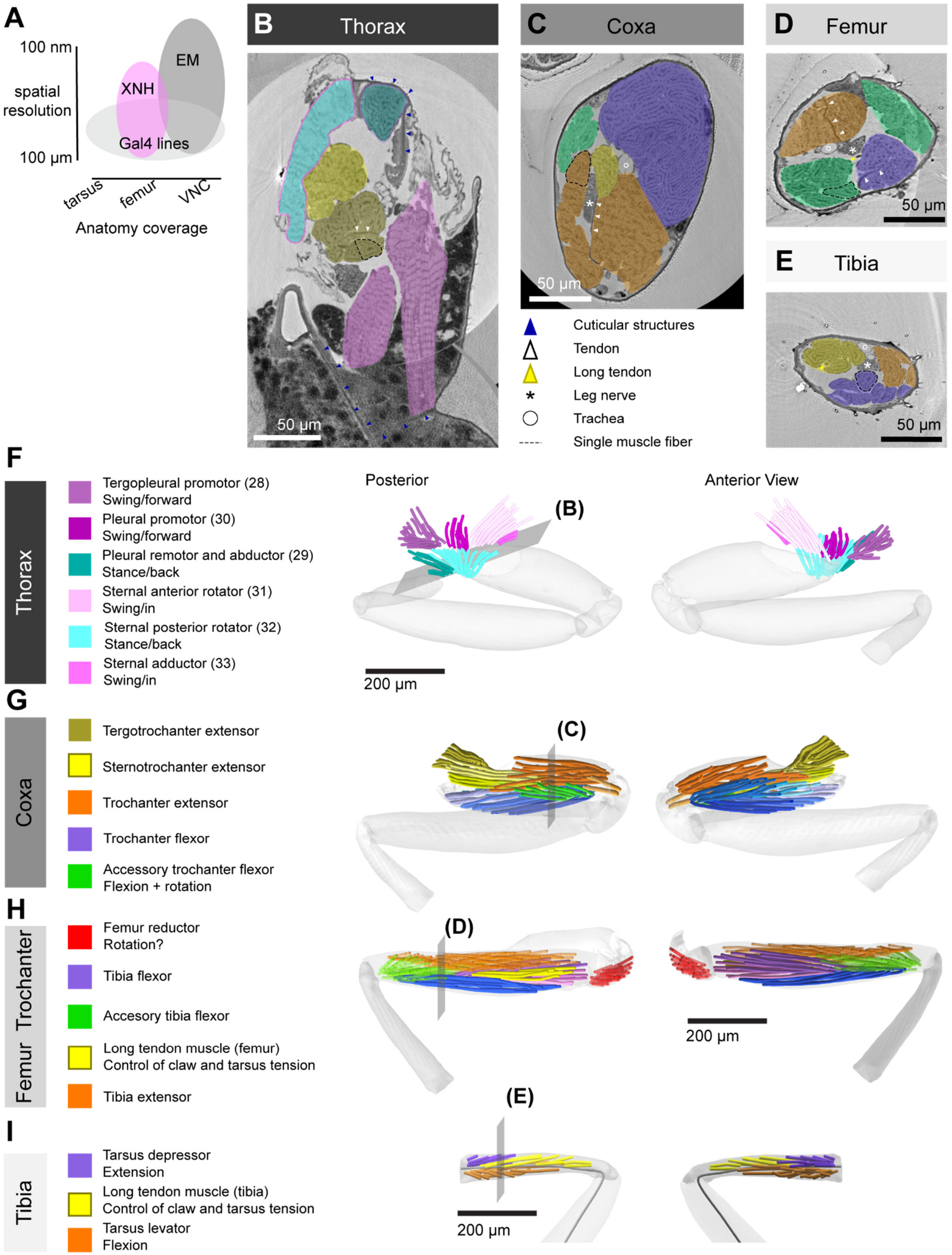
A
100 nm
spatial resolution
100 µm
XNH
EM
Gal4 lines
tarsus
femur
VNC
Anatomy coverage
B Thorax
50 µm
C Coxa
50 µm
D Femur
50 µm
E Tibia
50 µm
Cuticular structures
Tendon
Long tendon
Leg nerve
Trachea
Single muscle fiber
F
Thorax
Tergopleural promotor (28) Swing/forward
Pleural promotor (30) Swing/forward
Pleural remotor and abductor (29) Stance/back
Sternal anterior rotator (31) Swing/in
Sternal posterior rotator (32) Stance/back
Sternal adductor (33) Swing/in
Posterior
Anterior View
(B)
200 µm
G
Coxa
Tergotrochanter extensor
Sternotrochanter extensor
Trochanter extensor
Trochanter flexor
Accessory trochanter flexor Flexion + rotation
(C)
H
Femur Trochanter
Femur reductor Rotation?
Tibia flexor
Accesory tibia flexor
Long tendon muscle (femur) Control of claw and tarsus tension
Tibia extensor
(D)
200 µm
I
Tibia
Tarsus depressor Extension
Long tendon muscle (tibia) Control of claw and tarsus tension
Tarsus levator Flexion
(E)
200 µm

**Figure A1. Overview of leg musculature and the X-ray holographic nano-tomography dataset.** (**A**) The XNH volume has the resolution to view individual axons, and the spatial coverage to view most of the leg as well as the nerves entering the VNC (Kuan et al., 2020). (**B**) Section through the muscles in the thorax that actuate the coxa. Image plane is indicated in F, muscle colors are indicated in F and G. (**C**) Section through the muscles in the coxa that actuate the trochanter/femur. Image plane is indicated in G, muscle colors are indicated in G. (**D**) Section through the muscles in the femur that actuate the tibia. Image plane and muscle color are indicated in H. (**E**) Section through the muscles in the tibia that actuate the tarsus. Image plane and muscle color are indicated in I. (**F**) Annotated muscle fibers that actuate the thorax-coxa joint. Cyan muscle fibers move the coxa anteriorly and medially, magenta muscles move the coxa posteriorly and laterally. (**G**) Annotated muscle fibers that actuate the coxa-trochanter joint. The trochanter-femur joint is thought to be fused to the femur, and these muscles effectively actuate the trochanter/femur. The tergotrochanter, sternotrochanter and trochanter extensor muscle fibers all insert on the trochanter extensor tendon, though they originate from the tergum, the sternum, and the coxa, respectively. (**H**) Annotated muscle fibers in the trochanter and femur. The tibia flexor, accessory tibia flexor, and tibia extensor muscle actuate the femur-tibia joint. The different shades of blue/purple for tibia flexor fibers indicate fiber innervated by each MNs. The long tendon muscle fibers insert on the long tendon (see D and E). The function of the femur reductor muscle is unknown. (**I**) Annotated muscle fibers that actuate the femur-tibia joint. The XNH volume stops ~¾ the length of the tibia. Missing from the XNH are the tibia retro depressor muscle fibers (Table A1).

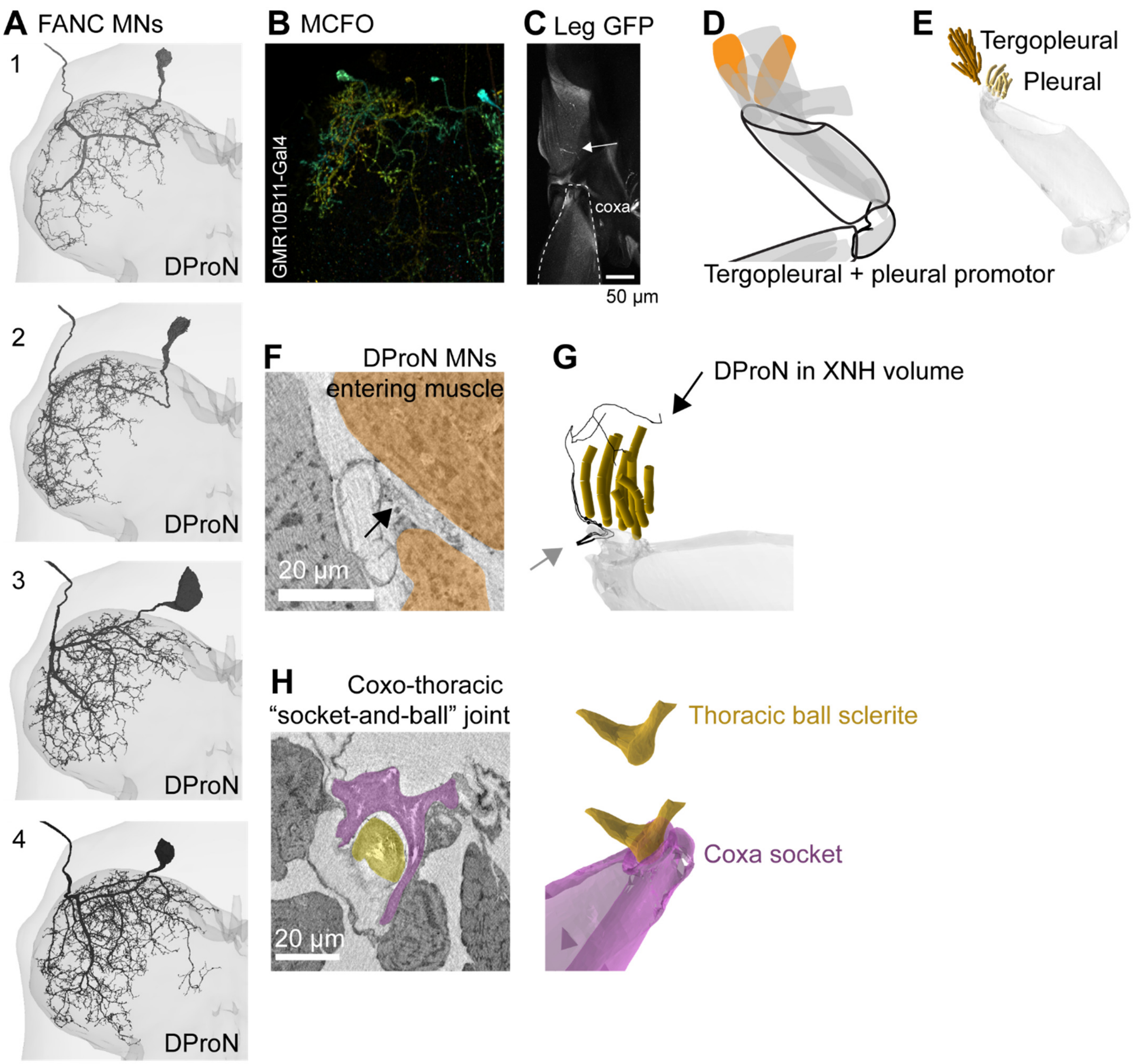

**Figure A2. MNs 1-4 innervate the tergopleural promotor and the pleural promotor muscles in the thorax.** (**A**) FANC reconstruction of MNs 1-4. Grayscale indicates different MNs as in Figure 4. MNs 1-4 exit the prothoracic dorsal nerve (DProN). MNs are numbered as they are ordered in Figure 2. (**B**) Depth-colored maximum intensity projection (MIP) of a multi-colored flip-out (MCFO) clone (GMR10B11-Gal4). Green colors indicate ventral, redder colors indicate dorsal objects. Other neurons are labeled in the image, but a single left T1 MN with an axon exiting the DProN is visible. (**C**) Expression of GFP (grayscale) in the thorax showing targeting of muscle fibers inserting onto the medial-anterior aspect of the coxa (dashed line outline). (**D**) Schematic of the tergopleural and pleural promotors within the leg musculature. (**E**) Muscle fibers annotated in the XNH volume. Muscle contraction would tend to pull the coxa forward (promotion), relative to the joint. (**F**) The DProN (arrow) weaves through the muscle fibers of the tergopleural promotor and the pleural promotor muscles (orange). (**G**) The same DProN carries sensory information from coxal hairplate 3 (gray arrow, Kuan et al. 2020), which can be identified in FANC, supporting the claim that the DProN is correctly identified in the XNH volume. In summary, four MNs exit the DProN and likely innervate these two muscles. In a companion paper, we show that all four receive input from common presynaptic partners (Lesser, Azevedo et al. 2023). However, we are uncertain about the exact target for each MN. (**H**) The thorax-coxa joint resembles a ball and socket joint, where the socket is on the proximal coxa, reversed from the mammalian shoulder or hip construction. In the XNH volume, a small sclerite can be seen emanating from the thoracic cuticle (gold), and a cup shaped-portion of the coxa (purple) appears to hang from the thoracic sclerite. Other insects, as well as crustaceans appear to have a different coxal-trochanter structure that limits the degrees of freedom of the joint (Frantsevich and Wang, 2009; Hessler, 1982).

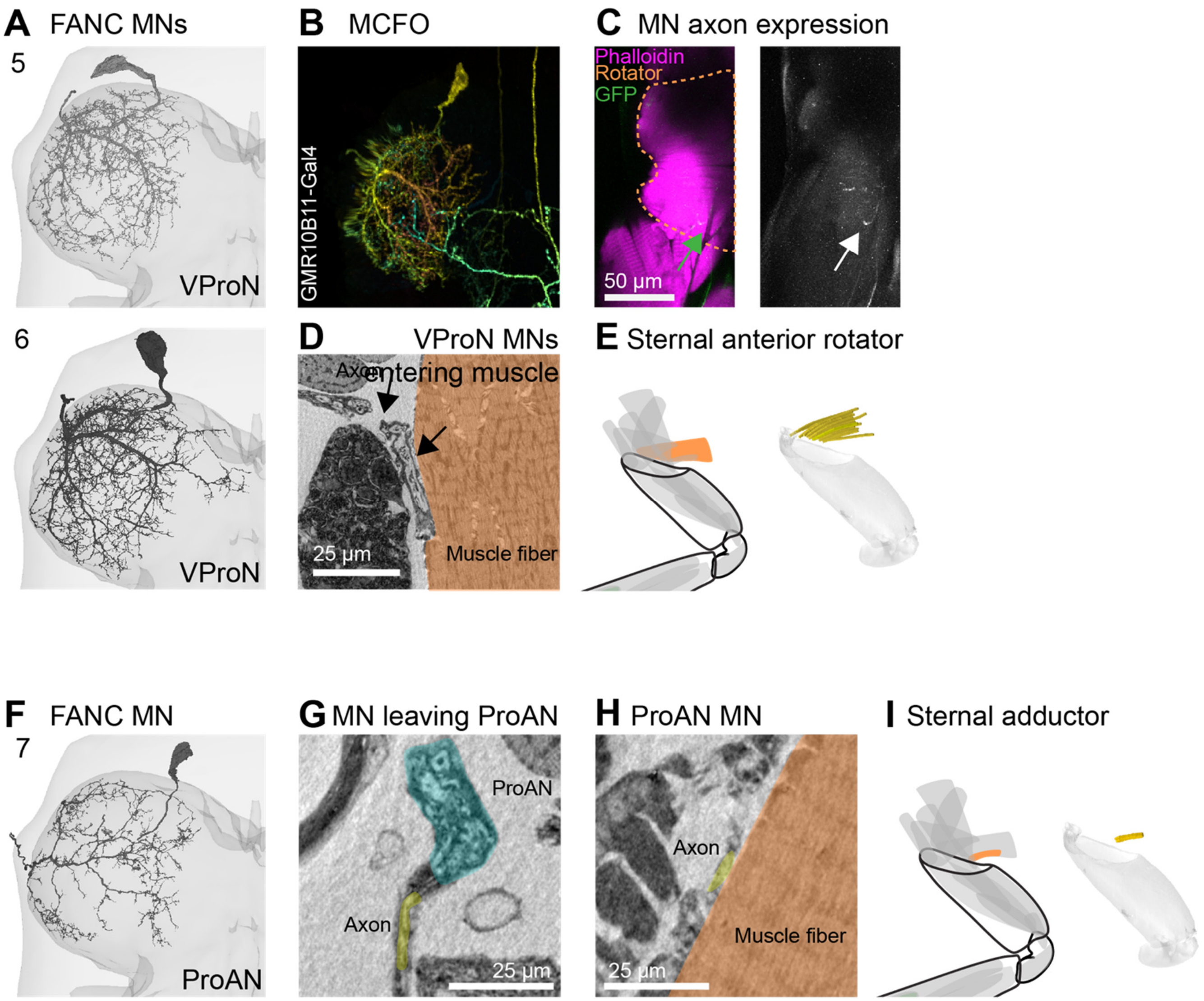

**Figure A3. MNs 5-6 innervate the sternal anterior rotator muscle. MN 7 innervates the sternal adductor muscle.** (**A**) MNs 5-6 exit the ventral prothoracic nerve (VProN). (**B**) Depth-colored MCFO clone (GMR10B11-Gal4). Other neurons are labeled in the image, but a single left T1 MN with an axon exiting the DProN is visible. (**C**) Left, GFP expression in the thorax, with phalloidin counterstain (magenta). An axon enters the sternal anterior rotator muscle (orange dashed line). Right, single channel in grayscale showing GFP expression in the thorax. Note, GMR10B11-Gal4 also labels a tergopleural promotor MN (Figure A2), and this GFP image shows a separate plane of the same confocal stack. (**D**) Two MN axons (arrows) leave the VproN and innervate the sternal anterior rotator muscle (orange). (**E**) The sternal anterior rotator muscle fibers originate from the sternal cuticle and insert on the medial edge of the rim of the coxal. Muscle contraction would likely tend to rotate the coxa about its long axis, and to adduct the coxa medially. (**F**) MN 7 exits the prothoracic accessory nerve (ProAN) (**G**) A single axon leaves the ProAN. (**H**) The same axon innervates the sternal adductor muscle fibers in the XNH volume. (**I**) Contraction of the sternal adductor muscle causes adduction (medial movement) of the coxa. The origins of the muscle fibers are not visible in the XNH volume. The muscle fibers insert on the posterior edge of the rim of the coxa cuticle.

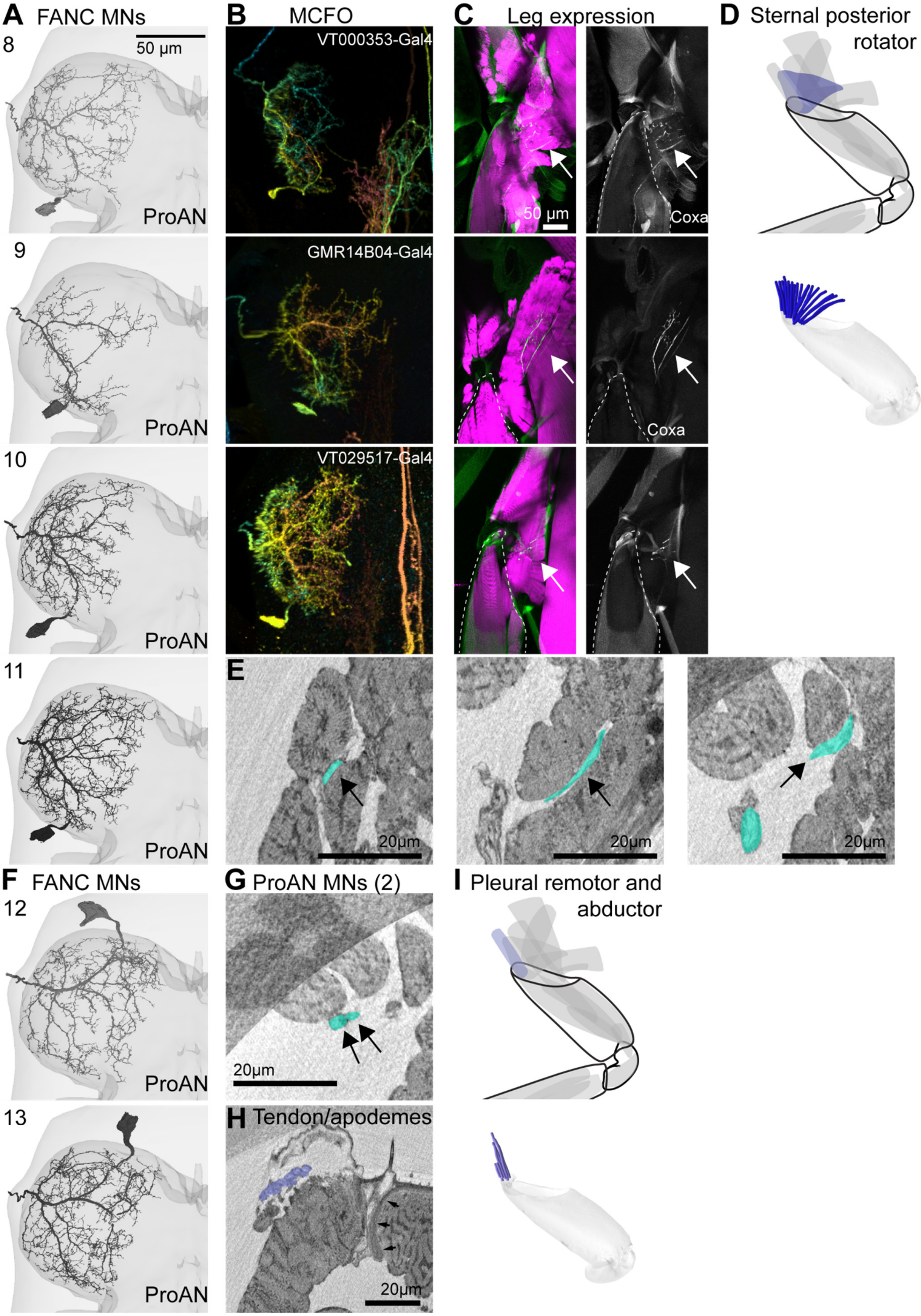

A FANC MNs
B MCFO
C Leg expression
D Sternal posterior rotator
8
50 µm
VT000353-Gal4
50 µm
Coxa
ProAN
9
GMR14B04-Gal4
Coxa
ProAN
10
VT029517-Gal4
ProAN
11
E
20µm
20µm
20µm
ProAN
F FANC MNs
G ProAN MNs (2)
I Pleural remotor and abductor
12
20µm
ProAN
13
H Tendon/apodemes
20µm
ProAN

Figure A4. MNs 8-11 innervate the sternal posterior rotator muscle. MNs 12-13 innervate the pleural remotor and abductor muscle. (A) MNs 8-11 have cell bodies on the posterior cortex of the left T1 neuropil and axon that exit through the ProAN. (B) MCFO clones (VT000353-Gal4, GMR14B04-Gal4, VT029517-Gal4) (C) Leg expression in Gal4 driver lines. Left, GFP with phalloidin counterstain (magenta). Right, GFP channel in grayscale. Dashed line indicates the coxal cuticle. Arrows indicate MN axons in the sternal posterior rotator. (D) Schematic and annotation of muscle fibers in XNH volume. The origins of the muscle fibers are unclear. The muscle fibers insert onto the posterior edge of the proximal rim of the coxa. Muscle contraction would tend to pull the coxal posteriorly (remotor), or rotate the coxal about its axis. (E) Example images of 3 of four MN axons (cyan) leaving the ProAN to innervate the sternal posterior rotator muscle fibers (XNH). (F) MNs 12-13 have axons that exit through the ProAN. (G) Two MN axons (cyan) pass through the muscle fibers of the sternal posterior rotator muscle (D) before finding and running along the muscle fibers of the pleural remotor and abductor muscle. (H) The origins of the muscle fibers are unclear in the XNH volume, but the insertions can be distinguished from the sternal posterior rotator fibers because they connect via apodemes (or tendons, blue), rather than directly onto cuticle as the sternal posterior rotator fibers do (arrowheads). (I) Annotated muscle fibers in the XNH volume. Muscle contraction would tend to pull the coxa laterally (abduction).

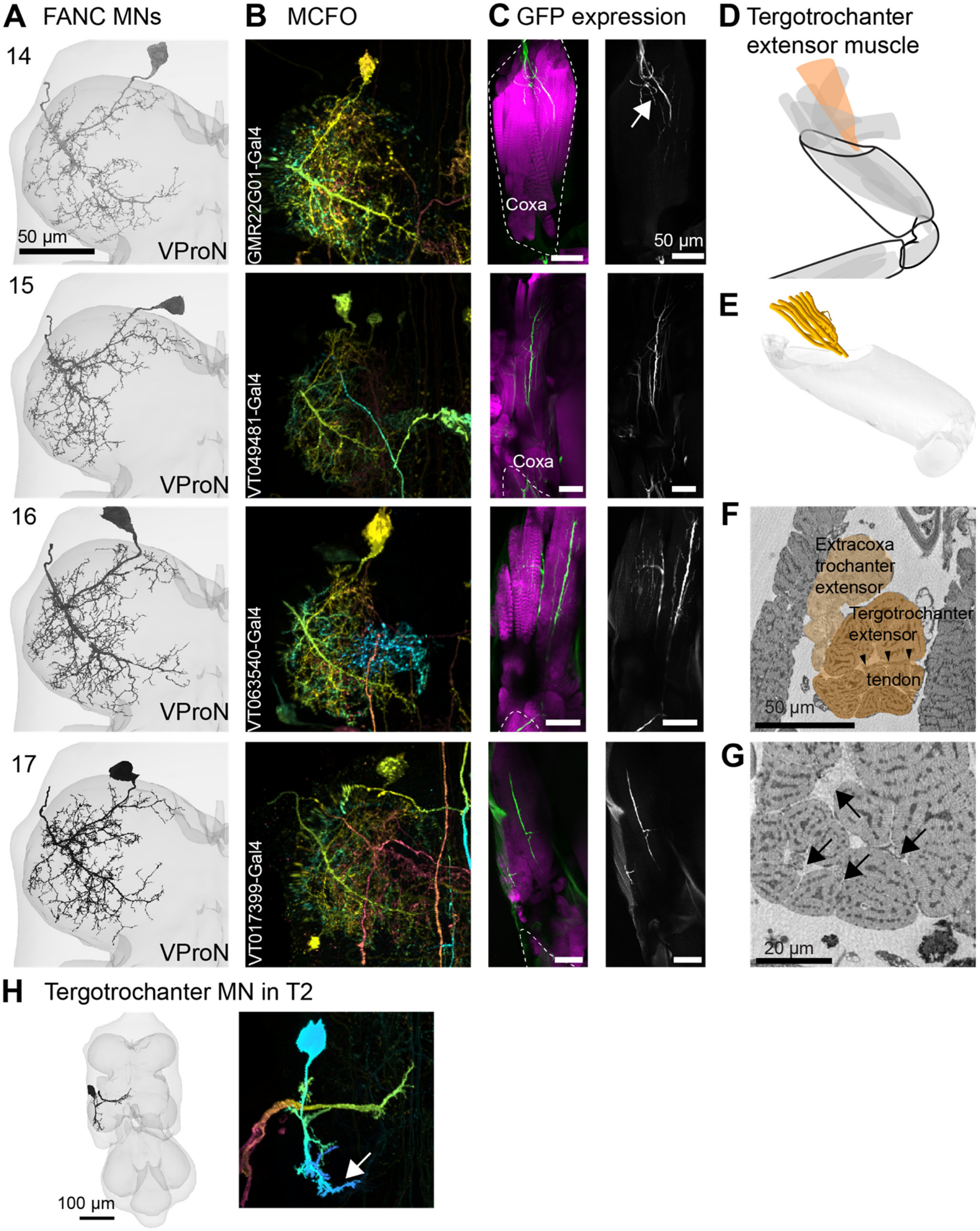

A FANC MNs
B MCFO
C GFP expression
D Tergotrochanter extensor muscle
14
50 µm
VProN
GMR22G01-Gal4
Coxa
50 µm
E
15
VProN
VT049481-Gal4
Coxa
16
VProN
VT063540-Gal4
F
Extracoxa trochanter extensor
Tergotrochanter extensor
tendon
50 µm
17
VProN
VT017399-Gal4
G
20 µm
H Tergotrochanter MN in T2
100 µm

**Figure A5. MNs 14-17 innervate the tergotrochanter muscle.** (**A**) MNs 14-17 have axons that exit through the VProN. The anterior exit point, relative to the leg nerve, together with the medial branch gives the neurons a distinctive rotated L-shape. (**B**) MCFO clones with this distinctive rotated 'L' shape (GMR22G01-Gal4, VT049481-Gal4, VT063540-Gal4, VT017399-Gal4). (**C**) GFP expression in the periphery. By luck, in the GMR22G01-Gal4 sample, the tergotrochanter muscle fibers rolled into the coxa. The other samples show axon labeling in the thorax, in muscle fibers that originate at the tergum. The coxa is outlined with a dashed line. (**D**) Schematic of the tergotrochanter muscle. (**E**) Annotated muscle fibers in the XNH volume. Muscle contraction pulls on the trochanter extensor tendon to extend the trochanter and femur. (**F**) The tergotrochanter muscle fibers (dark orange) insert on the trochanter extensor tendon (arrowheads). Another muscle, the sternotrochanter extensor (light orange), is composed of fibers that originate from the sternal cuticle and also insert onto the same tendon. (**G**) The longitudinal axons in C can be seen in cross-section within the tergotrochanter muscle. (**H**) A final piece of evidence supporting the claim that MNs 14-17 innervate the prothoracic tergotrochanter muscle is that they share a similar morphology with the well-known jump MN in T2. A large branch elaborates in the posterior-lateral portion of the neuromere (arrow). Like the MNs 14-17 that exit the VProN, the T2 TT MN exits a different nerve from other leg MNs.

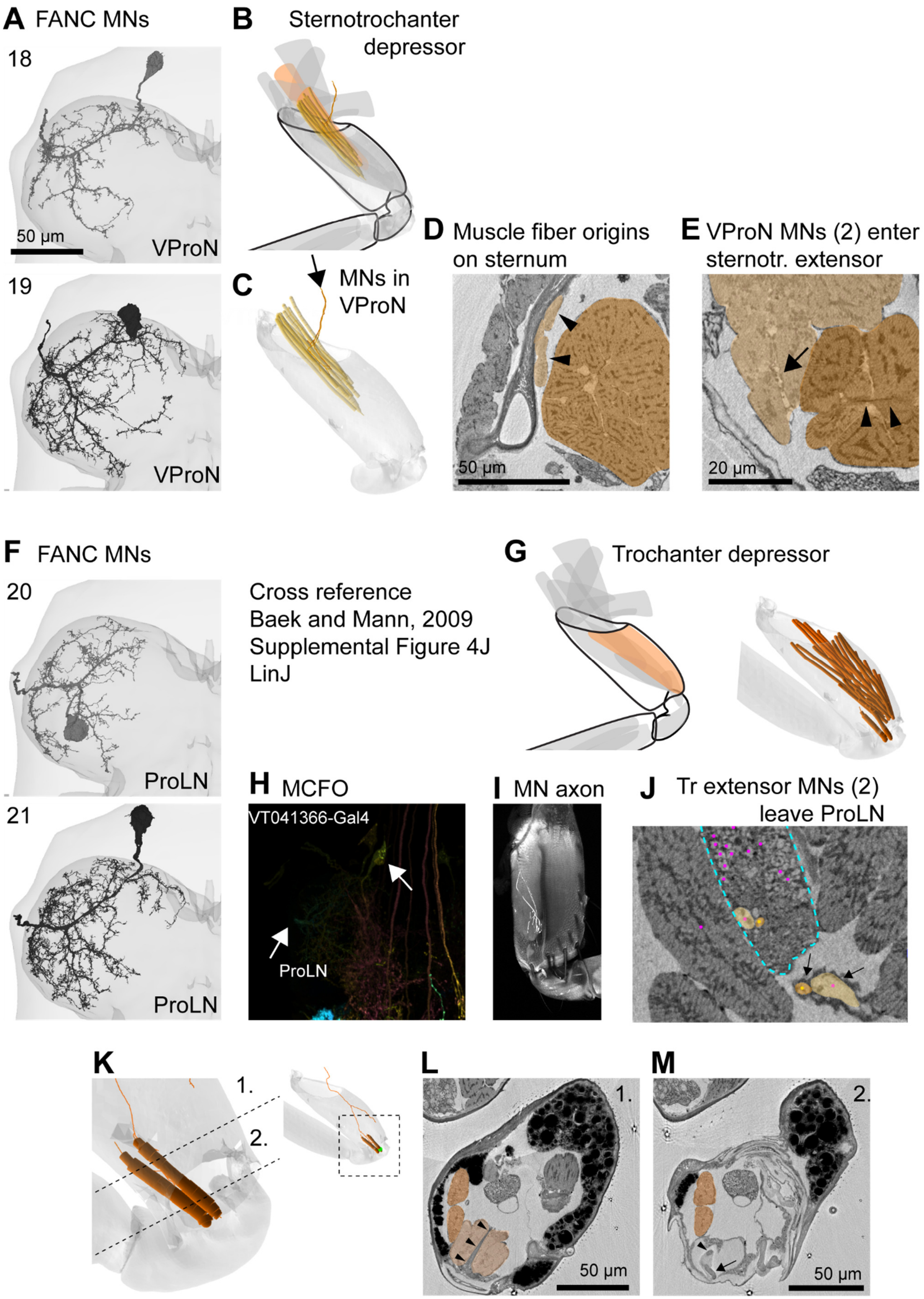
A FANC MNs
18
50 μm
VProN
19
VProN
B Sternotrochanter depressor
C MNs in VProN
D Muscle fiber origins on sternum
50 μm
E VProN MNs (2) enter sternotr. extensor
20 μm
F FANC MNs
20
ProLN
21
ProLN
Cross reference
Baek and Mann, 2009
Supplemental Figure 4J
LinJ
G Trochanter depressor
H MCFO
VT041366-Gal4
ProLN
I MN axon
J Tr extensor MNs (2) leave ProLN
K
1.
2.
L
1.
50 μm
M
2.
50 μm

**Figure A6. MNs 18-19 innervate the sternotrochanter extensor muscle. MNs 20-21 innervate the trochanter extensor muscle in the coxa.** (A) The axons of MNs 18-19 exit the VProN. (B) Schematic of the sternotrochanter extensor muscle. Orange indicates muscles that extend the coxa-trochanter joint, as during reaching. We did not find any Gal4 lines that label either MN. (C) The annotated muscle fibers of the sternotrochanter extensor muscle. The fibers insert on the trochanter extensor tendon, the same tendon in E and Figure A6F. The insertion of these fibers is not shown. (D) The muscle fibers (light orange) originate on the sternal cuticle of the thorax (arrowheads). The neighboring tergotrochanter extensor (dark orange) in cross-section looks like capocollo. (E) Two MN axons leave the VProN to innervate (arrows) the extracoxal trochanter extensor fibers (light orange). The trochanter extensor tendon can be seen (arrowheads) within the tergotrochanter fibers (dark orange). (F) MNs 20-21 exit the prothoracic leg nerve (ProLN) and innervate the trochanter extensor muscle in the coxa. MN 20 has a soma on the dorsal cortex of the neuromere. We did not find any MCFO clones for this neuron, but in the work of Baek and Mann, a MARCM clone, arising from lineage LinJ, with a dorsal soma was found to innervate the trochanter depressor (Baek and Mann, 2009). (G) Schematic of the trochanter extensor muscle. Right, annotated muscle fibers in XNH volume. The fibers originate from the interior surface of the coxa and insert on the trochanter extensor tendon. (E, arrowheads) (H) A dim MCFO clone (VT041366-Gal4). (I) GFP expression in the coxa. (J) Two axons (arrows) leave the ProLN (dashed cyan line) and eventually innervate the trochanter extensor muscle fibers. (K) Two annotated muscle fibers that are innervated by one of the two trochanter extensor MNs (inset). These two fibers do not insert onto the extensor tendon like the rest of the muscle fibers. Instead, they insert directly onto the surface of the trochanter. (L) Plane 1. Indicated in K. The two accessory muscle fibers (dark orange) bypass the trochanter extensor tendon (arrowheads). (M) Plane 2. Indicated in K. The trochanter extensor tendon (arrowhead) inserts onto a small sclerite on the trochanter (arrow). The two accessory fibers continue to their more distal insertion site. More details on the action and movement of the coxa-trochanter joint are shown below (Figure A7).

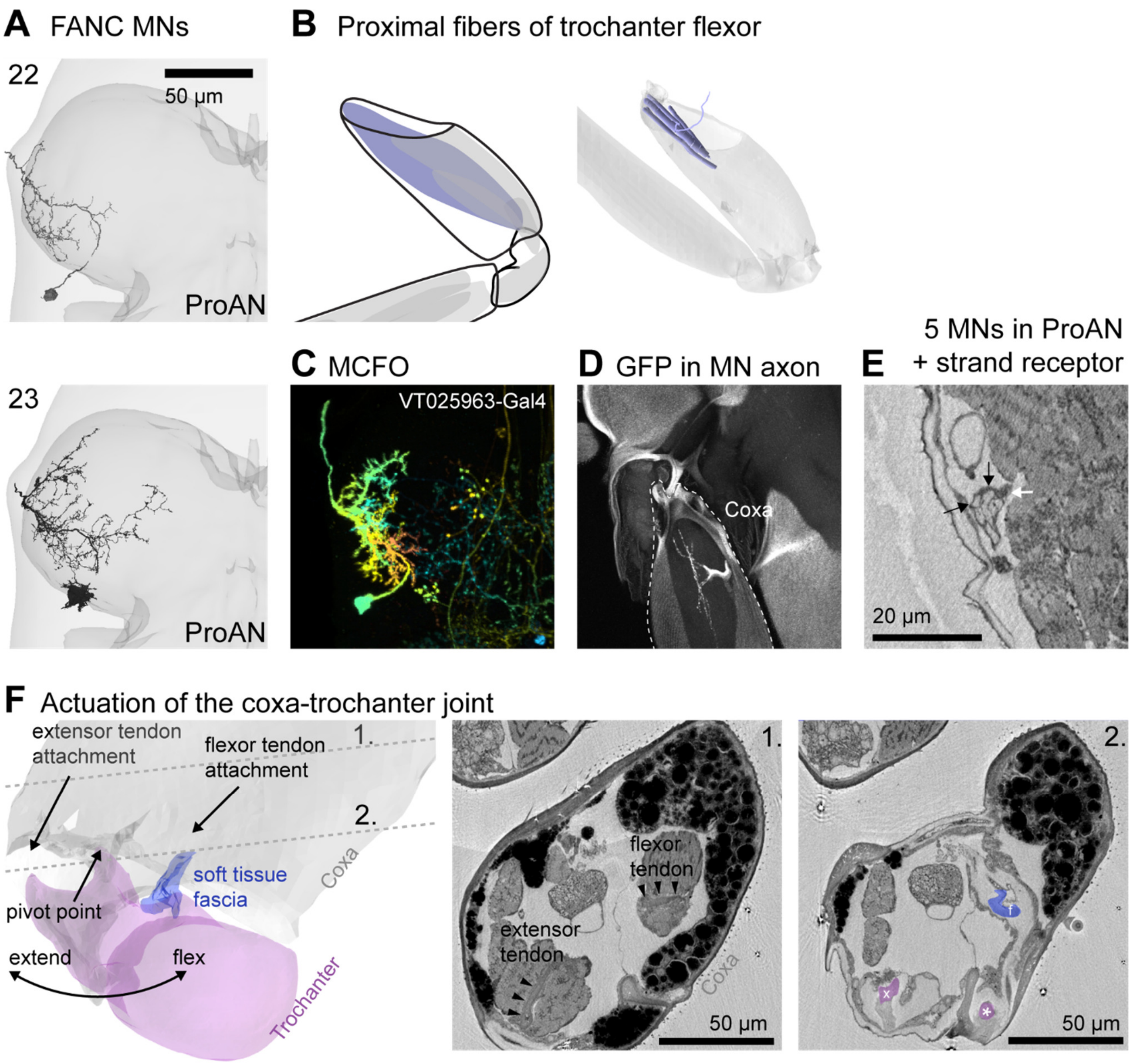

**Figure A7. MNs 22-23 have somas on the posterior cortex of the neuromere and innervate the proximal fibers of the trochanter flexor muscle.** (A) Two MNs in FANC have somas on the posterior cortex of the neuropil that are smaller than the MNs that innervate the sternal posterior rotator muscle (Figure A4A). (B) Left, schematic of the trochanter flexor muscle in the coxa. Right, proximal muscle fibers of the trochanter flexor originate on the interior surface of the coxa and insert on the trochanter flexor tendon (not shown). Two axons from the ProAN innervate the proximal fibers. (C) MCFO clone (VT025963-Gal4). (D) GFP expression (VT025963-Gal4) in the proximal coxa (dashed line). (E) Five MN axons innervate the trochanter flexor muscle (XNH volume), with two innervating the proximal fibers (black arrows). Thus far we have accounted for seven of the 12 neurons that exit the ProAN in FANC. The five neurons in this nerve account for the remainder, and all five innervate the trochanter flexor muscle. The sixth object in this nerve is the strand receptor. Together, these data support the claim that two neurons with posterior somas innervate the trochanter flexor muscle in the coxa. (F) Actuation of the coxa-trochanter joint. Left: segmented volumes of coxa (gray), trochanter (purple), and connective tissue connecting the trochanter flexor tendon to a protuberance of the trochanter cuticle (blue). The image planes 1. and 2. indicate the images to the right. 1.) The antagonist extensor and flexor muscles insert on the tendons (arrowheads). 2.) More distally, the extensor tendon (x) and flexor tendon (f) connect to the trochanter on opposite sides of a small protuberance of the trochanter that rests in a small cavity of the coxa. We propose that the extensor and flexor muscles move the trochanter about this pivot point.

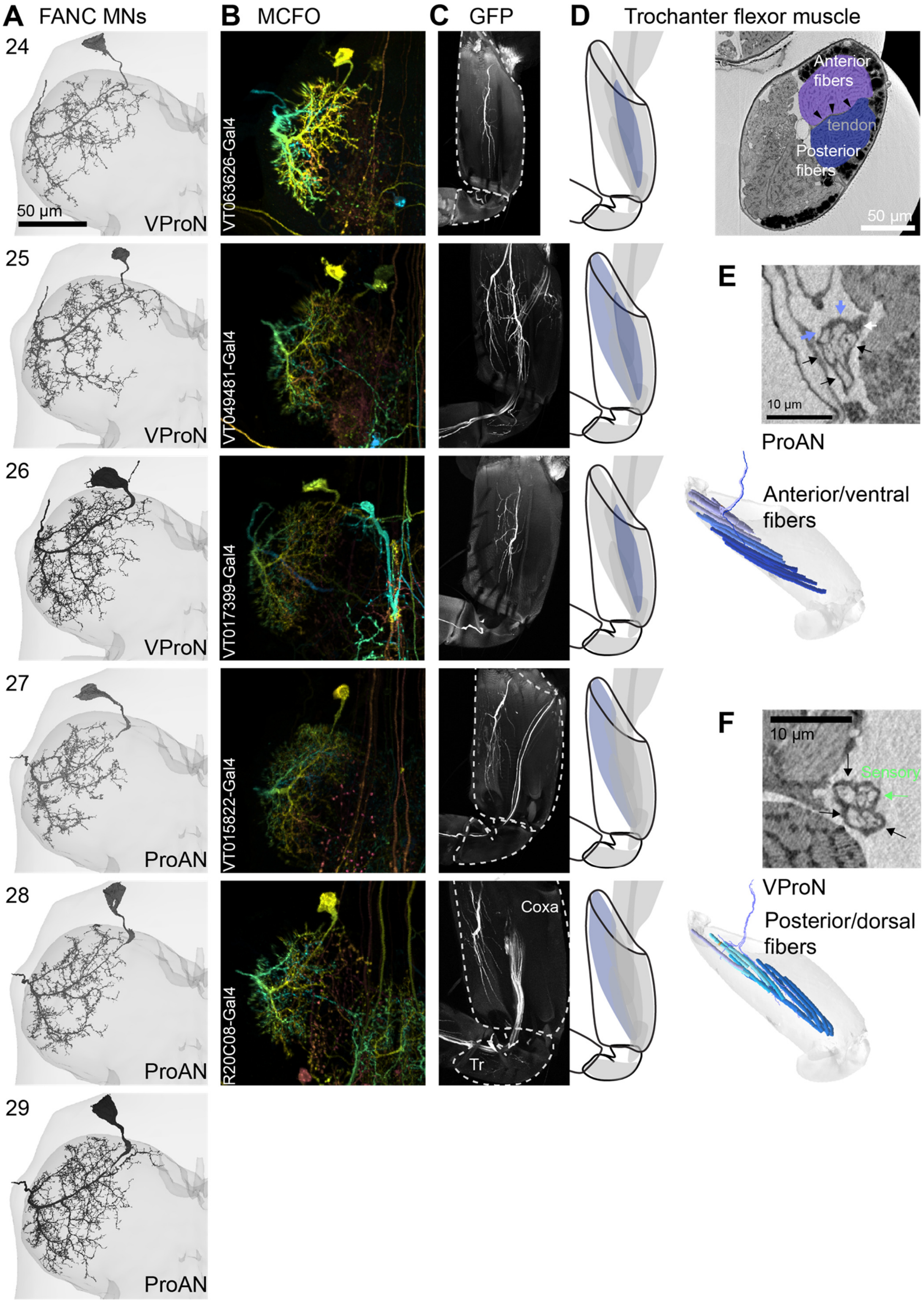

A FANC MNs
B MCFO
C GFP
D Trochanter flexor muscle
24
25
26
27
28
29
50 µm
VProN
ProAN
VT063626-Gal4
VT049481-Gal4
VT017399-Gal4
VT015822-Gal4
R20C08-Gal4
Coxa
Tr
Anterior fibers
tendon
Posterior fibers
E
10 µm
ProAN
Anterior/ventral fibers
F
Sensory
VProN
Posterior/dorsal fibers

**Figure A8. MNs 24-29 travel along the ProAN (N=3) or the VProN (N=3) to innervate the anterior or posterior fibers of the trochanter flexor muscle, respectively.** (A) Trochanter flexor MNs have characteristic morphology, with more lateralized branching in the neuropil and swooping primary neurites. The genetic basis of this morphology was investigated by Enriquez et al. (2015). Three of the neurons exit via the ProAN, three exit the VProN. (B) MCFO clones with characteristic morphology (VT063626-Gal4, VT049481-Gal4, VT017399-Gal4, VT015822-GAL4, GMR20C08-GAL4). It can be unclear via which nerve the axons travel. (C) Leg images of GFP expression in the coxa (dashed outlines). (D) Schematics of the portion of the muscle innervated in each line. The XNH volume reveals the trochanter flexor tendon (arrowheads) starts as a wide band that bisects the trochanter flexor muscle, with anterior muscle fibers (light blue) inserting on one side and posterior fibers (dark blue) inserting on the other side. (E) A branch of the ProAN innervates the anterior fibers, carrying five MNs and the strand receptor (white arrow). Two MNs that innervate proximal fibers are indicated with pale blue arrows, possibly the neurons with small posterior somas (Figure A7). A third (of the five) innervates the same proximal fibers. If the two neurons with dorsal somas are functional, this would be an example of polyneural innervation. (F) A branch of the VProN carries three MNs that innervate the posterior fibers, along with sensory axons.

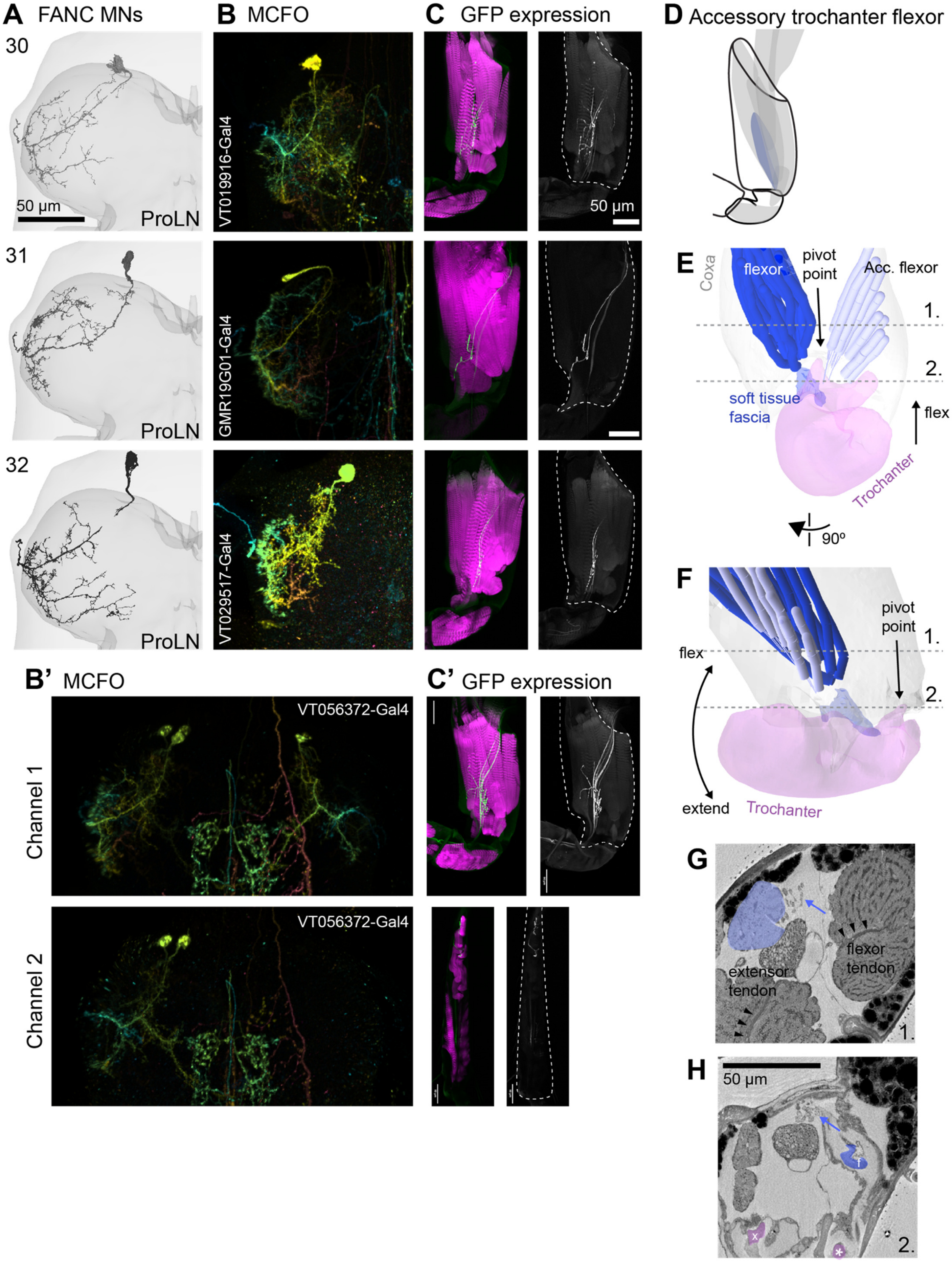

A FANC MNs
30
31
32
50 μm
ProLN
B MCFO
VT019916-Gal4
GMR19G01-Gal4
VT029517-Gal4
C GFP expression
50 μm
D Accessory trochanter flexor
E
Coxa
flexor
pivot point
Acc. flexor
1.
2.
soft tissue fascia
flex
Trochanter
90º
F
pivot point
flex
extend
Trochanter
B' MCFO
Channel 1
Channel 2
VT056372-Gal4
C' GFP expression
G
extensor tendon
flexor tendon
H
50 μm
f
x

**Figure A9. MNs 30-32 travel the ProLN to innervate the accessory trochanter (tr.) flexor muscles in the coxa.** (**A**) Accessory tr. neurons exhibit a similar morphology as tr. flexor neurons (Figure A8), with lateralized dendrites and swooping primary neurites, but with fewer branches and axons that travel the ProLN. (**B**) MCFO clones of accessory tr. flexor MNs. B') In one example line, two somas are visible in both channels, but the dendrites of the accessory tr. flexor neurons are visible in one channel whereas the dendrites of an ltm MN are visible in channel 2. (**C**) GAL4-driven GFP expression in the coxa (green) with phalloidin counterstain. Left: GFP expression. C') Coxa and tibia GFP expression showing innervation of both accessory trochanter flexor and ltm1 muscle in the tibia. (**D**) Schematic of accessory tr. flexor muscle. (**E**) View of the coxa-trochanter joint from an anterior point of view. The trochanter appears to pivot around a protuberance of the trochanter cuticle that fits into a cavity in the distal coxa. The accessory tr. flexor muscle fibers (light blue) insert onto a similar piece of trochanter cuticle as the tr. flexor tendon, but originate on the opposite, posterior-lateral surface of the coxa from the trochanter flexor fiber (dark blue). The image planes, 1. and 2., indicate the images in G and H. (**F**) A rotated view of the coxa-trochanter joint, from a lateral point of view, in the approximate plane of actuation. (**G**) XNH image at plane 1. The accessory tr. flexor fibers (light blue) connect to individual tendons or apodemes (blue arrow), rather than a single large tendon. The trochanter flexor and extensor muscle and tendons (arrowheads) are visible. (**H**) XNH image at more distal plane 2. The accessory tr. flexor tendons eventually insert at the same tr. location as the flexor tendon and its connective tissue.

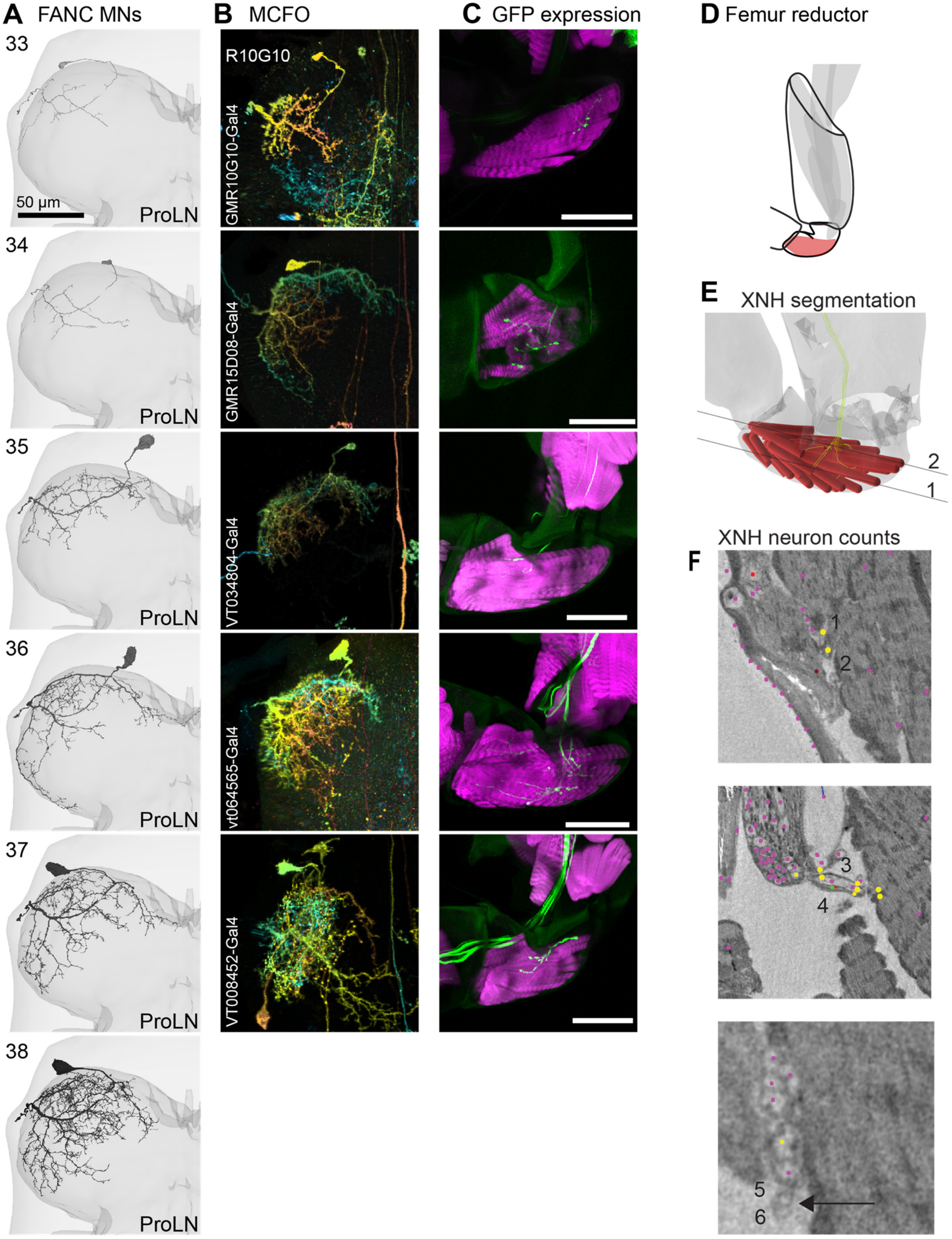

A FANC MNs
33
50 µm
ProLN
34
35
36
37
38
B MCFO
R10G10
GMR10G10-Gal4
GMR15D08-Gal4
VT034804-Gal4
vt064565-Gal4
VT008452-Gal4
C GFP expression
D Femur reductor
E XNH segmentation
2
1
XNH neuron counts
F
1
2
3
4
5
6

**Figure A10. MNs 33-38 innervate the femur reductor muscle in the trochanter.** (**A**) The genetic basis of femur reductor MN morphology was investigated by Enriquez et al. (2015). The MNs arborize in the anterior and medial portions of the neuropil, the medial branch being characteristic. (**B**) MCFO clones of femur reductor MNs. (**C**) GAL4-driven GFP expression in the trochanter (green) with phalloidin (magenta) counterstain. (**D**) Schematic of femur reductor muscle. (**E**) Muscle fiber and MN annotation in XNH data. (**F**) XNH images of MNs leaving the leg nerve, including 2 very small diameter axons, likely the top two MNs in (A).

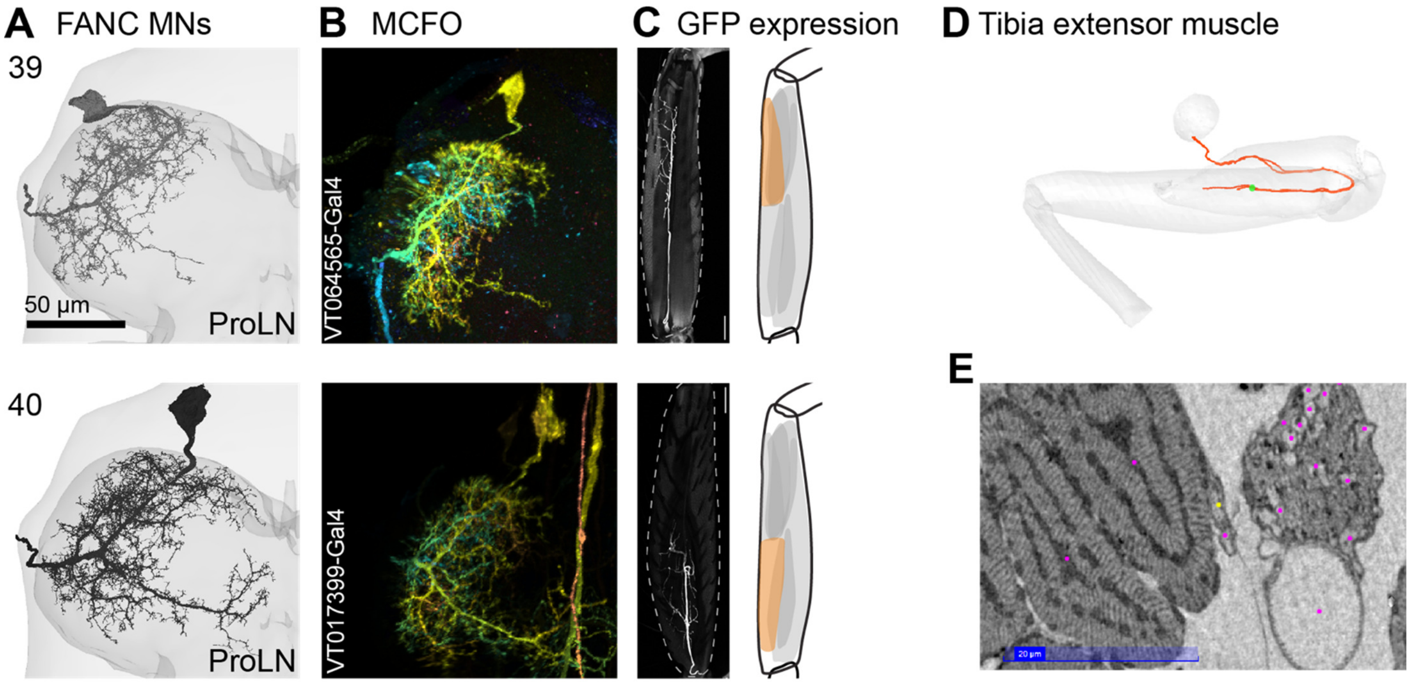

**Figure A11. MNs 39 & 40 are the slow extensor tibiae (SETi) and the fast extensor tibiae (FETi) MNs.** (**A**) The genetic basis of the SETi MN morphology was investigated by Enriquez et al. (2015). The MNs have a characteristic medial branch in the posterior portion of the T1 neuropil. (**B**) MCFO clones of tibia extensor MNs. (**C**) GAL4-driven GFP expression in the femur (outline). The SETi innervates the distal fibers of the muscle with are more pinnate. (**D**) XNH annotation of muscle fibers and MNs. The SETi targets the distal fibers of the muscle (C). The distal fibers are more pinnate, suggesting less mechanical advantage, perhaps a mechanism underlying the smaller forces produced by spikes in the SETi. (**E**) Cross-section of XNH volume through the femur. The tibia extensor MNs leave the ProLN and pass through a membrane that separates the extensor muscle from the flexor muscle, the ltm2 muscle, and the nerve, like fascia in vertebrate musculature.

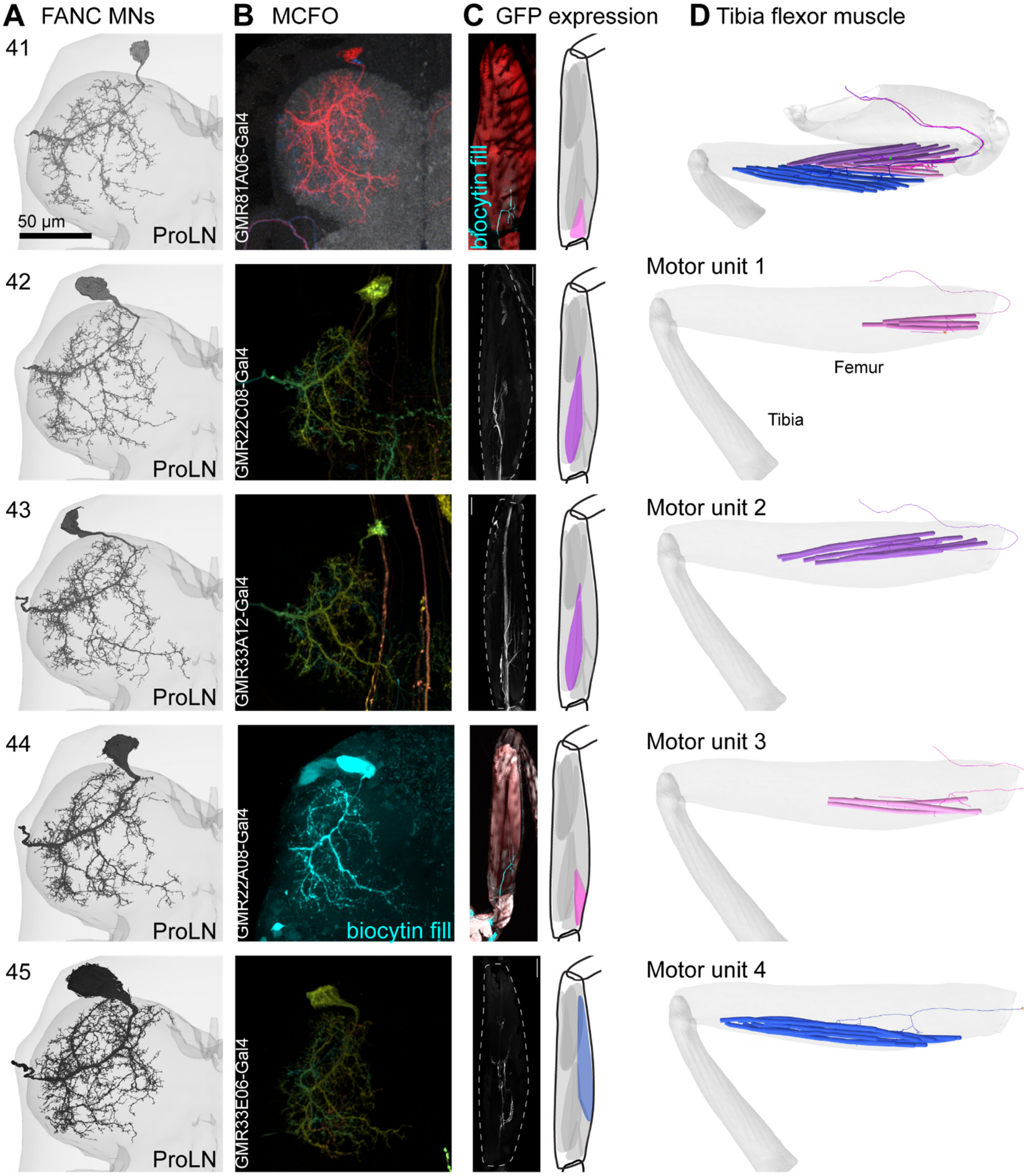
A FANC MNs
B MCFO
C GFP expression
D Tibia flexor muscle
41
42
43
44
45
50 µm
ProLN
GMR81A06-Gal4
GMR22C08-Gal4
GMR33A12-Gal4
GMR22A08-Gal4
GMR33E06-Gal4
biocytin fill
Motor unit 1
Motor unit 2
Motor unit 3
Motor unit 4
Femur
Tibia

**Figure A12. MNs 41-45 innervate the tibia flexor muscle.** (**A**) The tibia flexor MNs elaborate throughout the T1 neuromere. Salient morphological features include a dorsal-posterior branch, shared with accessory tibia flexor MNs (A13-A14), and prominent medial branches. The largest MN by volume in left T1 is the Fast tibia flexor (MN #45, Azevedo et al. 2020). The Fast tibia flexor MN in left T1 lacks the medial branch, though the right T1 pair does have a medial branch (not shown). (**B**) We have recorded from tibia flexor MNs in our past work, so some confirmation of the tibia flexor morphology comes from stains of biocytin fills during recordings. In those cases–#41, #44, and #45 (not shown)–we have also measured tibia force production from eliciting spikes in the MN. (**C**) Leg expression of GFP or biocytin fills. (**D**) Motor units in the tibia flexor muscle. The one motor unit we are unclear about is the motor unit innervated by the most distal axon. Baek and Mann (2009) reported a similar axon in the tibia flexor muscle, Fe X. In our EMG recordings in the tibia flexor muscle, we did not report a second identifiable unit in the distal tibia, but we did observe a third cluster in our GCaMP imaging of the tibia flexor muscle that we did not extensively analyze, which could be due to the action of this neuron.

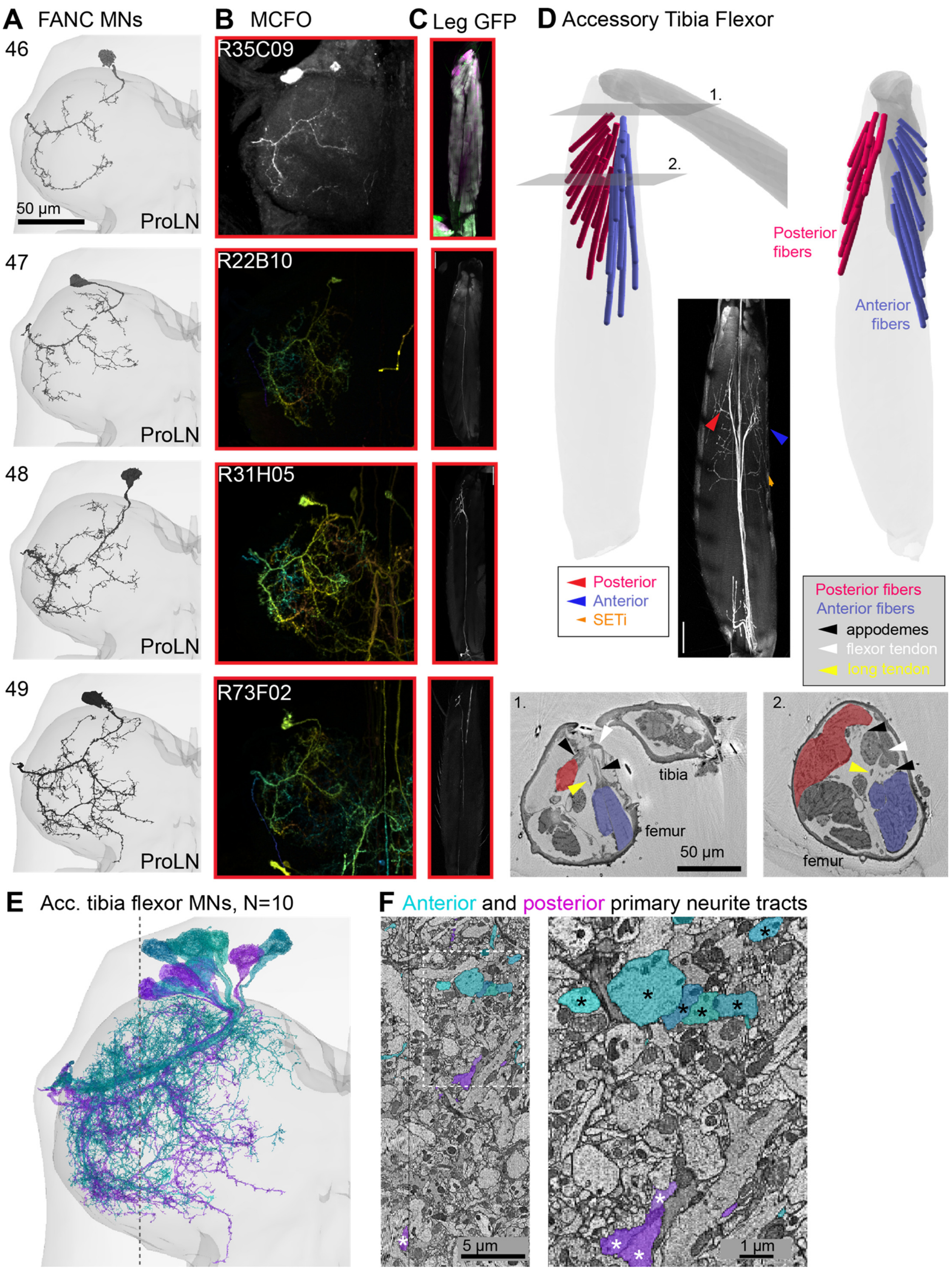

A FANC MNs
46
47
48
49
50 µm
ProLN
B MCFO
R35C09
R22B10
R31H05
R73F02
C Leg GFP
D Accessory Tibia Flexor
1.
2.
Posterior fibers
Anterior fibers
Posterior
Anterior
SETi
Posterior fibers
Anterior fibers
appodemes
flexor tendon
long tendon
tibia
femur
50 µm
E Acc. tibia flexor MNs, N=10
F Anterior and posterior primary neurite tracts
5 µm
1 µm

Figure A13. MNs 46-49 innervate the accessory tibia flexor muscle. We hypothesize that MNs 46-49 innervate the muscle fibers that originate from the posterior surface of the femur, though we are not certain. (A) Four of the ten tibia accessory MNs in FANC. A characteristic feature is the thin posterior process that leaves the primary neurite near the exit point. This process also projects dorsally. (B) MCFO clones and a biocytin fill from Azevedo et al. 2020 (GMR35C09-GAL4). (C) Leg expression of GFP or biocytin fills. The axons of these accessory tibia flexor MNs innervate the posterior fibers. (D) Anatomy of the accessory tibia flexor muscle, with posterior fibers in red and anterior fibers in blue. Inset shows a confocal image of two branches of accessory tibia flexor MNs projecting either anteriorly or posteriorly in the same image. The SETi is also labeled and identified (orange). (E) All ten accessory tibia flexor MNs. In a separate observation in FANC, these four MNs 46-49 (magenta) travel in a separate tract from the other six MNs, MNs 50-55 (cyan). (F) YZ-plane through the FANC EM volume at the gray dashed line in the image at left, showing that six cyan MNs run closely together (black asterisks), whereas the magenta neurons run in a more posterior tract (white asterisks). One possible explanation is that the magenta neurons innervate the posterior fibers and the cyan neurons tend to innervate the anterior fibers. This is certainly true of the single MN labeled by 35C09-GAL4, which we studied in Azevedo et al. (2020). The neuron has a characteristic medial projection, like several magenta neurons, which is not a prominent feature of MNs 50-55 (cyan).

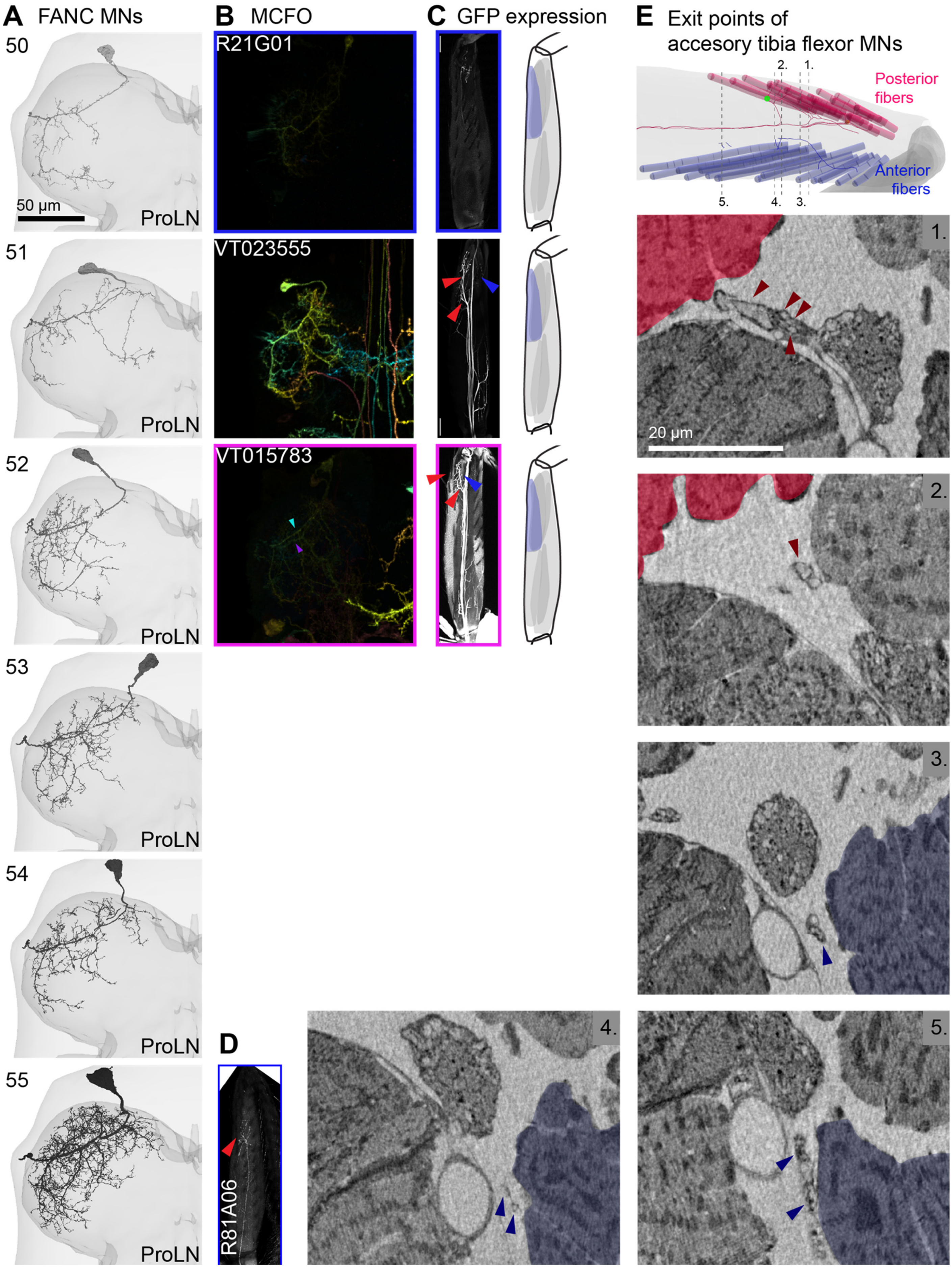
A FANC MNs
B MCFO
C GFP expression
E Exit points of accesory tibia flexor MNs
50
51
52
53
54
55
50 µm
ProLN
R21G01
VT023555
VT015783
D
R81A06
Posterior fibers
Anterior fibers
1.
2.
3.
4.
5.
20 µm

**Figure A14. MNs 50-55 innervate the accessory tibia flexor muscle.** (**A**) The remaining 6 accessory tibia flexor MNs. (**B**) MCFO clones from GAL4 lines that label axons innervating the anterior fibers of the accessory tibia flexors (GMR21G01-GAL4, VT023555-GAL4, VT015783-GAL4). We found more GAL4 lines that label neurons innervating the posterior fibers than anterior fibers. We found more GAL4 lines that label neurons innervating the posterior fibers than anterior fibers. (**C**) GFP expression in the leg showing axons targeting the anterior fibers. In one instance (VT015783-GAL4) the MCFO images showed two different MNs, one with an anterior primary neurite (cyan arrowhead), the other with a posterior neurite (magenta arrowhead), and the GAL4 line labeled axons targeting both the anterior and the posterior muscle fibers. (**D**) We recorded from an accessory tibia flexor MN with a large axon and filled it with neurobiotin. The axon targets anterior fibers (GMR81A06-GAL4). Spikes in this neuron produced large forces on the tibia (not shown), which leads us to hypothesize that we recorded from the largest accessory tibia flexor MN. Further experiments are necessary to confirm this hypothesis. (**E**) Axons leaving the ProLN and entering the accessory tibia flexor muscle. Top: schematic showing the location of the image planes below. The axons are indicated with arrowheads. The anterior (red) or posterior (blue) muscle fibers are shaded. We count 5 axons that innervate the anterior fibers and 5 that innervate the posterior fibers, which is inconsistent with the hypothesis that neurons 50-55 all innervate the anterior fibers (Figure A13). Importantly, the high number of accessory tibia flexor MNs is consistent with past work (Baek and Mann, 2009, Brierley et al., 2012) and supports our claim that MNs 46-55 innervate this muscle. (**F**) More work is required to precisely map the MNs in FANC to these axon exit points in XNH and to muscle fiber innervation.

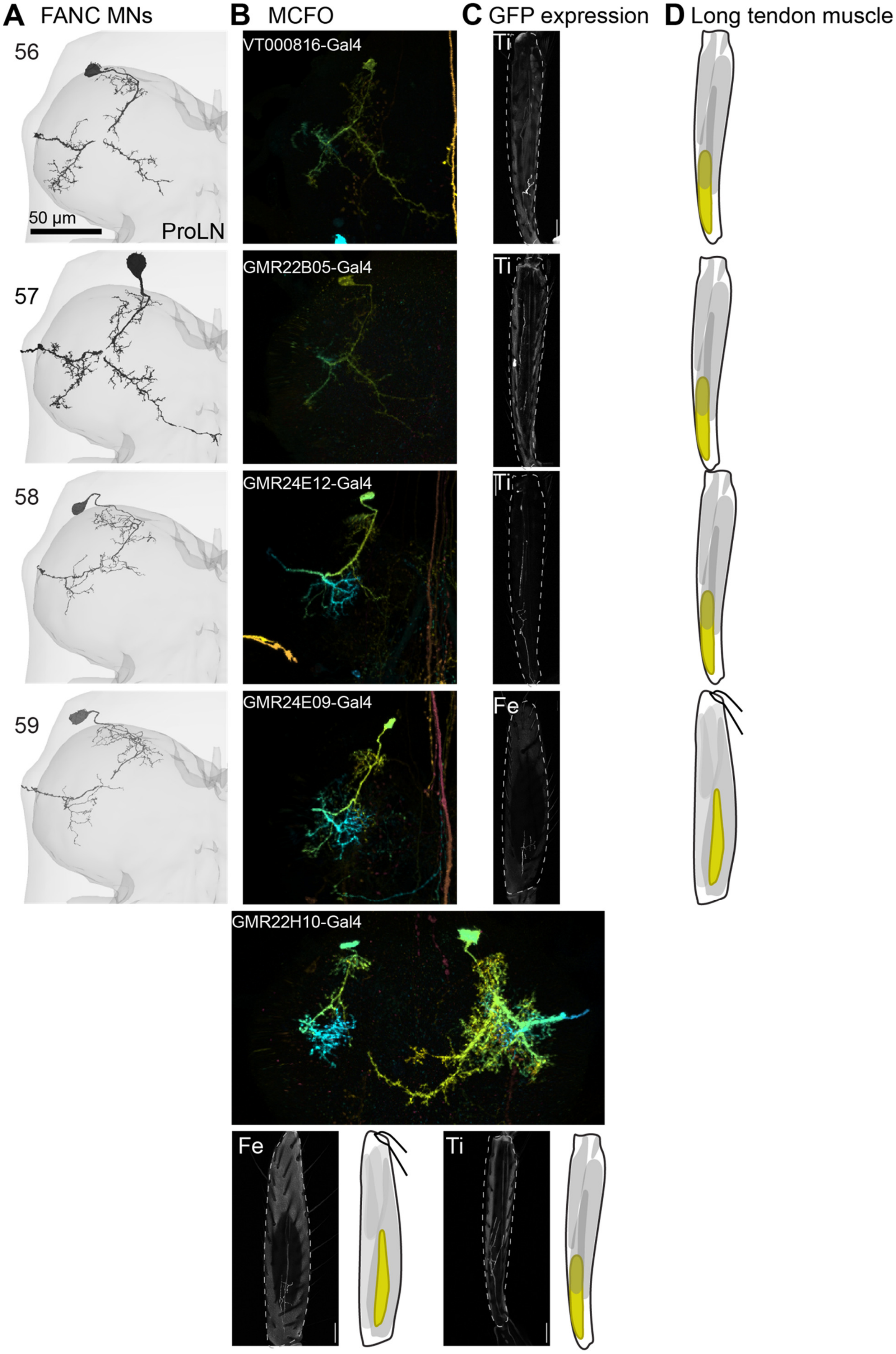
A FANC MNs
B MCFO
C GFP expression
D Long tendon muscle
56
57
58
59
50 µm
ProLN
VT000816-Gal4
GMR22B05-Gal4
GMR24E12-Gal4
GMR24E09-Gal4
GMR22H10-Gal4
Ti
Fe

**Figure A15. MNs 56-59 are small MNs that innervate the long tendon muscles in the femur and tibia.** (**A**) Reconstructed MNs in FANC. Two have medial branches, while two lack the medial branches. (**B**) MCFO clones (VT000816-GAL4, GMR22B05-GAL4, GMR24E12-GAL4, GMR24E09-GAL4). (**C**) GFP expression in the leg showing axons targeting either the ltm1 muscle in the tibia (Ti) or the ltm2 muscle in the femur (Fe). In one instance (GMR22H10-GAL4) the MCFO images showed two different MNs, one with the small morphology shown here and the other with the large, medially projecting morphology shown in Figure A16. We are certain that both ltm1 and ltm2 are innervated by an MN that lacks the medial branch, which both appear to express DIP-alpha (Venkatasubramanian et al., 2019). (**E**) Our data indicate that one of the MNs with a medial branch targets ltm1. We also see four axons innervating ltm2 in the femur in the XNH data (not shown). Together with the data in Figure A16, we conclude that both ltm muscles are innervated by one MN of each morphology.

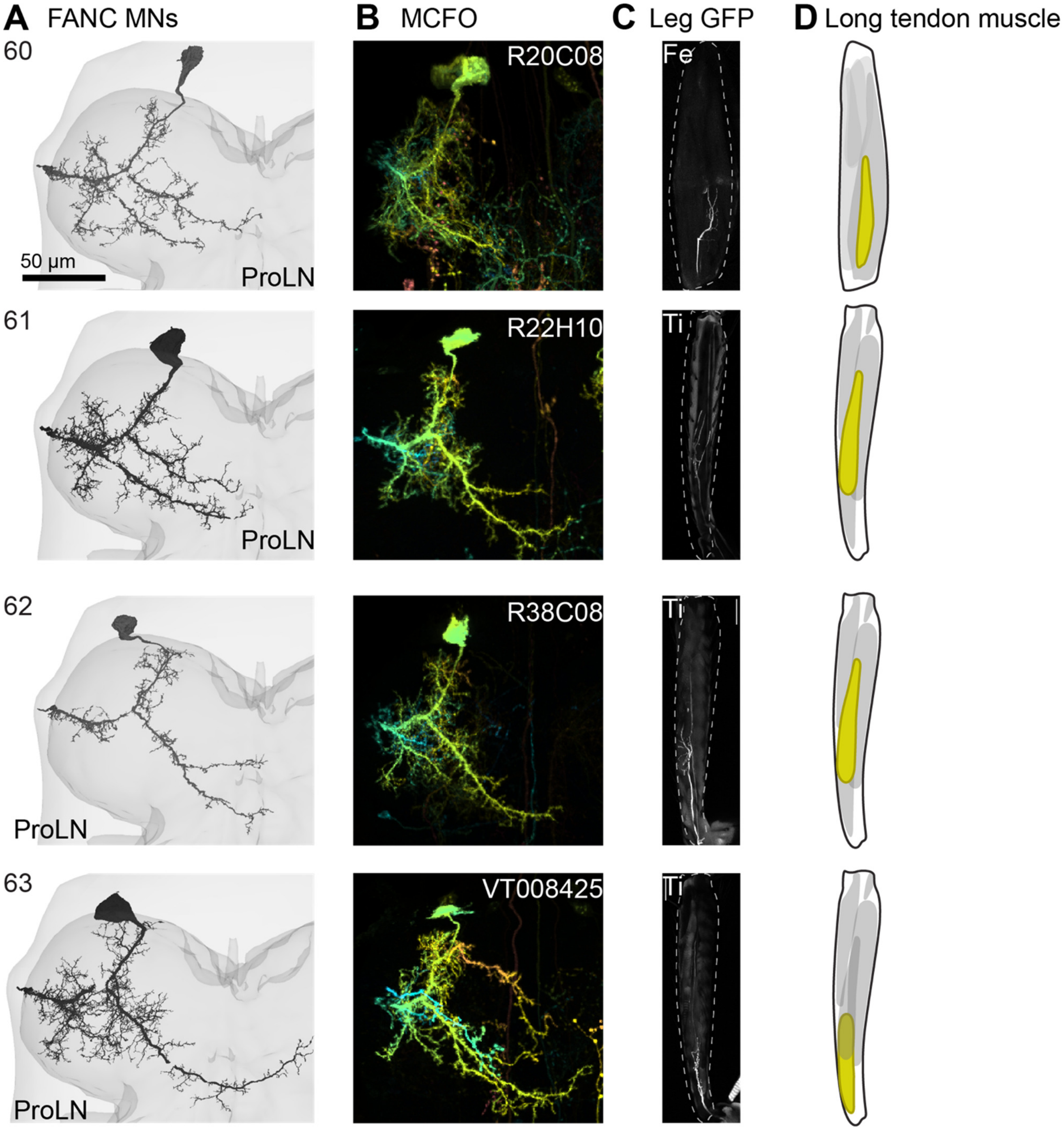

**Figure A16. MNs 60-63 are large MNs with medial branches that innervate the long tendon muscle.** (**A**) Reconstructed MNs in FANC. All four have branches that project medially, but two have medial dendrites that branch off at a more anterior location along the primary neurite, and the other two have additional medial branches that branch more posteriorly. (**B**) MCFO clones from GAL4 lines that label axons innervating ltm muscles in the femur (Fe) and tibia (Ti) (GMR20C08-GAL4, GMR22H10-GAL4, GMR38C08-GAL4, VT008425-GAL4). (**C**) GFP expression in the leg showing axons targeting the ltm muscles. Our data suggest that the MNs with posterior medial branches innervate the ltm2 in the femur. Consistent with this, the posterior medial branches arborize with a similar dendrite on tibia flexor MNs, which also innervate the femur.

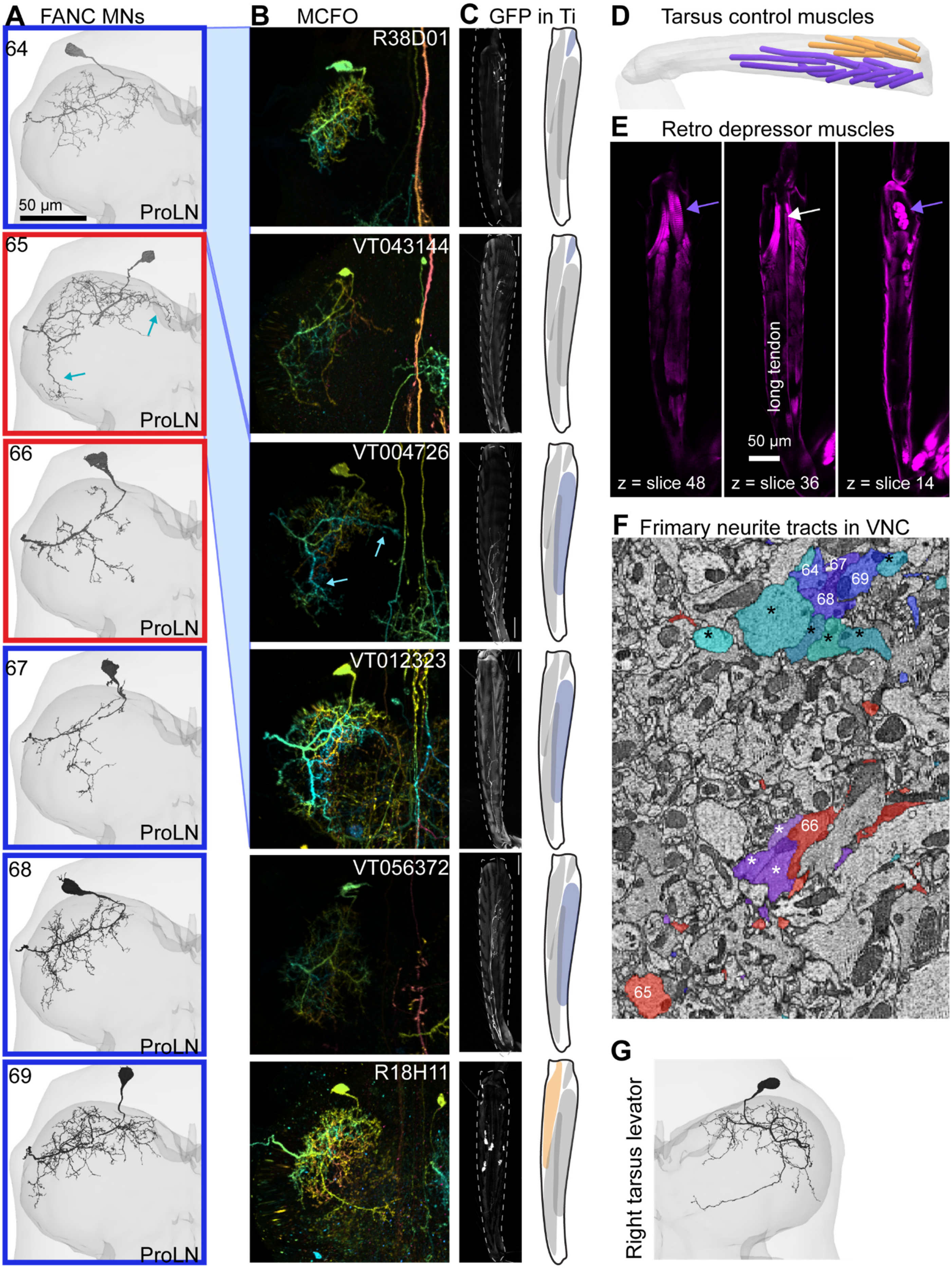
A FANC MNs
B MCFO
C GFP in Ti
D Tarsus control muscles
E Retro depressor muscles
F Frimary neurite tracts in VNC
G
64
65
66
67
68
69
50 µm
ProLN
R38D01
VT043144
VT004726
VT012323
VT056372
R18H11
long tendon
50 µm
z = slice 48
z = slice 36
z = slice 14
Right tarsus levator

**Figure A17. MNs 64-69 innervate the tarsus levator and depressor muscles.** (**A**) Tarsus control MNs appear to be the least elaborate MNs, with dendrites restricted to the anterior, lateral portion of the neuropil. The exception is the tarsus depressor MN with a U-shaped process in the ventral portion of the neuropil (65, cyan arrows). This neuron is the first MN born in the LinA lineage, which produces the majority of the MNs that travel in the ProLN (Baek and Mann, 2009; Venkatasubramanian et al., 2019). The edge colors indicate neurons with anterior (blue) or posterior (red) primary neurites, shown in (F). (**B**) MCFO clones from GAL4 lines that label axons innervating the tarsus control muscles. Clones with similar morphology are linked to the FANC neurons in (A) (VT043144-GAL4, VT040577-GAL4, GMR38D01-GAL4, VT004726-GAL4, VT012323-GAL4, GMR18H11). The ventral U-shaped dendrites of the large tarsus depressor neuron appear cyan in the depth-colored MIPs, indicated with cyan arrows. (**C**) GFP expression in the tibia (Ti). (**D**) Annotated muscle fibers in the incomplete tibia in the XNH volume. (**E**) Confocal images of tibia muscles (phalloidin, magenta), at three depths. The retro tarsus depressor muscle fibers are absent from the XNH volume. One set originates from the anterior, distal tibia cuticle (z=slice 48, blue arrow), and stretches proximally to insert on the tarsus depressor tendon (z=slice 36, white arrow). The other set originates from the posterior, distal tibia cuticle (z=slice 14, blue arrow), and stretches laterally to insert on the depressor tendon (z=slice 36, white arrow). The long tendon is visible in the same slice as the depressor tendon. (**F**) The primary neurites of the tarsus depressor and levator MNs (numbered) follow similar, grouped tracts as the accessory tibia flexor MNs (asterisks). (**G**) The right T1 tarsus levator MN shows a characteristic posterior, medial branch, which its paired neuron in left left T1 does not. To summarize, we are confident in the identity and muscle target of the large tarsus depressor MN (65), the tarsus levator MN (69), and the retro depressor MN (64). We are not certain of the precise muscle fibers innervated by the other three tarsus control MNs, but confident that they target tarsus muscles in the tibia. Past work suggested that tarsus depressor MNs outnumber tarsus levator MNs, consistent with our proposed targets here (Brierley et al., 2012).

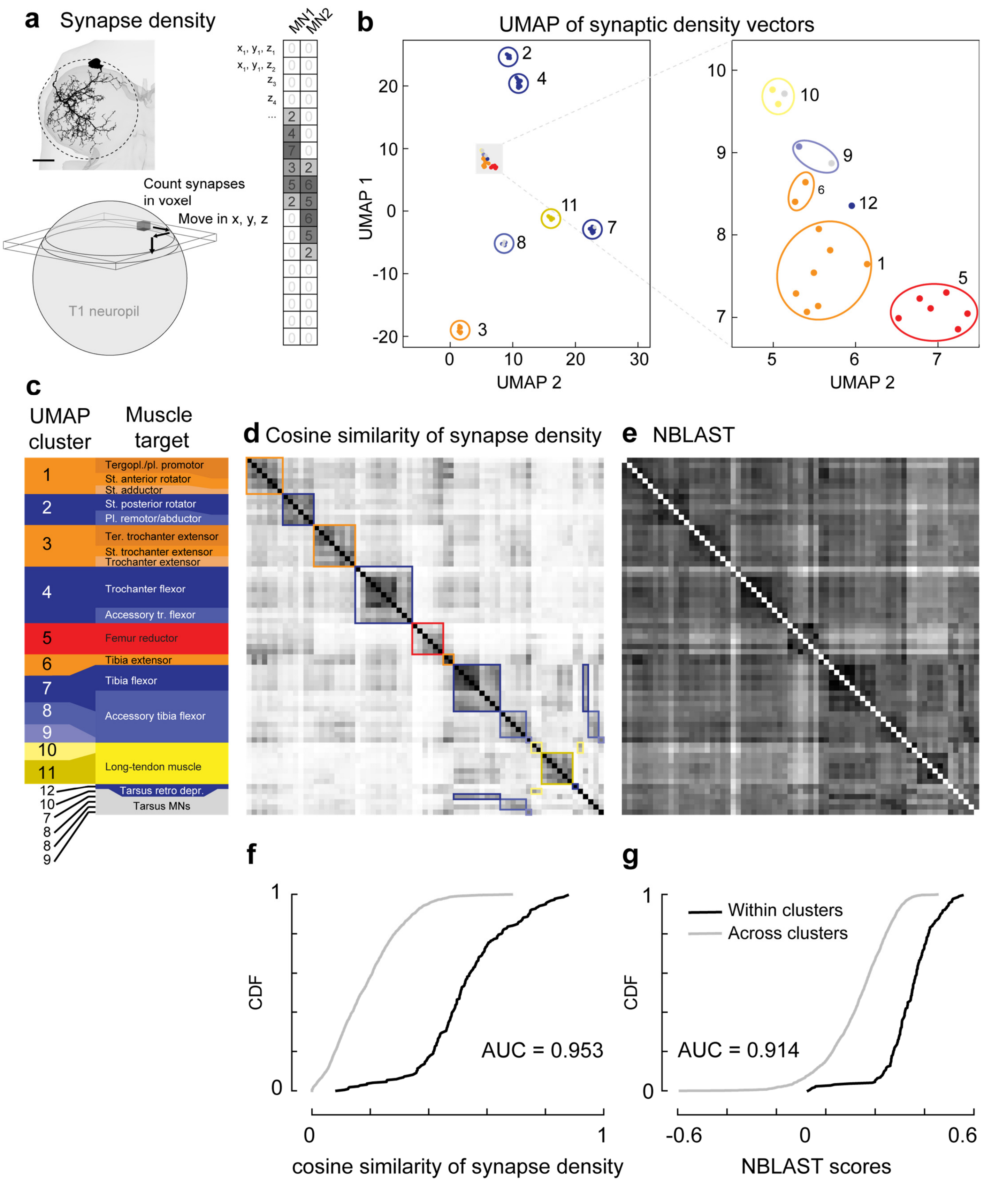
a Synapse density
MN1
MN2
Count synapses in voxel
Move in x, y, z
T1 neuropil
b UMAP of synaptic density vectors
UMAP 1
UMAP 2
c
UMAP cluster
Muscle target
Tergopl./pl. promotor
St. anterior rotator
St. adductor
St. posterior rotator
Pl. remotor/abductor
Ter. trochanter extensor
St. trochanter extensor
Trochanter extensor
Trochanter flexor
Accessory tr. flexor
Femur reductor
Tibia extensor
Tibia flexor
Accessory tibia flexor
Long-tendon muscle
Tarsus retro depr.
Tarsus MNs
d Cosine similarity of synapse density
e NBLAST
f
CDF
AUC = 0.953
cosine similarity of synapse density
g
Within clusters
Across clusters
AUC = 0.914
NBLAST scores

**Figure A18. UMAP algorithm identifies clusters of MNs that align with our proposed MN identities.** (**A**) Automatic prediction of synapse location allows us to compare the morphology of MNs by comparing the density of their input synapses in the T1 neuromere. We divided the neuropil into 8 µm x 8 µm x 8 µm voxels and counted the number of input synapses in each voxel for each MN. (**B**) The 69 MN vectors of synapse counts were normalized and then embedded from 1891-D voxel-space into two dimensions using the UMAP algorithm (McInnes et al., 2020). L2 normalization spread the clusters out from each other compared to L1 normalization. Most MN synapse density vectors fell into distinct clusters. Several MNs embedded near one another (inset). (**C**) The clusters identified by UMAP are aligned with the MN muscle targets. (**D**) An alternative metric to the UMAP embedding is the dot product of the synapse density vectors, called the cosine similarity. We used scikit-learn to compute the cosine similarity (Pedregosa et al., 2011). The matrix of pair-wise comparisons shows higher pair-wise similarity within the clusters identified by UMAP. (**E**) NBLAST is a standard method for computationally comparing neuron morphologies (Costa et al., 2016). The pairwise NBLAST scores are also higher within the UMAP clusters. (**F**) The cumulative density functions (CDFs) of cosine similarity comparisons within (black line) vs. across (gray line) clusters. (**G**) CDFs of NBLAST comparisons within (black line) vs. across (gray line) clusters. The area under the curve (AUC) is the Mann-Whitney U statistic divided by the product of numbers of elements. This measures the discriminability of within vs. across comparisons. The AUC is higher for the cosine similarity of the synapse density vectors than for NBLAST comparisons.

The UMAP clustering corroborates our assignments in our MN atlas above. The FANC MNs that we identified as belonging together are also identified as belonging together based on the locations of their synapses. This demonstrates that clustering neurons based on synaptic density is a potentially useful tool in conjunction with other clustering techniques like hierarchical clustering of a distance metric like cosine similarity or NBLAST. Hierarchical (Ward) clustering based on NBLAST or cosine similarity gives different clusters, particularly for smaller neurons (not shown). The cosine similarity matrix provides some intuition for why this is the case: MNs often have dendrites near MNs that target other muscles, such that the off-diagonal similarity can be high (**F**, gray curve). This is more pronounced for NBLAST (**G**, gray curve), since the method compares MN morphology, and MNs at least tend to have similar primary neurites.

UMAP clustering resolved many of the ambiguities that we encountered when trying to group MNs based on morphology. For example, cluster 4 contains all 11 neurons we identified as innervating the trochanter flexor muscle and accessory trochanter flexor muscle. In **Figure 3** in the main text, we show that MNs with two types of morphologies innervate the trochanter flexor muscle in the coxa. Cells with one morphology have anterior cell bodies and the other cells have posterior cell bodies. Both types congregate in cluster 4. For a second example, 9 of the 10 accessory tibia flexors separate into two clusters, cluster 7 (4 MNs) and cluster 8 (5 MNs), just as we observed in **Figure A13** and **A14**. Cluster 7 contains accessory tibia flexor MNs with more posterior primary neurites (**Figure A13F**), cluster 8 contains those with more anterior neurites.

UMAP clustering also revealed a novel phenomenon that we could not have predicted based on morphology alone, namely that the tarsus MNs that innervate muscles in the tibia do not cluster with one another, and instead separate into distinct clusters. Referencing the numbers in **Figure A17** above, FANC MN 65 clusters with two small ltm neurons. All three neurons express Dip-alpha, as previously described (Venkatasubramanian et al., 2019). FANC MN 66 clusters with the tibia flexor neurons and the four accessory tibia flexors in cluster 7. FANC MNs 67 and 68 cluster with the five accessory tibia flexors in cluster 8. FANC MN 69, which we believe is the tarsus levator MN, clusters with the tenth accessory tibia flexor MN (FANC MN 55) into cluster 9, which embeds near the FETi and the SETi in cluster 6. Finally, FANC MN 64 appears to be distinct. As described above, its unique morphology suggests it is the MN that targets what we term the retro tarsus depressor fibers that originate more distally to their insertion points. Together, this suggests that tarsus neurons may share more synapse locations with distinct clusters of accessory tibia flexor MNs. A companion paper analyses the input connections to these neurons and confirms that neurons that cluster together according to synapse density also receive common input from shared presynaptic partners (Lesser, Azevedo et al. 2023).

In summary, the UMAP embedding appears to cluster neurons that innervate shared portions of the neuropil. The most tantalizing interpretation of this result is that the T1 neuromere can be thought of as containing a distributed myotopic map, such that premotor neurons can target the MNs that control a particular joint by innervating locations that are common to that group of MNs.
